# Analog epigenetic memory revealed by targeted chromatin editing

**DOI:** 10.1101/2024.02.13.580200

**Authors:** Sebastian Palacios, Simone Bruno, Ron Weiss, Elia Salibi, Andrew Kane, Katherine Ilia, Domitilla Del Vecchio

## Abstract

Chemical modifications to histones and DNA play a crucial role in the regulation of transcription and in the maintenance of chromatin states that are not permissive to gene expression [1–3]. However, the landscape of gene expression states that these modifications stably maintain remains uncharted. Here, we show that gene expression can be memorized at a wide range of levels thus implementing analog epigenetic memory. Mechanistically, we find that DNA methylation serves a primary role in maintaining memory across cell divisions while histone modifications only follow DNA methylation to regulate gene expression. Employing targeted epigenetic editing and time-course analysis, we analyzed the temporal stability of gene expression and DNA methylation post removal of epigenetic effectors. We found that the grade of DNA methylation in the gene’s promoter, defined as the mean fraction of methylated CpGs, remains stable over time and inversely correlates with gene expression level. By contrast, Histone 3 lysine 9 trimethylation (H3K9me3) could not persist after removal of its writer in the absence of DNA methylation. These experimental findings, combined with our chromatin modification model, indicate that the absence of positive feedback mechanisms around DNA methylation - unlike those found in histone modifications - enable the temporal stability of the DNA methylation grade, which leads to analog memory. These results expand current knowledge on how epigenetic memory is achieved in natural systems. Moreover, we anticipate that analog memory through graded DNA methylation will enable to program mammalian cells with fine-grained information storage. This capability will significantly enhance the sophistication of engineered cell functionality in applications including tissue engineering, organoids, and cell therapies.

## Introduction

The ability to maintain multiple gene expression states through cell divisions and time is critical to a variety of biological functions, including the maintenance of distinct cell identities [4, 5], innate and adaptive immunity [6–8], and memory formation in the brain [9, 10]. Additionally, epigenetic memory of gene expression could be a pivotal tool for mammalian cell engineering [11]. Chromatin modifications, such as histone modifications and DNA methylation, are through to be critical mediators of epigenetic memory [1, 12–15], owing to their ability to affect gene expression by altering transcription [16–20] and because they self-propagate through DNA replication and cell division [16, 21, 22]. However, the complete spectrum of gene expression states that these chemical modifications stably maintain remains uncharted.

Heterochromatin is a well studied chromatin state in which expression is silenced and is maintained by concurrent DNA methylation and Histone H3 lysine 9 tri-methylation (H3K9me3) [21]. The temporal stability of this silenced state was assesed in prior studies by performing time-resolved analysis of reporter genes, which demonstrated that the silenced state is preserved long-term [23, 24]. Related studies have further shown that histone modifications mediate a bistable epigenetic switch where gene expression is either silenced or active [25, 26]. Therefore, current models of epigenetic memory posit that memory is *binary*, wherein genes are maintained either “on” or “off” based on autocatalytic histone modifications that induce bistability [27] (Fig. 1A). While informative, the above studies have been carried in a small set of contexts limited to artificial chromosomes or to a subset of endogenous genes. Because the effect of chromatin modifications on gene expression is highly context dependent [28–31], it remains unclear whether a model of binary memory generalizes across genes and cellular contexts, particularly for endogenous loci.

**Fig. 1.**
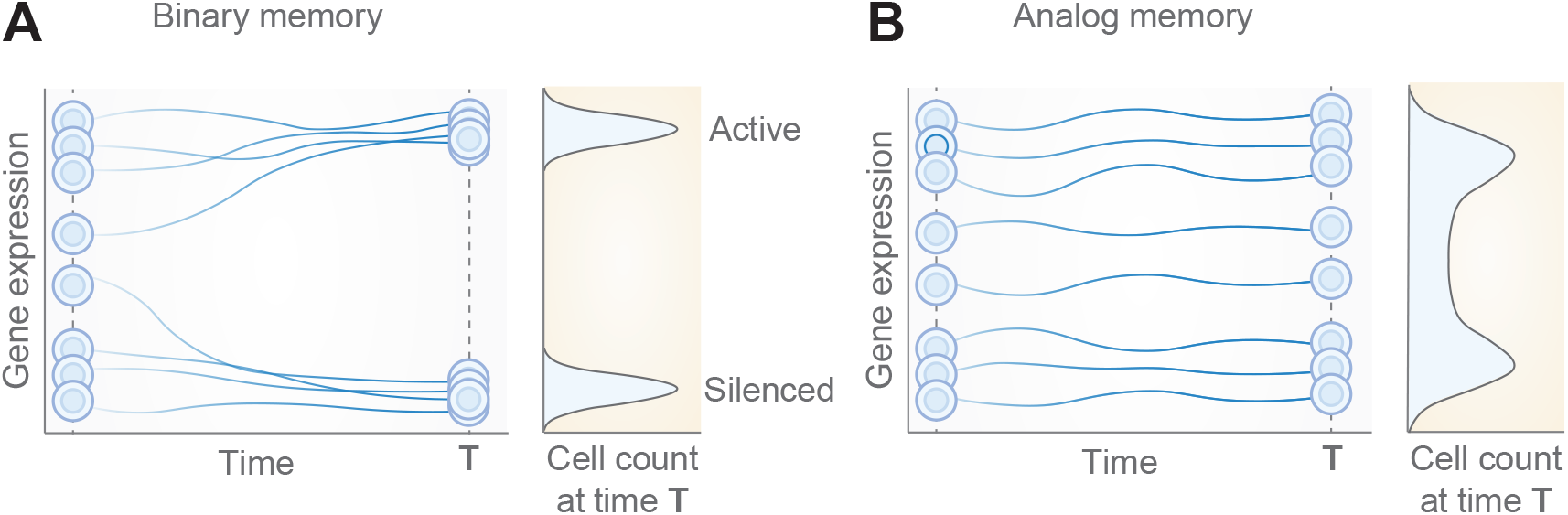
Binary versus analog epigenetic memory of gene expression. **A** A binary memory model posits that only silenced or active gene expression states are maintained long-term, but no other states. Each trajectory can be regarded as one cell. The emergent long-term distribution of gene expression across a cell population has two distinct peaks. **B** In an analog memory model, any gene expression level is maintained. The resulting long-term distribution of gene expression levels across a cell population can take any shape, including a bimodal shape with gene expression at all levels.

Beyond binary, additional forms of memory are possible: n-ary memory and analog memory. Memory that is n-ary extends the concept of binary memory to n*>* 2 instead of only two stably maintained states, and emerges from multi-stable systems, which have multiple, distinct, steady states that are attractive [32]. In the limit where the n stable states are next to each other to form a continuum of states, memory is *analog* (Fig. 1B). With analog memory, every possible state is thus stably maintained in time. None of these states is attractive since no temporal trajectory starting next to it will approach it as time progresses. Analog memory cannot thus emerge from multi-stability.

Elucidating whether additional forms of epigenetic memory exist and the chromatin modifications that enable them requires methods that initialize gene expression at various levels through targeted perturbations to the chromatin state, followed by time-resolved analysis of cell trajectories. Although methods using targeted editing of reporters knocked into specific endogenous genes allow for time-course analysis of cell trajectories, they can be subject to genetic context effects that are difficult to control and may influence the results [24, 29, 31, 33, 34]. To mitigate the influence of this context, prior studies have constructed a reporter gene within chromatin insulators in an artificial chromosome [23]; however, the extent to which these findings generalize to natural chromosomes remains unclear.

To overcome these difficulties, we developed a single-copy reporter gene integrated in a natural chromosome at a specific site separated from endogenous genes. We also constructed a set of engineered proteins for targeted editing of chromatin state and a chromatin modification model that, different from existing models, includes both histone and DNA methylation. We used these tools to map the spectrum of gene expression states that are memorized by chromatin modifications and determined the causal roles of histone modifications versus DNA methylation in long-term epigenetic memory. We show that memory is analog and that, despite recent emphasis on histone modifications, DNA methylation is the causal determinant of analog epigenetic memory with H3K9me3 only being an intermediate regulator of gene expression recruited by DNA methylation.

## Results

### An engineered system to study gene expression dynamics following targeted chromatin editing

We first set out to develop an experimental system to study epigenetic memory in a natural chromosome and the role that histone and DNA methylation play in its establishment and maintenance. To this end, we constructed a single-copy reporter gene integrated site-specifically in an endogenous mammalian locus and engineered a set of proteins for targeted epigenetic editing of the reporter gene (Fig. 2A, B; Fig. SE.1; Methods; Table SE1; Table SE2). The gene comprises the mammalian elongation factor 1a promoter (EF1a) driving the expression of fluorescent reporter (EBFP2). We flanked the gene with cHS4 chromatin insulators for isolation of the reporter from other genes [35]. The reporter comprises five binding sites upstream of the promoter for each of the DNA-binding proteins PhlF and rTetR (Fig. SE.1A, B; Table SE2) [36], where the binding of PhlF and rTetR to DNA can be modulated using 2,4-diacetylphloroglucinol (DAPG) and doxycycline (dox), respectively. Moreover, the region upstream of the promoter comprises five guide RNA (gRNA) target sites, which enable targeted epigenetic editing using epigenetic effectors fused to dCas9 (Fig. 2B; Fig. SE.1C; Table SE2; Table SE3). Epigenetic editing was performed by expressing epigenetic effectors fused to PhlF, rTetR, or dCas9 (Fig. SE.1A-F and Table SE2) using transient transfection. The epigenetic effectors were co-transfected with a fluorescent transfection marker (EYFP), which enabled the sampling of cells with varying levels of epigenetic effector expression using fluorescence-activated cell sorting (Fig. 2C).

**Fig. 2.**
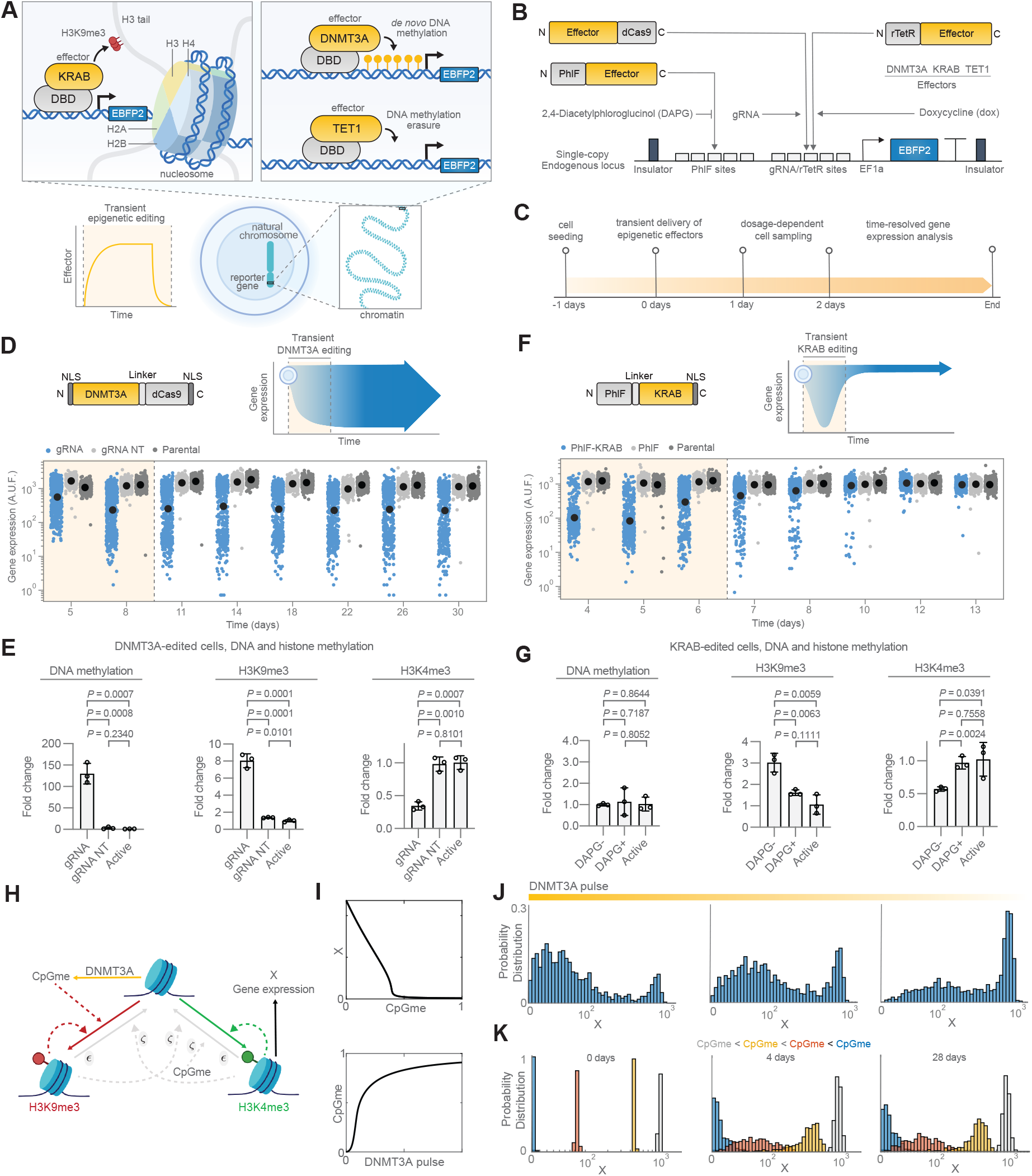
The gene expression dynamics in DNMT3A-edited cells indicate a form of memory different from binary. **A** Overview of transient epigenetic editing using KRAB, DNMT3A or TET1 fused to dCas9, PhlF or rTetR as DNA-binding domains (DBDs). **B** Schematic of the experimental system developed in this study. The reporter gene is integrated into an endogenous mammalian locus using site-specific chromosomal integration. A mammalian constitutive promoter (EF1a) drives the expression of fluorescent protein EBFP2. Upstream binding sites enable the targeted recruitment of epigenetic effectors, which are fused to DNA-binding proteins rTetR, PhlF or dCas9. The reporter gene is flanked by chromatin insulators for isolation from other genes. **C** Experimental overview describing transient transfection into cells bearing the reporter gene, fluorescence-activated cell sorting based on transfection level, and time-course flow cytometry measurements. **D** Gene expression dynamics of DNMT3A editing (DNMT3A-dCas9) of the reporter gene according to the experimental timeline shown in panel C. Shown are single-cell flow cytometry measurements (EBFP2) of DNMT3A-edited cells. DNMT3A-dCas9 was targeted to 5 target sites upstream of the promoter and a scrambled gRNA target sequence was used as control (Fig. SE.2A, B, Table S3). Shaded yellow denotes the time that the transfection marker was detected. Data shown are from a representative replicate from 3 independent replicates. **E** MeDIP-qPCR and ChIP-qPCR analysis 14 days after transfection of DNMT3A-dCas9 and cell sorting for high levels of transfection. The promoter region was analyzed (Table S4 and Methods). Data shown is from from three independent replicates. Reported are fold changes and their mean using the standard ΔΔ*C*_*t*_ method with respect to the active state. Error bars are s.d. of the mean. DNMT3A-dCas9 was targeted to 5 target sites upstream of the promoter (gRNA). A scrambled gRNA target sequence (gRNA NT) was used as control. \**P*<0.05, \*\**P*<0.01, \*\*\**P*<0.001, unpaired two-tailed t-test. **F** Gene expression dynamics of KRAB editing (PhlF-KRAB) according to the experimental timeline shown in panel C. Shown are flow cytometry measurements of the reporter gene (EBFP2) for single cells. The shaded yellow region indicates the time the transfection marker was detected during the time that DAPG was not applied. DAPG was applied in the PhlF-KRAB and PhlF conditions beginning at 6 days. A different independent replicate was measured on each day. Data shown are from 3 independent replicates. **G** MeDIP-qPCR and ChIP-qPCR analysis 6 days after transfection of PhlF-KRAB and cell sorting for high levels of transfection. The analysis is of the promoter region. Data is from three independent replicates. Shown are fold changes and their mean as determined by the standard ΔΔ*C*_*t*_ method with respect to the active state. Error bars are s.d. of the mean. \**P*<0.05, \*\**P*<0.01, \*\*\**P*<0.001, unpaired two-tailed t-test. **H** Chromatin modification circuit obtained when KRAB = 0, TET1 = 0. See SI Fig.SM.1C. **I** Top plot: dose-response curve for the (CpGme, X) pair. Bottom plot: dose-response curve between the intensity of a DNMT3A pulse and the DNA methylation grade (CpGme). The intensity of the pulse is increased by increasing its height. See SI Figs. SM.1D and SM.3. **J** Stationary probability distribution of gene expression for the system represented by reactions listed in SI Tables SM.1 and SM.4, with parameter values given in SI Section S.9.3. **K** Probability distributions of gene expression for the system after t = 28 days, obtained as described in panel J, with parameter values and initial conditions given in SI Section S.9.4. See SI Figs. SM.1B and SM.2. In panel I and J, the DNMT3A dynamics are modeled as a pulse that exponentially decreases over time (See Section S.1.1 - SI Equation (SM.7)). In our model, *ε* (*ζ*) is the parameter that scales the ratio between the basal (recruited) erasure rate and the auto-catalysis rate of each modification. See SI Fig. SM.1E and SM.3.

### Transient DNMT3A editing reveals non-binary memory of gene expression

We first investigated the gene expression dynamics following epigenetic editing by DNA methylation writer DNA methyltransferase 3 alpha (DNMT3A). To this end, we expressed a fusion of dCas9 and the catalytic domain of DNMT3A (DNMT3A-dCas9) [37] (Fig. SE.1 C; Table SE2), via transient transfection targeted to five target sites upstream of the promoter in the reporter gene using a gRNA (Fig. 2D; Fig. SE.2A, B; Table SE3). Subsequently, we utilized fluorescence-activated cell sorting to sample cell populations with various transfection levels and proceeded with time-course flow cytometry measurements of the resulting populations. We observed that gene expression was repressed, reaching maximal repression approximately 11 days after transfection (Fig. 2D and Fig. SE.2A). After reaching maximal repression, the cell distributions remained stationary for all levels of DNMT3A-dCas9 (Fig. 2D and Fig. SE.2A). This, along with no detection of the transfection marker of DNMT3A-dCas9 by day 8 (Fig. SE.2B), indicates memory of the transient DNMT3A editing. No repression was observed for cells where a gRNA with a scrambled target sequence was co-transfected with DNMT3A-dCas9 or in control experiments where dCas9 was targeted to the gene (Fig. SE.2C, D).

Although the epigenetic effector dosage increased the proportion of cells in the silenced state, all dosages of DNMT3A-dCas9 editing led to stationary distributions with cells expressing the reporter at all levels of gene expression ranging from silenced to fully active (Fig. 2D and Fig. SE.2A). Experiments where we transiently transfected the same DNMT3A catalytic domain fused to rTetR instead of dCas9 (rTetR-XTEN80-DNMT3A) (Fig. SE.1D; Fig.SE.3; Table SE2) also led to stationary distributions of gene expression spanning the whole spectrum of levels, from silenced to fully active. These observations ruled out binary memory (Fig. 1). Methylated DNA immunoprecipitation (MeDIP) and Chromatin immunoprecipitation (ChIP) followed by qPCR further showed an increase in DNA methylation and H3K9me3 levels, accompanied by a decrease in H3K4me3 level in the reporter gene compared to the untransfected (active) cells (Fig. 2E; Methods; Table SE4). This is in accordance with reports showing the ability of methylated DNA to recruit writers for H3K9me3 [38] and indicate that these two chromatin modifications play a role in the temporal maintenance of gene expression states.

### Transient KRAB editing leads to no permanent change of gene expression

In order to discriminate the roles of DNA methylation and H3K9me3 in the maintenance of gene expression states, we investigated the gene expression dynamics following epigenetic editing with writers of H3K9me3. To this end, we fused DNA-binding protein PhlF to KRAB (Fig. SE.1E and Table SE2) [39], an epigenetic effector that promotes H3K9 trimethylation [40, 41]. We thus expressed PhlF-KRAB via transient transfection in the cells bearing the reporter gene, sampled cell populations across different transfection levels using fluorescence-activated cell sorting, and performed time-course flow cytometry measurements (Fig. 2F and Fig. SE.4A-C). We observed rapid repression of gene expression for cells transfected with PhlF-KRAB and no repression for cells transfected with PhlF. Irrespective of the dosages of PhlF-KRAB, gene expression reverted to the active gene state within 4 days after reaching maximal repression. This aligns with previous reports where KRAB-based epigenetic editing resulted in temporary repression while failing to establish permament silencing [24, 37]. Further experiments with transient expression of PhlF-KRAB also resulted in temporary repression of the gene while no long-term memory was observed (Fig. SE.4D-F).

To determine what chromatin modifications emerged as a result of KRAB-mediated editing, we performed MeDIP and ChIP followed by qPCR on cells repressed with PhlF-KRAB (Fig. 2G and Methods). We observed an increase in H3K9me3 levels in the reporter gene compared to untransfected (active) cells, while DNA methylation levels remained unchanged and H3K4me3 levels decreased. Taken together, these data indicate that an increase in H3K9me3 in the absence of DNA methylation is unable to permanently downregulate gene expression. Therefore, DNA methylation but not H3K9me3 emerges as primary mediator of the temporal maintenance of gene expression states in cells edited by DNMT3A (Fig. 2D).

### A model where DNA methylation drives histone modifications predicts analog memory

Since *de novo* DNA methylation leads to H3K9me3 (Fig. 2E), we conclude that DNA methylation mediates the establishment of H3K9me3. This is consistent with studies reporting that methylated CpGs are bound by a reader protein (Methyl-CpG-binding Protein 2) that recruits H3K9me3 writers [38]. By contrast, H3K9me3 does not mediate the establishment of DNA methylation in our system (Fig. 2G). Taken together, these data suggest a model in which DNA methylation drives H3K9me3. Furthermore, the permanent change in gene expression resulting from transient recruitment of DNA methylation writer (Fig. 2D) but not from H3K9me3 writer (Fig. 2F), suggest that DNA methylation’s decay is negligible, while H3K9me3 decay is relatively fast. This long-term stability of DNA methylation has been attributed to the fast action of maintenance enzyme DNMT1, which re-writes DNA methylation on the daughter DNA strand upon DNA replication [16, 42].

We therefore propose a model in which the DNA methylation state remains constant in the absence of writer DNMT3A, drives the formation of H3K9me3 and the removal of H3K4me3 [14, 43, 44], consistent with our data (Fig. 2E). These histone marks, in turn, are known to mutually inhibit each other by recruiting each other’s erasers [45] and are autocatalytic since they each recruit their own writers [13, 21, 22, 46]. Furthermore, H3K4me3 but not H3K9me3 is associated with transcriptionally active promoters [22], leading to our chromatin modification model (Fig. 2H, see SI Table SM.3 for the chemical reaction system).

This model predicts an inverse graded relationship between the DNA methylation grade in the gene, defined as the mean fraction of methylated CpGs, and gene expression level (Fig. 2I (top), SI Section S.2). When the mutual inhibition between activating and repressive marks (*ζ*) is sufficiently strong, the map between the intensity of a DNMT3A pulse and the DNA methylation grade becomes ultrasensitive (Fig. 2I (bottom), and SI Fig. SM.1 D). This, in turn, with noise in the reactions and in the DNMT3A level leads to a bimodal distribution of gene expression in response to a transient DNMT3A input (Fig. 2J and SI Fig. SM.1E), which matches well experimental observations (Fig. SE.2A). The model also recapitulates the rest of the experimental observations (SI Fig. SM.5 and SI Tables SM.1 and SM.4 for the full chemical reaction system).

Despite the bimodal distribution of gene expression post recruitment and removal of DNMT3A, the model predicts that any initially set DNA methylation grade remains approximately constant in the absence of DNMT3A since the decay rate of DNA methylation is negligible. Therefore, any gene expression level initially set with the corresponding equilibrium DNA methylation grade persists in time with some variability under the effect of noise (Fig. 2K), demonstrating analog memory. Concerning noise properties, the model also predicts that intermediate DNA methylation grades will result in broader gene expression distributions (compare the orange and yellow distributions to the grey and blue distributions in Fig. 2K). The model explains this phenomenon when the basal erasure of histone modifications is sufficiently slow compared to their autocatalysis (small *ϵ*). In this regime, the histone modification circuit becomes bistable for intermediate DNA methylation grade (see SI Section S.4, SI Fig. SM.3A), which leads to increased gene expression variability under the influence of noise (SI Fig. SM.4). In summary, our data-educated model predicts that DNA methylation is the primary responsible for analog memory, with histone modifications following it to modulate gene expression level and its variability.

### Multiple gene expression levels are stably maintained and inversely correlate with DNA methylation grade

Following the model predictions, we hypothesized that epigenetic memory may be analog, thereby enabling cells to maintain any gene expression level. To test this hypothesis, we first sought to investigate whether cells with the gene in a state different from silenced and active within DNMT3A-edited populations exhibited long-term stability of gene expression. To this end, we performed fluorescence-activated cell sorting of the DNMT3A-edited cells from low, intermediate, and high gene expression levels (Fig. 3A, B; Methods). We then performed time-course flow cytometry measurements for 29 days to analyze the long-term gene expression trajectory (Fig. 3C-E). The cells with low and high levels of gene expression maintained an approximately stable distribution of gene expression levels after cell sorting, as expected by a binary memory model. Nonetheless, the cells with intermediate levels of expression, after an initial widening of the expression distribution, also kept a stable gene expression distribution across the population for the entire time-course. This is incompatible with binary memory (Fig. 1A). To characterize the epigenetic state of these cells, we performed ChIP followed by qPCR of the low, intermediate, and active cell populations (Fig. 3F and Methods). We found an increase of H3K9me3 in the low population when compared to the intermediate and active cell populations. Accordingly, we observed a decrease of H3K4me3 in the low and intermediate expressing cells when compared to the active cells. These results are consistent with the known role of H3K9me3 as a chromatin modification associated with repression and H3K4me3 as a chromatin modification associated with active transcription at promoters [47].

**Fig. 3.**
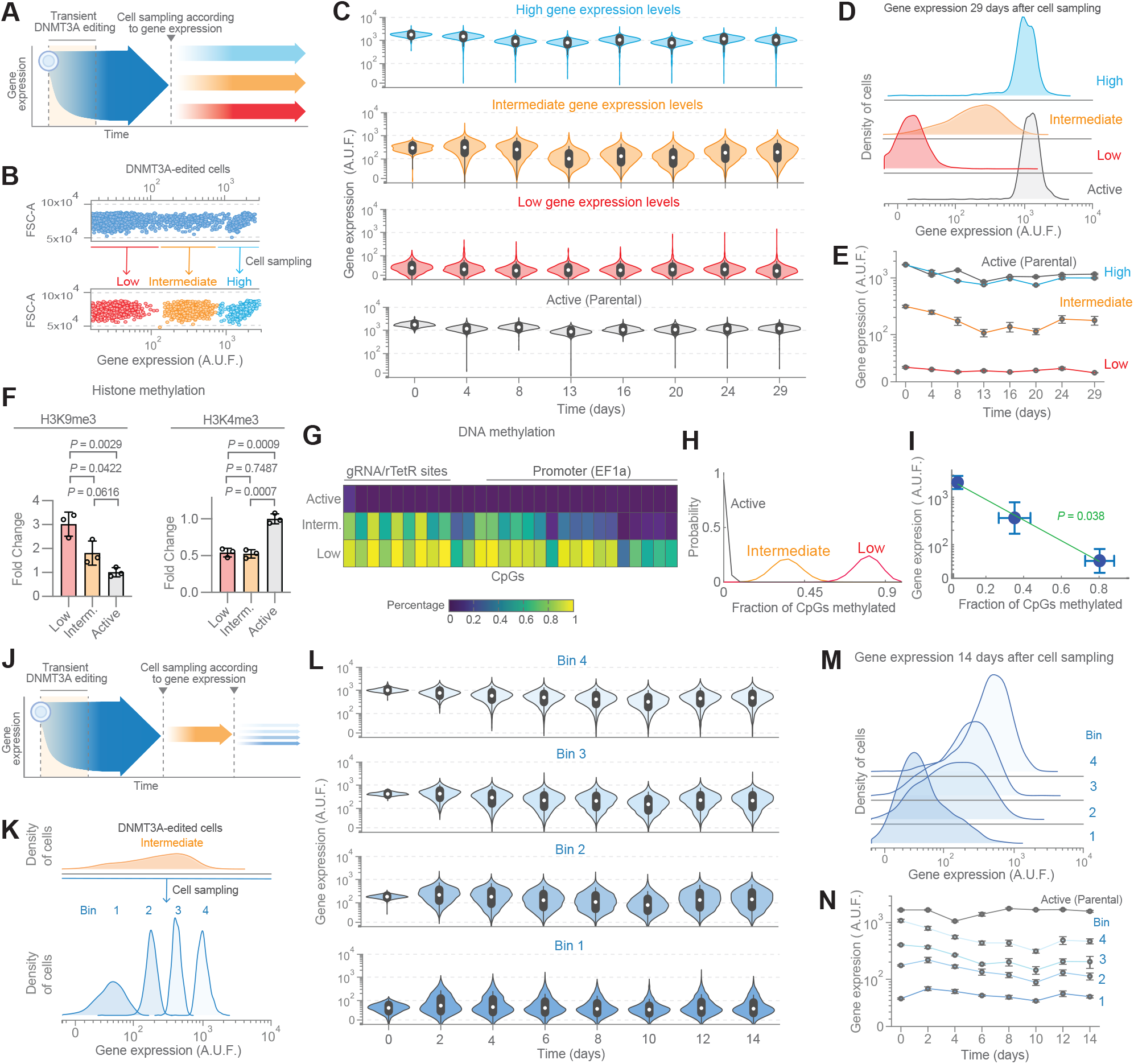
Gene expression is be stably maintained at multiple levels in DNMT3A-edited cells. **A** Conceptual overview of panels B-E. **B** Fluorescence-activated cell sorting of DNMT3A-edited cells (DNMT3A-dCas9) according to low, intermediate, and high levels of gene expression (EBFP2). Shown are single-cell gene expression measurements on the day of cell sorting using flow cytometry. Data are from a representative replicate from three independent replicates. **C** Time-course flow cytometry analysis of the dynamics of gene expression exhibited by cells obtained in panel B. Data are from one representative replicate from three independent replicates. **D** Density of cells from flow cytometry measurements obtained on the last time point in panel C (29 days after cell sorting). Data shown are from one representative replicate from three independent replicates. **E** Dynamics of gene expression exhibited by cells shown in panels B-D. Shown is the mean of geometric means from three independent replicates. Error bars are s.d. of the mean. **F** ChIP-qPCR for H3K9me3 and H3K4me3 of the promoter in the reporter gene (untransfected cells bearing the reporter gene). Shown are fold changes and corresponding mean from three independent replicates as determined by the ΔΔ*C*_*t*_ method with respect to the active state. Error bars are s.d. \**P*<0.05, \*\**P*<0.01, \*\*\**P*<0.001, unpaired two-tailed t-test. **G** Heatmap of targeted bisulfite sequencing of the reporter gene. Shown is the mean percentage of DNA methylation for each CpG from three independent replicates. **H** Probability distribution of the fraction of CpGs methylated (Methods). **I** Correlation plot between the mean fraction of CpGs methylated and mean gene expression level. The green line represents the best-fit line *y* = −0.5059*x* + 0.5497 obtained by linear regression. The p-value associated with the slope coefficient is p = 0.038 *<* 0.05 (Methods). **J** Conceptual overview for panels K-N. **K** Fluorescence-activated cell subsorting of DNMT3A-edited cells (DNMT3A-dCas9) exhibiting intermediate levels of gene expression (EBFP2). Data is from gene expression measurements using flow cytometry on the day of cell sorting. Data is from a representative replicate from three independent replicates. **L** Flow cytometry-based time-course analysis of the dynamics of gene expression exhibited by cells obtained in panel K. Data is from one representative replicate from three independent replicates. **M** Density of cells from flow cytometry measurements obtained on the last time point in panel L (14 days after cell sampling). Data shown are from one representative replicate from three independent replicates. **N** Dynamics of gene expression exhibited by cells shown in panels K-N. Shown is the mean of geometric means from three independent replicates. Error bars are s.d. of the mean.

Because we pinpointed DNA methylation as the critical mediator of the maintenance of gene expression states, we performed targeted bisulfite sequencing of the reporter gene for each of the three cell populations: low, intermediate, and active (Fig. 3G, Methods). While the active state is devoid of methylated CpGs, the probability of finding any given CpG methylated in the promoter and rTetR binding sites in the intermediate state is distinctly smaller than in the low state and higher than in the active state. From these data, we computed the probability distributions of the fraction of methylated CpGs in the promoter and rTetR binding sites, for each of the low, intermediate, and active states (Fig. 3H and Methods). These distributions reinforce that the intermediately expressing cells have intermediate levels of DNA methylation. In particular, the mean value of the fraction of methylated CpGs, which we define here as the DNA methylation grade, is intermediate in cells with intermediate expression levels compared to that of the low and active cells and has a loglinear negative correlation (p-value <0.05) with the mean gene expression level (Fig. 3I). All these findings are recapitulated by our model (see SI Fig. SM.6B-E).

We additionally tested the hypothesis that epigenetic memory can operate in an analog manner by further subdividing DNMT3A-edited cells according to gene expression level, followed by time-resolved tracking (Fig. 3J-N; Methods). If epigenetic memory were analog, cells sampled from different gene expression levels should approximately maintain these distinct levels over time. Each of the four resulting cell populations maintained distinct distributions of gene expression levels according to the level on the day of cell sampling (Fig. 3K-N). We also sampled cells that had been edited with DNMT3A fused to rTetR (rTetR-XTEN80-DNMT3A) instead of dCas9 using fluorescence-activated cell sorting (Fig. SE.5A-E; Methods), and the same temporal stability was observed. These observations support the hypothesis that the memory is analog (Fig. 1B) in agreement with the model (Fig. 2K and SI Fig. SM.6F-H), and that DNA methylation grade is the primary mediator of the stable maintenance of gene expression levels.

### A continuum of gene expression levels are memorized and inversely correlate with a stable DNA methylation grade

To further test the hypothesis that epigenetic memory operates on an analog basis, we set out to characterize the long-term trajectories of gene expression with extensive sampling of single cells from DNMT3A-edited cell populations followed by monoclonal expansion and analysis (Fig. 4). This approach provides a high-resolution evaluation of gene expression trajectories generated exclusively from one initial DNA methylation profile. We first performed highly dense sampling of single cells, using fluorescence-activated cell sorting, from three distinct cell populations that had been previously sampled from DNMT3A-edited cells (Fig. 4A, B and Methods). Specifically, we selected eight different consecutive regions of gene expression levels spanning across low (2 regions), intermediate (4 regions), and high (2 regions) gene expression levels in DNMT3A-edited cells (Methods). After sampling single cells from each of those regions, we performed clonal expansion, resulting in ≥10 monoclonal populations from each of the eight gene expression regions for a total of 93 monoclonal populations. Flow cytometry measurements were performed for each of the cell populations after 15 days of monoclonal expansion (Fig. 4C). The monoclonal populations showed a continuum of gene expression levels ranging from background expression level to active gene expression level. To test whether the gene expression levels from the monoclonal populations could be maintained in time, we selected 8 monoclonal populations (highlighted in Fig. 4C), spanning the gene expression range from the 93 clones, and performed time-course flow cytometry measurements up to 161 days (> 5months) since the beginning of monoclonal expansion (Fig. 4D). In these measurements, we observed that the gene expression distributions remained approximately stable over time, validating the initial hypothesis that cells can maintain memory of any gene expression level, which defines analog memory.

**Fig. 4.**
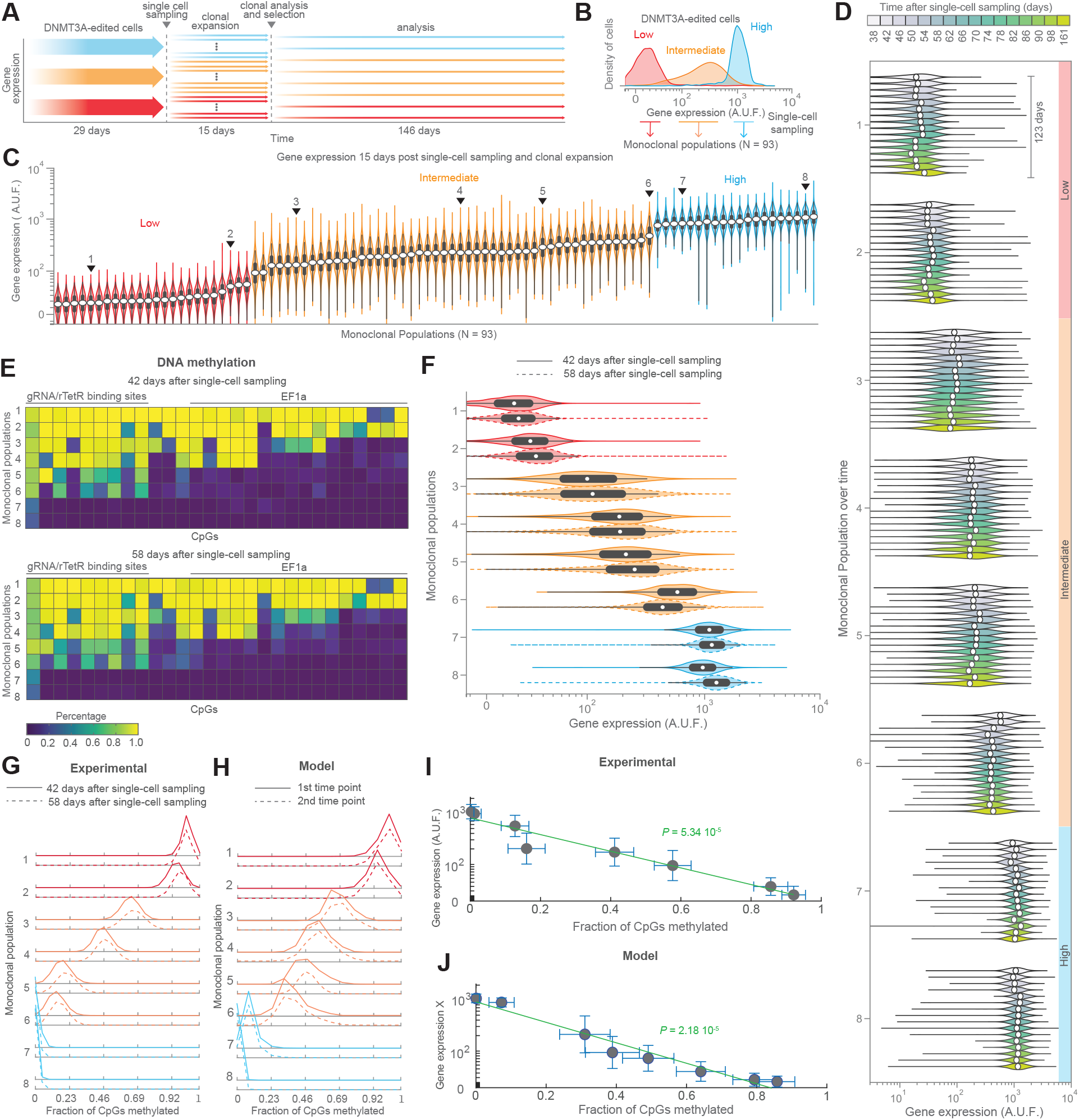
The memory is analog and is mediated by DNA methylation. **A** Conceptual experimental overview. **B** Single cells were obtained after sampling of DNMT3A-edited cells with low, intermediate and high levels of gene expression. After monoclonal expansion, the total number of monoclonal populations obtained was N = 93 (Methods). **C** Violin plots of the monoclonal populations (N = 93) measured 15 days after single-cell sampling from the cell populations shown on panel B. Monoclonal populations were positioned on the x-axis according to geometric mean of gene expression. Clones indicated with numbers 1-8 were selected for further analysis. Colors represent the cell population of origin as shown in panel B. **D** Flow cytometry-based time course analysis of gene expression dynamics of the eight monoclonal populations selected in panel C. **E** Bisulfite sequencing of the reporter gene in the eight monoclonal population at two different time points. **F** Flow cytometry measurements of the reporter gene in the eight monoclonal population at two different time points corresponding to those shown on panel E. **G** Probability distribution of the fraction of CpGs methylated (See “Methods” section for details on deriving these plots from data in panel E). The standard deviation for each clone is the following: 0.1752, 0.4707, 0.9492, 1.2714, 1.3101, 1.262, 1.1683, and 0.8095, ordered from clone 1 to clone 8. **H** Simulations of the probability distributions of the system represented by the reactions listed in SI Tables SM.6 and SM.4. The distributions are obtained computationally using SSA [52]. The standard deviation for each clone is: 0.1346, 0.5472, 1.2585, 1.1189, 1.0431, 1.1117, 0.8512, and 0.8259, ordered from clone 1 to clone 8. The parameter values used for these simulations are listed in SI Section S.9. **I** Correlation plot between the mean fraction of CpGs methylated and mean gene expression level at 42 days after single-cell sort. The green line represents the best linear fit *y* = −0.3756*x* + 0.4358. The p-value associated with the slope coefficient is *P* = 5.34 · 10^−5^ *<* 0.05 (See “Methods” section for details on the linear regression). **J** Correlation plot between the mean fraction of CpGs methylated and mean gene expression level, X, obtained via simulations. The green line represents the best linear fit *y* = −0.4673*x* + 0.4580. The p-value associated with the slope coefficient is *P* = 2.18 · 10^−5^ *<* 0.05 (See “Methods” section for details on the linear regression).

We next investigated DNA methylation grade as a determinant of the maintenance of gene expression level. To this end, we performed targeted bisulfite sequencing of the reporter gene in each of the eight populations at two different time points (Fig. 4E,F; Methods). The data showed that clones (7 and 8) selected from cells in the high gene expression distribution (Fig. 3B) are largely devoid of methylated CpGs. By contrast, clones (1 and 2) selected from cells within the low distribution have the majority of CpGs methylayed. The remaining clones (3, 4, 5, and 6), selected from the intermediate distribution, display a graded decreasing number of CpGs that are methylated with high probability (Fig. 4E). These trends inversely correlate with the gene expression levels in the clones (Fig. 4F). To acquire a quantitative understanding of the graded nature of this variation, we computed from the bisulfite sequencing data the probability distribution of the fraction of methylated CpGs in the gene’s promoter and rTetR binding sites (Methods). This confirmed a progressive and gradual shift of the distributions through the clonal populations (Fig. 4G), in accordance with model predictions (Fig. 4H). The correlation between the mean gene expression level of each clonal population and the mean number of methylated CpGs is loglinear (Fig. 4I,J), confirming a gradual increase of the DNA methylation grade through the clonal populations from high to low gene expression levels. Furthermore, the distribution of the fraction of methylated CpGs remained approximately unchanged between the two time points where bisulfite sequencing was performed (Fig. 4G), indicating that the DNA methylation grade is stably maintained.

Further analysis of the clonal distributions of the fraction of methylated CpGs revealed that the variability of the number of methylated CpGs about their mean value, measured by the standard deviation, is larger for intermediate DNA methylation grades when compared to low and high methylation grades (Fig. 4G). The mathematical model recapitulates this property (Fig. 4H) when it includes the two following additional possible molecular interactions: (a) H3K9me3 helps recruit DNA methylation maintenance enzyme (DNMT1) to the gene [48] and (b) DNMT1 can cause *de novo* DNA methylation, although at very low rate [49, 50] (see SI Section S.5 for model derivation). With these interactions, the errors made by DNMT1 in maintaining methylation are less probable for a fully methylated gene due to higher recruitment rate of DNMT1 via H3K9me3. Similarly, errors made by DNMT1 in *de novo* methylating additional CpGs are less probable for a gene with no methylation since DNMT1 recruitment is practically absent (see SI Section S.5 and Fig. SM.7 for model analysis.) Overall, these interactions can explain the tighter distributions observed for highly and lowly methylated gene states (Fig. 4G).

The genome engineering system that we utilized in this study has been demonstrated to be highly precise [51]. Nevertheless, we still sought to verify that the different clones all had one genomic copy of the reporter gene to rule out the possibility that the variation of gene expression observed across the clones could be due to variability in the genomic integration copy number. To this end, we performed copy number variation analysis using digital PCR (dPCR), which ruled out this possibility (SE.5F and Methods). Taken together, these data establish that the DNA methylation grade is maintained by the clonal populations and that this grade fine-tunes the gene expression level.

### Transient editing of DNA methylation permanently impairs gene expression level maintenance

Next, we sought to establish a causal link between the DNA methylation grade and the level of gene expression. To this end, we investigated whether altering the DNA methylation grade in cells maintaining an intermediate level of gene expression would impair this maintenance. To enhance DNA methylation, we utilized rTetR-XTEN80-DNMT3A as we have shown that DNMT3A enhances DNA methylation (Fig. 2D) in accordance with prior reports [37]. To remove DNA methylation, we engineered a fusion protein comprising rTetR fused to the catalytic domain of the ten eleven translocation protein 1 (TET1) [34, 53] to construct rTetR-XTEN80-TET1 (Fig. SE.1F and Table SE2). This catalytic domain has been shown to enable targeted DNA demethylation of mammalian genes [34]. Moreover, we fused rTeR to KRAB to construct rTetR-KRAB (SE.1G and Table SE2) in order to evaluate changes to gene expression when enhancing H3K9me3 without changing DNA methylation. We transiently transfected these epigenetic effectors in cells that had been sampled from intermediate levels of gene expression in DNMT3A-edited cells. As before, we utilized fluorescence-activated cell sorting to obtain transfected cells and subsequently performed time-course flow cytometry measurements (Fig. 5 and Fig. SE.5G-I). By 10 days after transient transfection with rTetR-XTEN80-DNMT3A, the gene expression was repressed, remained silenced over time, and overlapped with the distribution of cells that had been previously sorted for low levels of gene expression post DNMT3A-treatment (Fig. 5A). Transient transfection with rTetR-XTEN80-TET1 resulted in nearly complete and permanent reactivation of gene expression (Fig. 5B). By contrast, transient transfection with rTetR-KRAB did not result in any permanent change of gene expression. Specifically, 6 days after transfection, we observed levels of repression comparable to those of the low levels in DNMT3A-edited cells, but then expression reverted to the initial intermediate level and remained stable afterwards (Fig. 5C). These data indicate that transient recruitment of a DNA methylation writer or eraser to an intermediadely methylated, and partially active gene, can permanently silence or reactivate the gene. By contrast, transient recruitment of an H3K9me3 writer does not exert a permanent change in gene expression. Taken together, these data show that DNA methylation grade is the causal determinant of the maintenance of gene expression levels in accordance with the model predictions (Fig. SM.8).

**Fig. 5.**
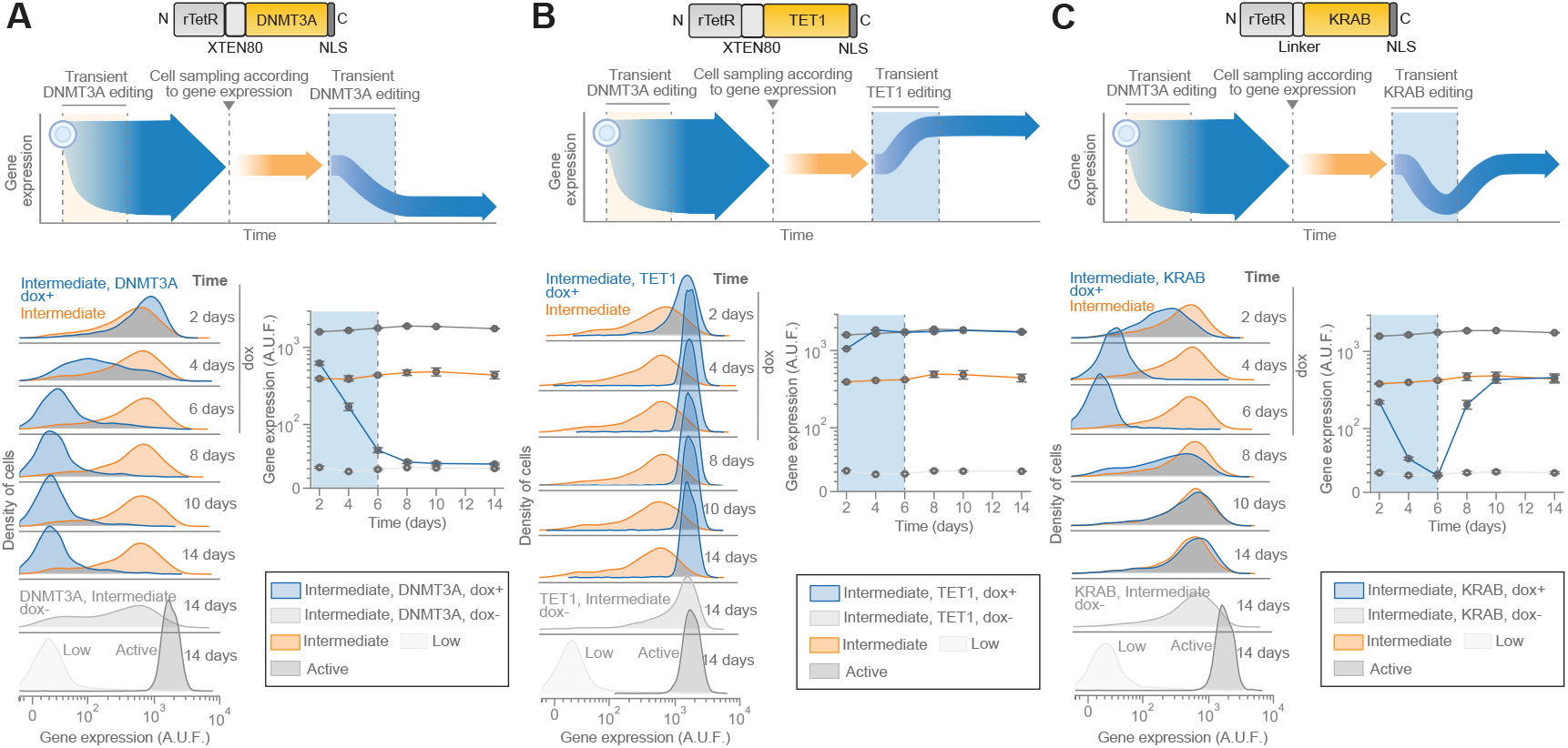
DNA methylation effectors (DNMT3A or TET1) impair the maintenance of intermediate gene expression states in DNMT3A-edited cells. Time-course of flow cytometry measurements after transient editing using **A** rTetR-XTEN80-DNMT3A **B** rTetR-XTEN80-TET1 or **C** rTetR-KRAB in DNMT3A-edited cells with stable intermediate levels of gene expression. Densities of cells shown are from one representative replicate from three independent replicates. Gene expression is from the reporter gene (EBFP2). Shaded blue region indicates the time the transfection marker was detected during the time that doxycycline was added. Doxycycline was applied for 6 days. Shown is also the mean of geometric means from three independent replicates. Error bars are s.d. of the mean.

## Discussion

Current models of gene regulation by chromatin modifications posit that epigenetic memory is binary, largely based on autocatalytic histone modifications that lead to bistability where gene expression is maintained “on” or “off” [27]. Although some experimental studies support this hypothesis [23, 25, 26], the context-dependent nature of the effect of chromatin modifications on gene expression [28–31] has left doubts on whether a binary memory model extends beyond the specific experimental systems considered and whether other forms of memory exist. To address this question, we have developed a synthetic reporter gene in mammalian cells and a set of sequence-specific chromatin regulators, which allow to study the temporal dynamics of gene expression following targeted perturbations to the chromatin state. With these tools, we have discovered that gene expression can be memorized at a continuum of levels and that DNA methylation enables this analog memory. By contrast, histone modifications only follow DNA methylation to modulate gene expression.

Despite the growing emphasis on histone modifications as regulators of chromatin state [28, 47], these modifications do not operate in isolation but in orchestration with DNA methylation [38, 54–57], albeit in a highly context-dependent manner [2]. In principle, DNA methylation can recruit writers of H3K9me3 [38], consistent with our ChIP and MeDIP data (Fig. 2E), and H3K9me3 can recruit writers of DNA methylation [57, 58], which instead is not the case in our study (Fig. 2H) nor in similar studies [23]. In turn, it has been reported that H3K4me3 and DNA methylation inhibit each other [55, 56], consistent with our data (Fig. 2E). Our chromatin modification model emerges from this evidence and thus includes all the named interactions except for the establishment of DNA methylation mediated by H3K9me3 (Fig. 2G).

In this model, unlike histone modifications, DNA methylation is not under positive feedback (Fig. 6A). Coupled with practically absent decay rate, this feature makes any DNA methylation grade persist in time long-term. Since DNA methylation grade also modulates gene expression by inhibiting H3K4me3 [22, 56], it confers gene expression levels long-term memory. By contrast, histone modifications H3K9me3 and H3K4me3 alone, when under strong positive feedback and mutual inhibition, collapse to a state of either full H3K9me3 (gene repression) or full H3K4me3 (gene activation) (SI Section S.4). In our model, the DNA methylation grade biases the balance between H3K4me3 and H3K9me3, thereby modulating through these marks the gene expression distribution. This distribution, in turn, spreads depending on the DNA methylation grade, which influences the bistability of the histone modification circuit (SI Figs. SM.3A,B and SM.4).

**Fig. 6.**
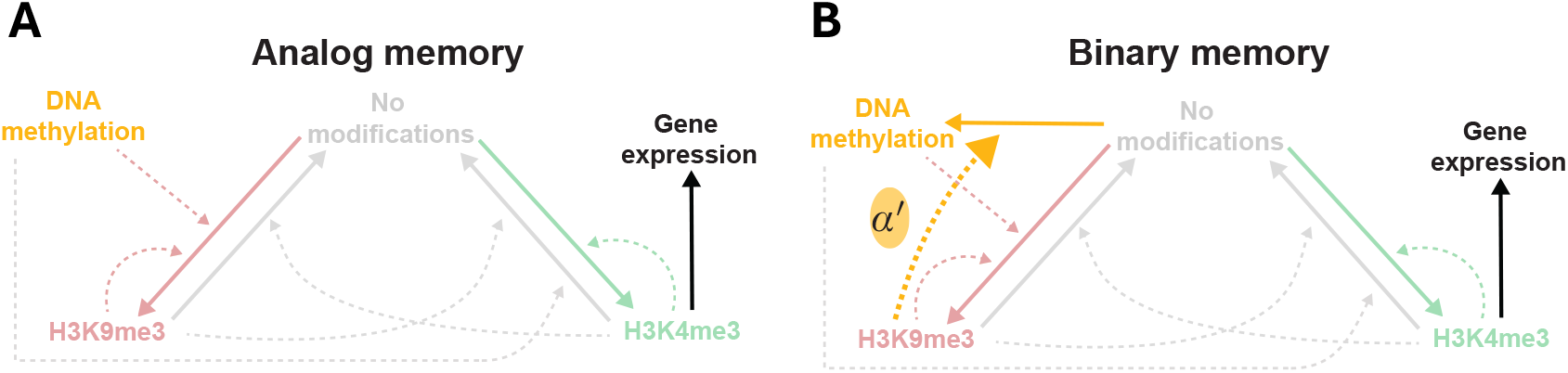
The interactions between DNA methylation and histone modifications explain when analog verus binary epigenetic memory emerges. **A** Analog memory. DNA methylation is not under positive feedback because the loop between DNA methylation and H3K9me3 is broken. **B** Binary memory. DNA methylation is under positive feedback because H3K9me3 recruits writer enzymes for DNA methylation (yellow dashed arrow marked by *α*_′_) and DNA methylation mediates the recruitment of H3K9me3. In the diagrams, we have assumed that no external epigenetic effectors (such as KRAB, DNMT3A, and TET1) are present. Refer to Fig. SM.1A, C for the complete diagram and model.

Because the crosstalk between DNA methylation and histone modifications is context-dependent [2], it is plausible that different patterns of such interactions appear in different cellular and genetic contexts. When H3K9me3 mediates the establishment of DNA methylation, in addition to the reverse, also DNA methylation is under positive feedback (Fig. 6B and SI Fig.SM.1). This can push an initially partially methylated gene to be fully methylated. The competitive interaction of DNA methylation with H3K4me3, in turn, can cause H3K4me3 to win over an initially low DNA methylation state and to revert this to the unmethylated state. When this positive feedback is sufficiently strong, then binary memory emerges (SI Section S.3, Fig. SM.2), and fully methylated and completely unmethylated states are the only states that are inherited. In practice, we expect that the specific cellular and genetic context will determine the strength by which H3K9me3 mediates the establishment of DNA methylation (parameter *α*′, Fig 6B), and hence the temporal duration of analog states of gene expression (Fig. SM.2). This context-dependence may explain why the expression of fate specific genes is binary in certain cell types while it appears continuous in others [59, 60] and is associated with graded DNA methylation [61].

More broadly, analog epigenetic memory may provide a new angle to understand the role of DNA methylation in important biological processes, such as for memory formation in the brain [9, 10] and regulation of immune response in the immune system [6–8]. However, more research will be required to determine the extent to which naturally occurring gradients of gene expression correlate with DNA methylation grade and are temporally maintained at the single cell and gene level. Although in principle single-cell nucleosome, methylome, and transcriptome sequencing (scNMT-seq) [62] can determine the correlation between DNA methylation and gene expression [63], temporal data of the expression of single genes in single cells will be necessary to investigate persistence of DNA methylation grade and gene expression levels through cell divisions and time. Furthermore, an understanding of how analog memory is realized can become a formidable tool for engineering mammalian cells with analog information storage capabilities. Programming analog memories through DNA methylation grade could allow the engineering of gradients of cell identities, thereby enhancing the sophistication and functionality of tissues and organoids that we can create. To this end, additional research will be needed in order to re-engineer the chromatin modification circuit such that we can precisely control DNA methylation grade by user-defined inputs, such as small signaling molecules and drugs.

In summary, our study has revealed analog memory of gene expression and identified DNA methylation as the primary mediator. Our model further suggests how genetic and cellular context influences the emergence of binary or analog memory. Finally, our epigenetic effectors and reporter system can serve as a tool to explore the context-dependent nature of epigenetic memory across genes and cellular processes and as a starting point to engineer user-defined programming of cellular memories.

## Methods

### Cell culture conditions

Cells were maintained in tissue culture plates (Corning) with growth media comprised of Dulbecco’s Modified Eagle Medium (DMEM) (Corning) supplemented with 10% FBS (Corning), 1X non-essential amino acids (Gibco), 100 U/ml penicillin (Sigma), and 100 *µ*g/ml streptomycin (Sigma) at 37°C in a 5% CO2 incubator. The growth media was changed every 2-4 days. For routine passaging, the growth media was removed, followed by the addition of 0.25% trypsin (Corning) and incubation at 37°C in a 5% CO2 incubator for 7 minutes. Trypsin was then neutralized with growth media at room temperature. The cells were then centrifuged for 3 minutes at 200g, the supernatant was discarded, and the cell pellet was resuspended in fresh growth media. The cells were then diluted (ratio between 1 and 20) and reseeded into new tissue culture plates for subsequent cell culture.

### Transfections

Cells were cultured in growth media at 37°C in a 5% CO2 incubator. The cells were then treated with 0.25% trypsin (Corning) and incubated for 7 minutes at 37°C in a 5% CO2 incubator. The cell suspension was subsequently neutralized with growth media at room temperature. Cells were centrifuged for 3 min at 200g followed by removal of the supernatant and cell resuspension in fresh growth media. Cells were then seeded in 12-well tissue culture plates (Corning) at 300,000 cells/well and cultured for 24 hours at 37°C in a 5% CO2 incubator. Growth media was replaced with growth media without antibiotics. Transfection was then performed using Lipofectamine LTX (Invitrogen) according to the manufacturer’s instructions. After 24 hours of incubation at 37°C in a 5% CO2 incubator, replacement for routine growth media supplemented with penicillin and streptomycin was performed. All transfections with rTetR-XTEN80-DNMT3A, rTetR-XTEN80-DNMT3A, rTetR-XTEN80-TET1 and rTetR-KRAB were performed using 500ng of the fusion protein and 300ng of trasfection marker (EYFP). Transfections with PhlF-KRAB that involved cell sorting were performed using 450ng of either PhlF-KRAB or PhlF, and 450ng of the transfection marker (450ng). Experiments of PhlF-KRAB without cell sorting were perfomed using 450ng of PhlF-KRAB and 450ng of the transfection marker (1:1 stochiometry), 45ng of PhlF-KRAB and 450ng of the transfection marker (0.1:1 stochiometry), or 4.5ng of PhlF-KRAB and 450ng of the transfection marker (0.01:1 stochiometry). An empty vector was used to maintain the DNA amounts constant for the various stoichiometries. Transfections of DNMT3A-dCas9 that involved cell sorting were performed using 500ng of DNMT3A-dCas9, 600ng of the gRNA expression vector, and 300ng of the trasfection marker (EYFP). In experiments using dCas9, 450ng of the gRNA expression vector, 300ng of either DNMT3A-dCas9 or dCas9, and 300ng of the transfection marker (EYFP) were used. All the experiments presented use 2*µ*M of dox and 30*µ*M of DAPG.

### Chromosomal integration

The reporter gene was integrated using site-specific chromosomal integration into an endogenous mammalian locus (Rosa26) in CHO-K1 cells bearing a landing pad (monoallelic) for efficient and precise integration [51]. This locus had been previously identified by a homology-based computational search (BLAST) using the murine Rosa26 locus as reference. EYFP is expressed from the landing pad exclusively before site-specific chromosomal integration into the Rosa26 locus. The landing pad confers puromycin resistance exclusively after site-specific chromosomal integration into the Rosa26 locus. For integration, 500ng of the chromosomal integration vector and 500ng of the Bxb1 recombinase expression vector were transfected into the cells. After 24 hours, routine cell culture was commenced with the supplementation of growth media with puromycin (8*µ*g/ml) for 2 weeks. Subsequently, a fluorescence-activated cell sorter (BD Aria) was utilized to obtain cells that were positive for EBFP2 expression and negative for EYFP expression. Digital PCR was used to confirm 1 reporter gene per genome (Fig. SE.5F; Methods). The cells were also assayed and confirmed to test negative for mycoplasma contamination (MIT’s Koch Institute Preclinical Modeling Core).

### DNA construction

DNA cloning was performed using a combination of Golden Gate Assembly, Gibson Assembly, restriction cloning, and PCR cloning. Expression vectors for the epigenetic effectors were cloned using Golden Gate Assembly. The cloning was performed in a single reaction with the plasmids comprising a destination vector, cHS4 chromatin insulator, EF1a promoter, 5’ UTR, gene sequence, 3’ UTR, and polyadenylation signal. Gene sequences for rTetR-TET1, rTetR-XTEN80-TET1, rTeTR-DNMT3A, rTeTR-XTEN80-DNMT3A, PhlF-KRAB, rTetR-KRAB, and PhlF were synthesized and cloned into a plasmid with compatible overhangs for Golden Gate Assembly (Azenta Life Sciences). The expression vectors for TET1 and rTetR were cloned by performing PCR cloning using the rTetR-XTEN80-TET1 expression vector as a template. For the reporter gene, the region comprising the PhlF and rTetR binding sites was synthesized (Azenta Life Sciences) and subsequently cloned upstream of a vector containing the EF1a promoter using restriction cloning. Golden Gate Assembly was then performed in a reaction comprising the EF1a promoter with upstream DNA-binding sites, destination vector, upstream cHS4 chromatin insulator, 5’ UTR, EBFP2, 3’ UTR, and polyadenylation signal. The resulting vector was cloned along with the downstream cHS4 chromatin insulator into the chromosomal integration vector using Gibson Assembly. All sequences were sequenced verified via Sanger sequencing (Azenta Life Sciences) or whole plasmid sequencing (Primordium).

### Flow cytometry

Cells were trypsinized using 0.25% trypsin (Corning) and incubation at 37°C in a 5% CO2 incubator for 7 minutes. Following this, trypsin was neutralized by adding growth media at room temperature. Afterwards, the cells were centrifuged at 200g for 3 minutes. The supernatant was removed, and the resulting cell pellet was then resuspended in fresh growth media. The cell suspension was filtered through a filter cap into flow cytometry tubes (Falcon). Flow cytometry data was then collected using a flow cytometer (BD LSRFortessa). EBFP2 was measured using a 405nm laser and a 450/50 filter with 259 V, EYFP was measured using a 488nm laser and 530/30 filter with 190 V, mKO2 was measured using a 561nm laser and 610/20 filter with 250 V, iRFP was measured using a 637nm laser and 670/30 filter with 326 V. Cytoflow (version 1.2) was used for analyzing all flow cytometry data and obtaining cell densities, means, standard deviations, and percentages of flow cytometry measurements. For analysis, measurements were gated for single cells based on forward and side scatter (Fig. SE.5J), and only gated measurements were used for subsequent analysis. After processing the flow cytometry data using Cytoflow, the data was plotted using either Cytoflow, GraphPad Prism or Matplotlib. Graphpad Prism was used for generating bar plots. For the correlation plot between mean EBFP2 fluorescence and DNA methylation, Cytoflow’s gated measurements were imported into MATLAB, where the correlation was performed using the polyfit function.

### Time-course flow cytometry

A fluorescence-activated cell sorter (BD Aria) was used to obtain cells with specific levels of transfection (EYFP) or gene expression (EBFP2). For transfection experiments that involved cell sorting, cell sorting was performed between 42-48 hours after transfection. When flow cytometry was performed on the day of cell sorting, a fraction of the cells that were sorted was utilized for flow cytometry measurements (BD LSRFortessa), and the rest of the cells were reseeded on tissue culture plates. For flow cytometry measurements, cells were trypsinized using 0.25% trypsin (Corning) followed by incubation at 37°C in a 5% CO2 incubator. Trypsin neutralization was performed with growth media at room temperature. The cells were then centrifuged for 3 min at 200g. The supernatant was removed followed by resuspension of the cells in fresh in growth media. A fraction of the cells was utilized for the acquisition of flow cytometry measurements (BD LSRFortessa). The rest of the cells were diluted (ratio between 1 and 20) and reseeded into new tissue culture plates for subsequent measurements.

### Cell sorting

Cells were trypsinized using 0.25% trypsin (Corning) followed by incubation at 37°C in a 5% CO2 incubator for 7 minutes. Subsequently, trypsin was neutralized with growth media at room temperature. The cells were then centrifuged at 200g for 3 minutes. The supernatant was discarded and the cell pellet was resuspended in growth media supplemented with 2.5g/ml BSA and corresponding small molecules according to the experiment. The cell suspension was then filtered through a filter cap into flow cytometry tubes. Cell sorting was then performed using a fluorescence-activated cell sorter (BD Aria) approximately 42-48 hours after transfection, and cells were collected in growth media supplemented with penicillin/streptomycin and corresponding small molecules according to the experiment. EBFP2 was measured using a 405nm laser and a 450/50 filter with 332 V. EYFP was measured using a 488nm laser and 515/20 filter with 200 V. Cells were sorted either for transfection level (EYFP) or gene expression level (EBFP2). During cell sorting for EBFP2, a gate was also applied to sort exclusively for EYFP negative cells. Cell sorting to study the gene expression dynamics after epigenetic editing using DNMT3A-dCas9 (Fig. 2D and Fig. SE.2A, B) used ranges 100-200, 222-445, 500-1000 and 1150-2300 EYFP A.U.F.

Fig. 2D shows the 222-445 range. Cell sorting to study the gene expression dynamics after epigenetic editing using PhlF-KRAB (Fig. 2F and Fig. SE.4A-C) used ranges 25-100, 100-1000, 1000-10000, and 10000 - 100000 corresponding to EYFP A.U.F. Fig. 2F shows the 100-1000 range.

Cell sorting to study histone and DNA methylation changes using MeDIP-qPCR and ChIP-qPCR (Fig. 2E, G) used ranges 3000-100000 EYFP A.U.F. Cell sorting to demonstrate the stability of DNMT3A-edited (DNMT3A-dCas9) cells expressing intermediate levels of gene expression (Fig. 3B) was performed according to <100 A.U.F for low expression, 200-500 A.U.F for intermediate expression, and >1000 A.U.F for high expression. Cells had been edited with transfection of DNMT3A-dCas9 targeted to the upstream binding sites of the promoter (Tables SE1 and SE3), cell sorting for 7000-100000 EYFP A.U.F., and were maintained in culture for *>*2 weeks. Cell sorting to subdivide cell populations from stable intermediate levels of gene expression (Fig. 3K) was performed according to <100 A.U.F for bin 1, 179-224 for bin 2, 400-500 for bin 3, >1000 A.U.F for bin 4. Cells expressing intermediate levels of gene expression had been sorted from DNMT3A-edited cells (DNMT3A-dCas9) from the range 300-400 EBFP A.U.F. and maintained in culture for *>*2 weeks. Cells had been edited with transfection of DNMT3A-dCas9 targeted to the upstream binding sites (Tables SE1 and SE3), cell sorting for 3000-100000 EYFP A.U.F. (transfection marker), and maintained in culture for *>*2 weeks. Cell sorting for rTetR-XTEN80-DNMT3A (Fig. 5A) and rTetR-KRAB (Fig. 5C) transfections in cells with stable intermediate levels of expression used 3000-100000 EYFP A.U.F. Cell sorting for rTetR-XTEN80-TET1 (Fig. 5B) transfections in cells with stable intermediate levels of expression used 410-2800 EYFP A.U.F. Cells expressing low and intermediate levels of gene expression had been sorted from DNMT3A-edited cells (DNMT3A-dCas9) using <100 A.U.F for low expression and 200-500 A.U.F for intermediate expression and maintained in culture for *>*2 weeks. Cells had been edited with transfection of DNMT3A-dCas9 targeted to the upstream binding sites (Tables SE1 and SE3), cell sorting for 3000-100000 EYFP A.U.F., and maintained in culture for *>*2 weeks. Cell sorting to study the gene expression dynamics after epigenetic editing using rTetR-XTEN80-DNMT3A (Fig. SE.3A-C) used ranges 60-445, 500-7000, and 7000-100000 EYFP A.U.F. Fig. SE.3A shows the 60-445 range. Cell sorting to demonstrate the stability of DNMT3A-edited (rTetR-XTEN80-DNMT3A) cells expressing intermediate levels of gene expression (Fig. SE.5) was performed according to <80 EBFP2 A.U.F for low expression, and ranges 304-380, 400-500, and 530-662 EBFP2 A.U.F for intermediate expression. The cells had been edited with transfection of rTetR-XTEN80-DNMT3A, supplementation of dox for 8 days, cell sorting for 60-445 EYFP A.U.F., and maintained in culture for *>*2 weeks.

### Single-cell sampling and expansion

Single cell sorting was performed using a fluorescence-activated cell sorter (BD Aria) for various levels of EBFP2 fluorescence. For the cells exhibiting low levels of fluorescence, single cells were obtained from range 21-30 A.U.F. and range 50-60 A.U.F. For the cells exhibiting intermediate levels of fluorescence, single cells were obtained from range 200-240 A.U.F, range 254-305 A.U.F, range 320-384 A.U.F, and range 417-500 A.U.F. For the cells exhibiting high levels of fluorescence, single cells were obtained from range 1000-1200 A.U.F and range 1750-2100 A.U.F. From each range, 90 single cells well obtained for a total of 720 single cells, each seeded on a separate well of a 96-well tissue culture plate (Corning) with growth media supplemented with penicillin/streptomycin. Single cells were then cultured under standard cell culture conditions at 37°C in a 5% CO2 incubator. Growth media was replaced independently for each well every four days. After 14 days of cell culture, 93 monoclonal populations were observed, representing ≥10 monoclonal populations for each of the 8 ranges sampled. The cells in Fig. 3 B-E were used for single-cell sampling at day 29.

### MeDIP-qPCR and ChIP-qPCR

MeDIP-qPCR was performed using the MagMeDIP-qPCR kit (Diagenode) according to the manufacturer’s instructions. ChIP-qPCR was performed using the True MicroChIP-seq kit (C01010132, Diagenode) according to the manufacturer’s instructions. All sonication was performed using a Bioruptor Pico (Diagenode). We utilized the anti-5mC antibody in the MagMeDIP-qPCR kit (Diagenode) for DNA methylation. We utilized antibodies for H3K4me3 (C15410003, Diagenode) and H3K9me3 (C15410056, Diagenode) for histone methylation. The qPCRs were performed using Power SYBR Green PCR Master Mix (Applied Biosystems) and primers (Table SE4) in a LightCycler 96 (Roche). Melt curves were used to confirm a single amplification product and the LightCycler 96 Application Software (Roche) was used to obtain the *C*_*t*_ values. For the reporter gene, the promoter region was amplified. *β*-Actin (H3K4me3) and IgF2 (H3K9me3 and DNA methylation) were used as reference endogenous genes [23]. Folds reported refer to the standard ΔΔ*C*_*t*_ method.

### Targeted bisulfite sequencing

Genomic DNA was extracted using the DNeasy Blood & Tissue Kit (Qiagen) according to the manufacture’s instructions. Samples were then submitted to Zymo Research for conducting the rest of the assay. Assays targeting CpG sites along the chromosomal integration region comprising the continuous sequence covering all DNA binding sites, promoter and EBFP2 were developed using the Rosefinch tool from Zymo Research. Primers were used to amplify bisulfite-converted DNA from the samples, using the EZ DNA Methylation-Lightning Kit for conversion. Amplified products were pooled, barcoded, and prepared for sequencing on an Illumina MiSeq system with a V2 300bp kit, following paired-end sequencing. Sequencing reads were aligned to the chromosomal integration sequence using Bismark. The estimation of methylation levels for each sampled cytosine was calculated by dividing the count of reads identifying a cytosine (C) by the sum of reads identifying either a cytosine (C) or a thymine (T).

### Digital PCR

Genomic DNA extraction was performed with the DNeasy Blood & Tissue Kit (Qiagen), according to the manufacturer’s instructions. Samples were then submitted to Azenta Life Sciences for conducting the rest of the assay. A NanoDrop spectrophotometer was first used to assess sample concentration and confirm purity of the samples. The samples were then loaded in triplicate for analysis on a QIAcuity 8.5k 96-Well Nanoplate using the Qiagen QIAcuity digital PCR system, with probes targeting EBFP2 and COG1. Copy number calculation was conducted using the QIAcuity Software Suite (version 2.2.0.26). Each sample was compared to a no-template control to verify the absence of external contamination.

### Desing of gRNA sequences

Sequences for gRNAs were designed in Benchiling using the chromosomal integration sequence and the CHO-K1 genome sequence (National Library of Medicine (NCBI) RefSeq assembly GCF_000223135.1).

### Protein structure predictions

Protein structure predictions were performed with AlphaFold2 [64] using MMseqs2 on the Google Colab Platform (ColabFold v1.5.5)[65] on GPUs. AlphaFold2 was used to perform structural predictions using 5 models for each protein. The top ranking predicted structure (according to plDDT score for single chains and pTM score for protein complexes) was then relaxed using Assisted Model Building with Energy Refinement (AMBER) relaxation. The protein structures of PhlF-KRAB, rTetR-KRAB, PhlF, and rTetR were predicted as multimers (homodimers). The protein structures of DNMT3A-dCas9, rTetR-XTEN80-DNMT3A, rTetR-XTEN80-TET1 were predicted as single chains. The structures of rTetR-XTEN80-TET1 and rTetR-XTEN80-DNMT3A were then aligned using the predicted homodimer of rTetR as a template in PyMOL (2.5.2). Minimal rotations (≤1 per linker) were performed at the flexible linkers in PyMOL.

### Quantification and statistical analysis

To determine the probability of a specific fraction of CpGs being methylated based on bisulfite sequencing data, which offers the probability of a specific CpG being methylated for the cell population, we employ the following method. This approach is applied to both bisulfite sequencing datasets presented in Figs. 3G and 4E, resulting in the corresponding plots in Figs. 3H and 4H, respectively.

Let us assume that the gene of interest has a total of *N* CpGs. Then, for any 0 ≤ *j* ≤ *N*, having *j* CpGs methylated corresponds to the event where precisely *j* CpGs are methylated, and all other CpGs (i.e., *N* − *j*) are unmethylated. Now, let us define the probability of the described event as *P*_*j*_, where 0 ≤ *j* ≤ *N*. Additionally, let the probability of CpG *i* being methylated be *p*_*i*_, and the probability of CpG *i* being unmethylated be *q*_*i*_, where 1 ≤ *i* ≤ *N*. The bisulfite sequencing data provide us with *p*_*i*_, and since the event “CpG unmethylated” is the complement of “CpG methylated”, *q*_*i*_ is known and can be expressed as *q*_*i*_ = 1 − *p*_*i*_. Then, the probability that exactly *j* CpGs are methylated out of a total of *N* CpGs, where each CpG has its own probability of being methylated (*p*_*i*_) or not being methylated (*q*_*i*_), can be calculated by using the following formula:

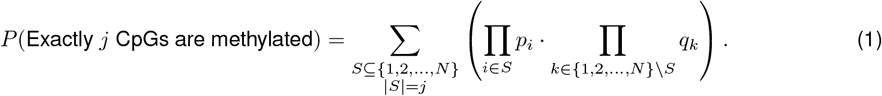

More precisely, the sum (Σ) is taken over all possible subsets *S* of events with exactly *j* elements. For each subset *S*, we calculate the product of the probabilities of the events in *S* occurring (Π_*i*∈*S*_ *p*_*i*_). We then calculate the product of the probabilities of the events not in *S* occurring (Π_*k*∈{1,2,…,*N* }\*S*_ *q*_*k*_). The overall probability is the sum of these products for all subsets *S*. In simpler terms, the formula considers all possible combinations of *j* events occurring out of *N* events, multiplying the probabilities of those events occurring and the probabilities of the remaining events not occurring in each combination. The sum aggregates these probabilities for all valid combinations, providing the probability that exactly *j* events happen.

As an illustrative example, let us consider the case in which the total number of CpG in the gene of interest are *N* = 3. Then, defining the probability of CpG *i* to be and not to be methylated as *p*_*i*_ and *q*_*i*_ = 1 − *p*_*i*_, with *i* = 1, 2, 3, we then can calculate the probability of having *j* CpGs methylated, i.e., *P*_*j*_, with *j* = 0, 1, 2, 3, using (1), obtaining *P*_0_ = *q*_1_*q*_2_*q*_3_, *P*_1_ = *p*_1_*q*_2_*q*_3_+*q*_1_*p*_2_*q*_3_+*q*_1_*q*_2_*p*_3_, *P*_2_ = *p*_1_*p*_2_*q*_3_+*p*_1_*q*_2_*p*_3_+*q*_1_*p*_2_*p*_3_, and *P*_3_ = *p*_1_*p*_2_*p*_3_. The probability distribution of the fraction of CpGs methyalated is then obtained by plotting (*j, P*_*j*_) for *j* = 0, 2, …, *N*.

Concerning the best-fit linear regression lines shown in Figs. 3I, 4I, and 4J, they were obtained using MATLAB polyfit function. Furthermore, the p-value associated with the slope coefficient was determined through the MATLAB fitlm function.

For statistical tests comparing percentages calculated from flow cytometry measurements or fold changes from MeDIP-qPCR and ChIP-qPCR, unpaired two sample t-tests were performed.

### Computational analysis

All simulations conducted in this paper were done using Gillespie’s Stochastic Simulation Algorithm (SSA) [52].

## Author contributions

S.P., S.B., and D.D.V. conceptualized the study. D.D.V. directed and supervised the study. S.P., S.B., R.W., E.S., A.K. and D.D.V. contributed to experimental design and/or data analysis. S.P., E.S., A.K. and K.I. conducted or assisted experimental investigation. S.B. performed mathematical modeling and data analysis. S.P., S.B. and D.D.V. wrote the manuscript.

## Data and materials availability

Data to evaluate the conclusions of the study will be in the manuscript and supplementary materials before publication. Original data is available from the corresponding author upon request. All sequences used in the study are to be included in the manuscript, and plasmids will be deposited in Addgene before publication. Targeted bisulfite sequencing data will be available from a public repository before publication.

## Code availability

Code for mathematical modeling is available from the corresponding author upon request.

## Acknowledgements

We thank Dr. James J. Collins for providing scientific comments and suggestions on the link between the model predictions and the experimental data, Kalon Overholt for providing useful references on DNA methylation and providing feedback on some of the figures, and the Biological Engineering Comm Lab for helping us refine some of the diagrams. We would like to thank the Koch Institute’s Flow Cytometry Core at MIT where the cell sorting was performed. Furthermore, we would like to thank our funding: NSF-MODULUS, Award number 2027949.

## Supplementary Information (SI)

### S.1 Model of chromatin modification circuit

The chromatin modification circuit reaction model considered in our study, shown in Fig. SM.1A, was constructed starting from the one in [66] and it is based on known molecular interactions from the literature. Here, we give a brief description of the reaction model (see [66] for additional details). The chromatin modifications considered in the circuit are H3K9 methylation (H3K9me3), H3K4 methylation (H3K4me3), and DNA methylation. The basic unit of the model is the nucleosome with DNA wrapped around it, D, that can be modified with H3K4me3, D^A^, DNA methylation,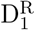, H3K9me3, 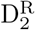, or both H3K9me3 and DNA methylation, 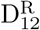. The reaction model can be graphically represented by the circuit of Fig. SM.1A, whose associated reactions are given in SI Section S.6, Table SM.1. More precisely, writer enzymes have the ability to *de novo* establish chromatin marks. Additionally, histone modifications recruit writer enzymes for the same modification to nearby modifiable nucleosomes (*auto-catalysis*) [13, 21, 22, 46], and DNA methylation and repressive histone modification cooperate by recruiting each other’s writer enzymes (*cross-catalysis*) [38]. Ultimately, these modifications can be passively removed through dilution during DNA replication or by the action of eraser enzymes (*basal erasure*). Both activating and repressive modifications are involved in the recruitment of each other’s eraser enzymes (*recruited erasure*) [14, 43–45].

Now, defining the number of 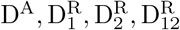 as 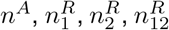 and *n*^*D*^, let us derive the related ordinary differential equation (ODE) model in terms of the fractions 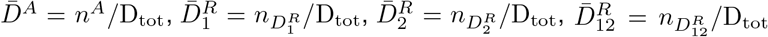,and 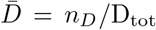, with D_tot_ the total number of nucleosomes in a gene of interest. This can be done by assuming that D_tot_ is sufficiently large so that 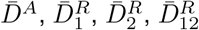 and 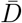 can be treated as real number. Now, let us introduce *D*_*tot*_ = D_tot_*/*Ω where Ω is the reaction volume, and define the normalized inputs: 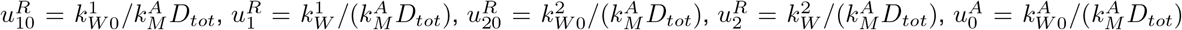, and 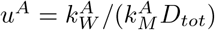. Furthermore, let us introduce the following dimensionless parameters:

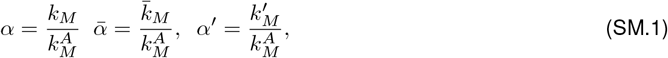

in which 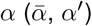 is the non-dimensional rate constant associated with auto-catalysis (cross-catalysis). Let

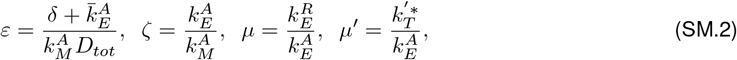

with *b* = *O*(1) such that 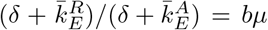 and *β* = *O*(1) such that 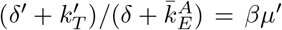. Then, *µ* (*µ*′) encapsulates the asymmetry between the erasure rate of repressive histone modifications (the DNA demethylation rate) and the erasure rate of activating marks. Furthermore, given that 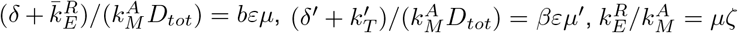 and 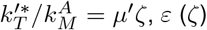 is a parameter that scales the ratio between the basal (recruited) erasure rate and the auto-catalysis rate of each modification. Considering the normalized time 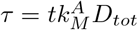, the ODEs associated with the chromatin modification circuit are

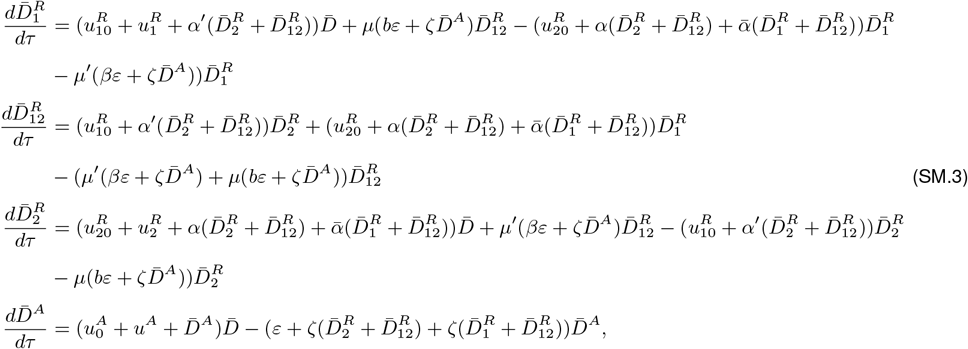

with 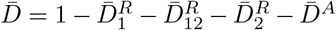 (see [66] for full derivation). It is important to note that in our reporter gene, the expression of fluorescent protein EBFP2 is driven by a constitutive promoter. This means that the promoter remains active continuously under normal conditions, without the need for external activators. This is the reason why, for D^A^, we introduce a constant non-basal *de-novo* establishment term (*u*^*A*^ *>* 0). As for the other species, we will describe in the following section how external inputs, such as KRAB, DNMT3A, and TET1, modulate the parameters of the chromatin modification circuit model.

#### S.1.1 How KRAB, DNMT3A and TET1 modulate model parameters

In our study, we exploit KRAB, DNMT3A, and TET1 to modulate chromatin modification state and they affect the model parameters as follows. KRAB is a well-known epigenetic effector that mediates the recruitment of H3K9me3 writers [40]. As we introduce KRAB through transient transfection, its concentration will gradually decrease due to dilution until it eventually dies out. Furthermore, KRAB is fused to the DNA binding domain (DBD) PhlF. This fusion not only allows us to obtain sequence-specific recruitment, but also to regulate the binding and unbinding of KRAB through DAPG, as DAPG inhibits the ability of PhlF-KRAB to bind to DNA. To incorporate these aspects into our model, we write the expression for 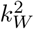, i.e., reaction rate constant for H3K9me3 establishment (see reactions in SI Table SM.1), as follows:

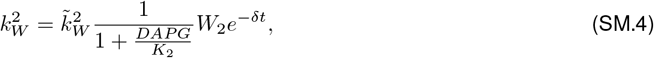

in which *W*_2_ is the total amount of PhlF-KRAB, 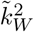 is a parameter independent of PhlF-KRAB, *K*_2_ is the dissociation constant related to the binding - unbinding reactions between DAPG and PhlF-KRAB, and *δ* is the dilution rate constant. Then, the normalized input 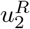 in the ODE model (SM.3) can be written as

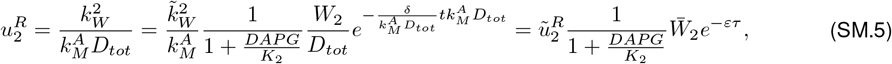

in which we define 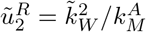 and 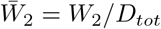.

The enzyme DNMT3A is a DNA methyltransferase that catalyzes *de novo* DNA methylation ([13], Chapter 5) and, as a consequence, it modulates the rate of *de novo* DNA methylation establishment 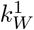 (see reactions in SI Table SM.1). As for KRAB, we introduce DNMT3 through transient transfection, and therefore its concentration will gradually decrease. To incorporate this aspect into our model, we can modify the expression for 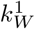 as follows:

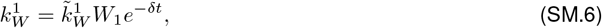

in which *W*_1_ is the total amount of DNMT3A and 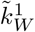 is a parameter independent of DNMT3A. Then, the normalized input 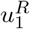 in the ODE model (SM.3) can be written as

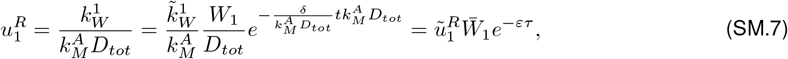

in which we define 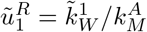 and 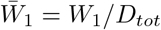.

Finally, TET1 is an enzyme that catalyzes the conversion of DNA methylation into modifications that are not recognized by the maintenance enzyme DNMT1, and hence are passively removed through dilution [14, 43, 67]. This enzyme then modulates the constants 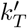 and 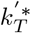 in the reactions of SI Table SM.1. As for KRAB, we introduce TET1 through transient transfection and we fuse it to a DBD. More specifically, we employ rTetR, a DBD that binds DNA exclusively in the presence of doxycycline (dox). To incorporate these aspects into our model, we can modify the expressions for 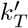 and 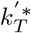 as follows:

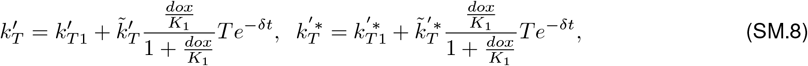

in which 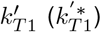 is the component of the rate coefficient that does not depend on the external rTetR-TET1 transiently transfected, *T* is the total amount of rTetR-TET1 transiently transfected, 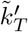 and 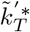 are parameters independent of rTetR-TET1, and *K*_1_ is the dissociation constant related to the binding - unbinding reactions between dox and rTetR-TET1. Then, the non-dimensional parameter *µ*′ in the ODE model (SM.3) can be written as

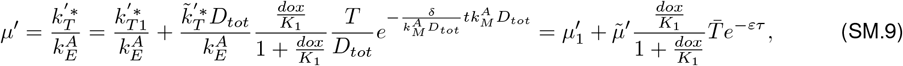

in which we define 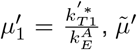, and 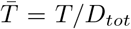. From (SM.9), it is possible to note that, without transfection of rTetR-TET1, 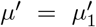, indicating that *µ*′ is non-zero. This is because of the presence of passive DNA demethylation. However, previous experimental studies suggest that passive DNA demethylation, although present, is a slow process [23, 42].

### S.2 Model of transcriptional regulation

The gene expression process involves two main steps. The first step is transcription, during which the genetic information is transcribed from DNA into mRNA (m). The second step is translation, during which the mRNA is translated into the gene product (X). Chromatin state affects transcription by modulating nucleosome compaction and then gene expression [13, 68]. Thus, we assume that transcription is predominantly allowed by D^A^, while allowing a low level of transcription to D and to all the species characterized by repressive chromatin modifications. Furthermore, mRNA m is subject to dilution due to cell division and degradation, while the gene product X is subject to dilution only. This is because fluorescent reporters are highly stable and thus not degraded but only diluted [69]. The reactions associated with the gene expression model can be found in SI Table SM.4. Considering the normalized time 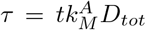 and introducing the non-dimensional parameters 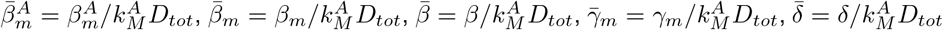, the ODE model for X and m can be written as

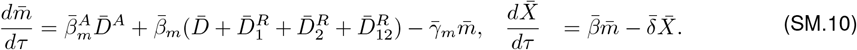

In this paper, we will refer to the model that combines the complete chromatin modification circuit model (SI Section S.1, equations (SM.3)) along with the transcriptional regulation model outlined in this section as the “4D+X model”.

### S.3 Effect of *α*′ and *µ*′ on the behavior of the system

In this section, we analyze the stochastic behavior of the 4D+X model with a specific focus on understanding how the parameters *α*′ (normalized rate of DNA methylation establishment through repressive histone modifications) and *µ*′ (ratio between the DNA demethylation rate and the activating histone modification erasure rate) affect the probability distribution of gene expression levels and thus the type of memory that can be achieved.

When the rate of DNA demethylation is 0 (*µ*′ = 0), achieving analog memory is possible only when repressive histone modification (H3K9me3) does not catalyze the *de novo* establishment of DNA methylation (*α*′ = 0) (Fig. SM.1B). For *α*′ *>* 0, the gene expression level shifts either to a low or high level. Furthermore, the higher *α*′, the more the distribution tends to shift towards a low gene expression level (Fig. SM.2 - top panel). When *µ*′ is different from 0, only binary memory can be achieved (Fig. SM.2 - intermediate and bottom panels). Moreover, the higher *µ*′ is, the more the gene expression level shifts towards a high level (Fig. SM.2 - bottom panel). Overall, these results suggest that analog memory can be achieved only when DNA demethylation rate is sufficiently small compared to histone modification erasure rate (that is, *µ*′ can be approximated as zero on the time scales of interest) and DNA methylation is not catalyzed by repressive histone modifications (*α*′ = 0).

If these conditions are verified, and external inputs (KRAB, DNMT3A, and TET1) are not applied or are applied transiently (transient transfection), then the number of nucleosomes with methylated DNA stabilizes at a constant value. Introducing the variable 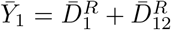, that is, the fraction of nucleosomes with methylated DNA in the gene, the behavior of the original model (SM.3) can be captured by a reduced 2D ordinary differential equation (ODE) model below:

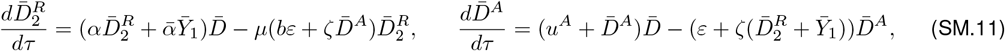

with 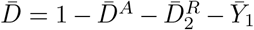 and 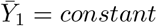(full derivation below). A representative diagram of the reduced system is depicted in Fig. SM.1C. It is important to point out that in the original model of chromatin modification circuit, it is assumed that the basic unit is a nucleosome with DNA wrapped around it and with only one modifiable CpG [66]. There are cases in which this assumption is not verified. However, it is possible to demonstrate that removing this assumption yields a similar reduced model. This indicates that the assumption does not impact the qualitative results regarding the influence of the parameters *α*′ and *µ*′ on the probability distribution of gene expression levels (see SI Section S.7).

#### Derivation of model (SM.11)

Let us first merge the rates linked to the enhancement of H3K9me3 establishment by 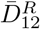 into a single rate. We will assume that this rate is identical to the one associated with the enhancement of H3K9me3 establishment by 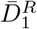. Similarly, let us merge the rates linked to the recruited erasure of H3K4me3 by 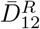 into a single rate and assume that this rate is identical to the one associated with the recruited erasure of H3K4me3 by 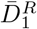. These simplifying assumptions do not affect the qualitative results related to the effect of the cooperative and competitive interactions among chromatin modifications on epigenetic cell memory. The ODE system (SM.3) can then be rewritten as

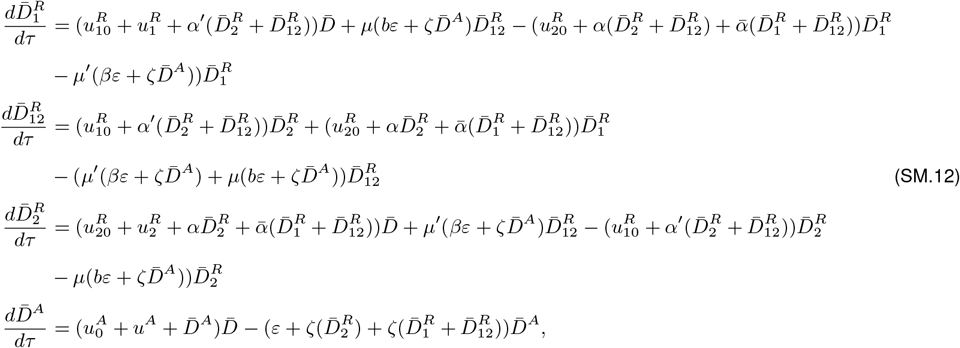

Now, let us rewrite system (SM.12) by assuming negligible basal *de novo* establishment 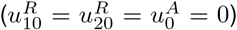 and introducing the variable 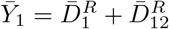, which corresponds to the fraction of nucleosomes with methylated DNA. We then obtain

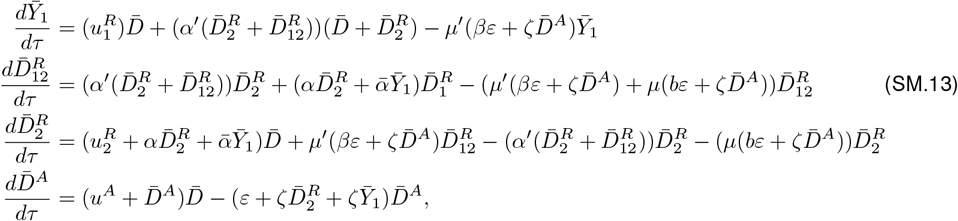

with 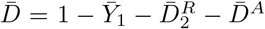. Based on previous experimental results [23] and our experimental findings (see Fig. 2G), we can further simplify our model by accounting for the fact that the enhancement of DNA methylation establishment by H3K9me3 is negligible, i.e., *α*′ = 0. The ODE system (SM.13) can then be rewritten as

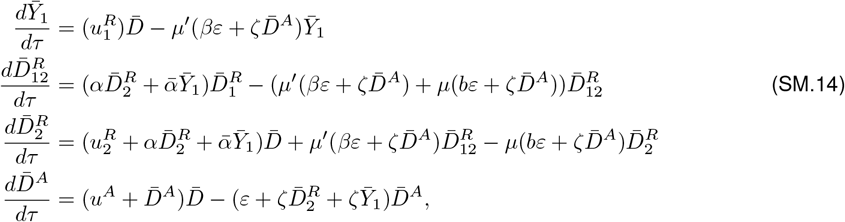

with 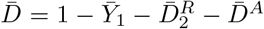. Now, let us rewrite the ODE system (SM.14) introducing the expressions for 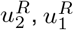 and *µ*′ derived in SI Section S.1.1 (Expressions (SM.5), (SM.7), and (SM.9)):

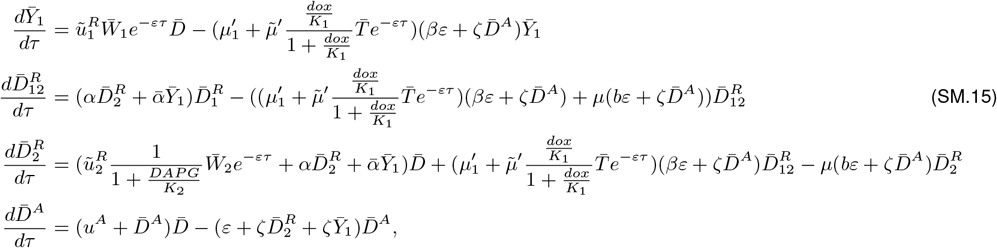

with 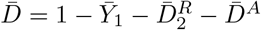 and 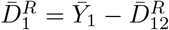. Let us now consider the parameter regime in which the rate of DNA demethylation, without transfection of rTetR-TET1, is sufficiently low compared to histone modification dynamics, i.e.,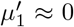. This assumption is consistent with experimental data that suggest that the passive DNA demethylation is a slow process [23, 42]. The system (SM.15) can then be rewritten as follows:

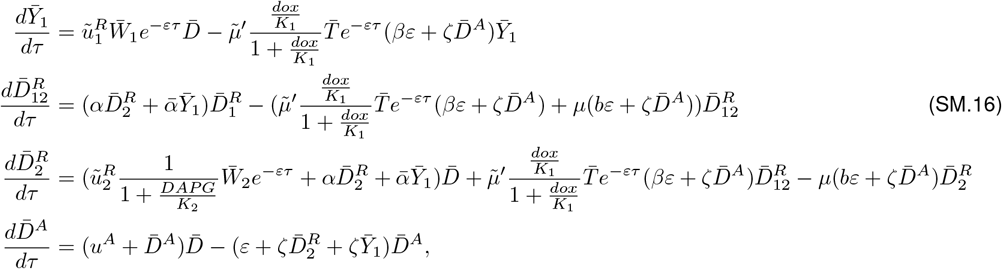

with 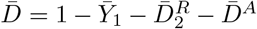 and 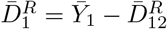. From (SM.16), it follows that, after a temporary phase during which the externally transfected inputs decrease, *e*^−*ετ*^ ≈ 0 and then 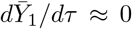, and the last two ODEs in (SM.16) depend only on 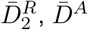, and 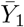. This implies that, after a transient change, 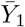 stabilizes at a constant value and the behavior of the original model can be captured by the reduced two-dimensional ODE model given by the last two equations in (SM.16), that corresponds to the ODE model in (SM.11).

### S.4 Effect of ε and ζ on the behavior of the system

In this section, we analyze the deterministic and stochastic behavior of our system, with a specific focus on understanding the effect of parameters *ε* (parameter that scales the ratio between the basal erasure rate and the auto-catalysis rate of each modification) and *ζ* (parameter that scales the ratio between the recruited erasure rate and the auto-catalysis rate of each modification) on the probability distribution of gene expression levels.

We first conduct a deterministic analysis by studying the ODE model associated with the simplified chromatin modification circuit, i.e., equations (SM.11), and determine how *ε* and *ζ* affect the value of 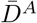 at the equilibrium for different fractions of methylated CpGs in the gene, i.e., 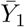 (Fig. SM.3A). When *ε* is large, the system consistently exhibits a unique stable steady state characterized by low 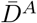 (Fig. SM.3A). The value of 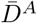 is smaller for higher 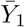. As *ε* decreases, the steady-state value of *D*^*A*^ increases, particularly for low values of 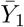, where 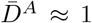(Fig. SM.3A). Further reduction in *ε* results in the system becoming bistable for intermediate values of 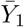 (Fig. SM.3A). Changes in *ζ* do not significantly impact these trends, except for when *ε* is small. In such cases, larger values of *ζ* reduce the range of 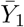 for bistability and diminish the difference in the values of 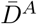 between the two steady states (Fig. SM.3A). We next study the impact of *ε* and *ζ* on the number of methylated CpGs at equilibrium for various initial levels of DNMT3A, represented by 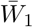 as defined in SI Eq. (SM.6) (Fig. SM.3B). In general, 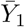 increases with higher 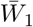. However, for large values of *ε*, 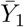 remains low over an extended range of 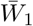 values. As *ε* decreases, 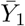 reaches higher values. In the case of small *ε*, an increase in *ζ* results in a more ultrasensitive curve.

Overall, this analysis suggests that high fractions of nucleosomes with H3K4me3, and then a high level of gene expression, is achievable only when *ε* is sufficiently small (Fig. SM.3A). In this parameter regime, when *ζ* is sufficiently high, 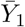 has an ultrasensitive response to DNMT3A transient dosage and then different ranges of DNMT3A pulse height would lead to either high or low gene expression levels only (Fig. SM.1D and Fig. SM.3B). To validate these findings, we use the Stochastic Simulation Algorithm (SSA) [52] to perform a computational study on the 4D+X model, that is, the model that incorporates the full chromatin modification circuit, whose reactions are listed in SI Tables SM.1 and SM.4 (Fig. SM.1E and Fig. SM.3C). Here, we consider several ranges of the initial amount of DNMT3A transfected 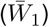, which are uniformly distributed.

For different ranges of initial DNMT3A transfection levels, a bimodal probability distribution of gene expression levels emerges when *ζ* is sufficiently large (Fig. SM.3C, left-hand side plots). As *ζ* decreases, the stationary distribution becomes unimodal (Fig. SM.3C, left-hand side plots). When considering ranges that include higher values of 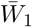, it becomes more evident how the unique peak of the distribution shifts towards lower gene expression levels for sufficiently high values of 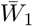(Fig. SM.1E, left-hand side plots and Fig. SM.3C, right-hand side plots).

We then conduct a similar computational analysis to assess the impact of *ε* and *ζ* on number of methylated CpGs and gene expression level across various initial levels of TET1 enzyme 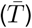 (Fig. SM.3D,E). The obtained results mirror those described in the previous analysis (Fig. SM.3B,C).

By looking at our experimental results involving transfection of KRAB (Fig. 2F), it is possible to note that the probability distribution of gene expression levels is bimodal. Previous computational study suggests that this behavior can be obtained only for sufficiently small values of *ε* [66, 70]. Furthermore, our experimental results involving transfection of DNMT3A (Fig. 2D and SI Fig. SE.2A) also show a bimodal probability distribution for gene expression levels. Based on our mathematical analysis (Fig. SM.3B,C), these outcomes can be observed only when *ε* is sufficiently small and *ζ* is sufficiently large. We validate these hypotheses by successfully replicating our experimental data via simulations considering this parameter regime, i.e, small *ε*, large *ζ* (Fig. SM.5).

### S.5 Refined chromatin modification circuit model to replicate the bisulfite sequencing data from the monoclonal experiment (Fig. 4)

In the study conducted in the previous sections, we observed that, in the absence of external stimuli and when *α*′ and *µ*′ can be assumed equal to zero, the chromatin modification circuit model, whose associated reactions are listed in Table SM.1, predicts a “frozen” level of DNA methylation, i.e., 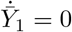 (SI Section S.3). However, this model does not perfectly align with the experimental results obtained from the bisulfite sequencing data of the synthetic reporter system in the eight different clones, each once associated with a distinct EBFP2 expression level (Fig. 4A, B, D - G). In fact, for each clone, we do not observe a single value for methylated CpGs across the entire cell population. Instead, we observe a distribution with mean and standard deviation that remain approximately constant throughout the observed time points (Fig. 4G). Notably, the distribution of methylated CpGs appears more spread for clones associated with an intermediate level of methylated CpGs.

In order to recapitulate this observed behavior, we incorporated in the full model known molecular mechanisms involved in the DNA methylation maintenance process that were omitted before. Specifically, in the original full model we have

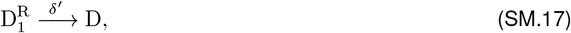

in which *δ*′ represents the net passive demethylation rate constant obtained from the balance between the dilution and the DNMT1 maintenance process. In order to recapitulate the experimental results, we added the following reactions:

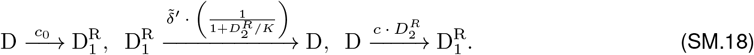

It is important to point out that, for the sake of simplicity in the description of the DNA methylation maintenance model above, we used, with abuse of notation, 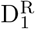 to denote any species with DNA methylation and 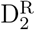 to denote any species with H3K9me3. The first reaction in (SM.18) represents the possibility that the DNA methylation maintenance enzyme DNMT1 can cause *de novo* DNA methylation, although at a very low rate (low *c*_0_) [49, 50]. The second reaction represents the fact that H3K9me3 helps recruit DNMT1 to the gene and then reinforces DNMT1-mediated maintenance [48]. Finally, we introduce the third reaction since H3K9me3 helps recruit DNMT1 and DNMT1 can, by mistake, establish DNA methylation on previously unmethylated sites.

Considering the normalized time 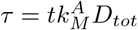 and introducing the non-dimensional parameters 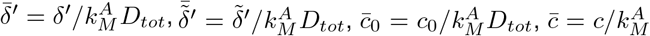, the ODE describing the dynamics of fraction of nucleosomes with methylated DNA, that is, 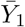, can be then rewritten as

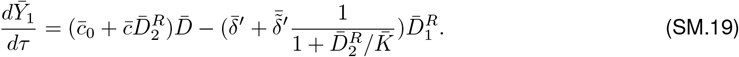

Then, the average level of DNA methylation remains approximately constant in time when 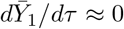, that is, if all the reaction rate constants are sufficiently small 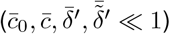.

This is confirmed by the computational study conducted by simulating the reactions of the refined chromatin modification circuit model that incorporates the DNA methylation maintenance model described above, whose reactions are listed in SI Table SM.6, using the SSA (SI Fig. SM.7A). Additionally, this analysis shows that the reactions encapsulating the recruitment mechanism of DNMT1 by H3K9me3 (the second and the third in (SM.18)) are crucial for obtaining broader distributions in correspondence of intermediate levels of methylated CpGs (SI Fig. SM.7B). In fact, as illustrated in the figure, when the rate constants associated with these reactions are low, all distributions have approximately the same height and width. However, higher values of these rates result in broader distributions at intermediate levels of methylated CpGs.

### S.6 4D+X model: reactions

#### S.6.1 Chromatin modification circuit

In this reaction model, it is assumed that the rate constants of the establishment, auto and cross-catalytic, and erasure processes of H3K9me3 (DNA methylation) do not change if the other repressive mark is present on the same modifiable unit. The chromatin modification circuit reaction model can then be written as in SI Table SM.1.

**Table SM.1:**
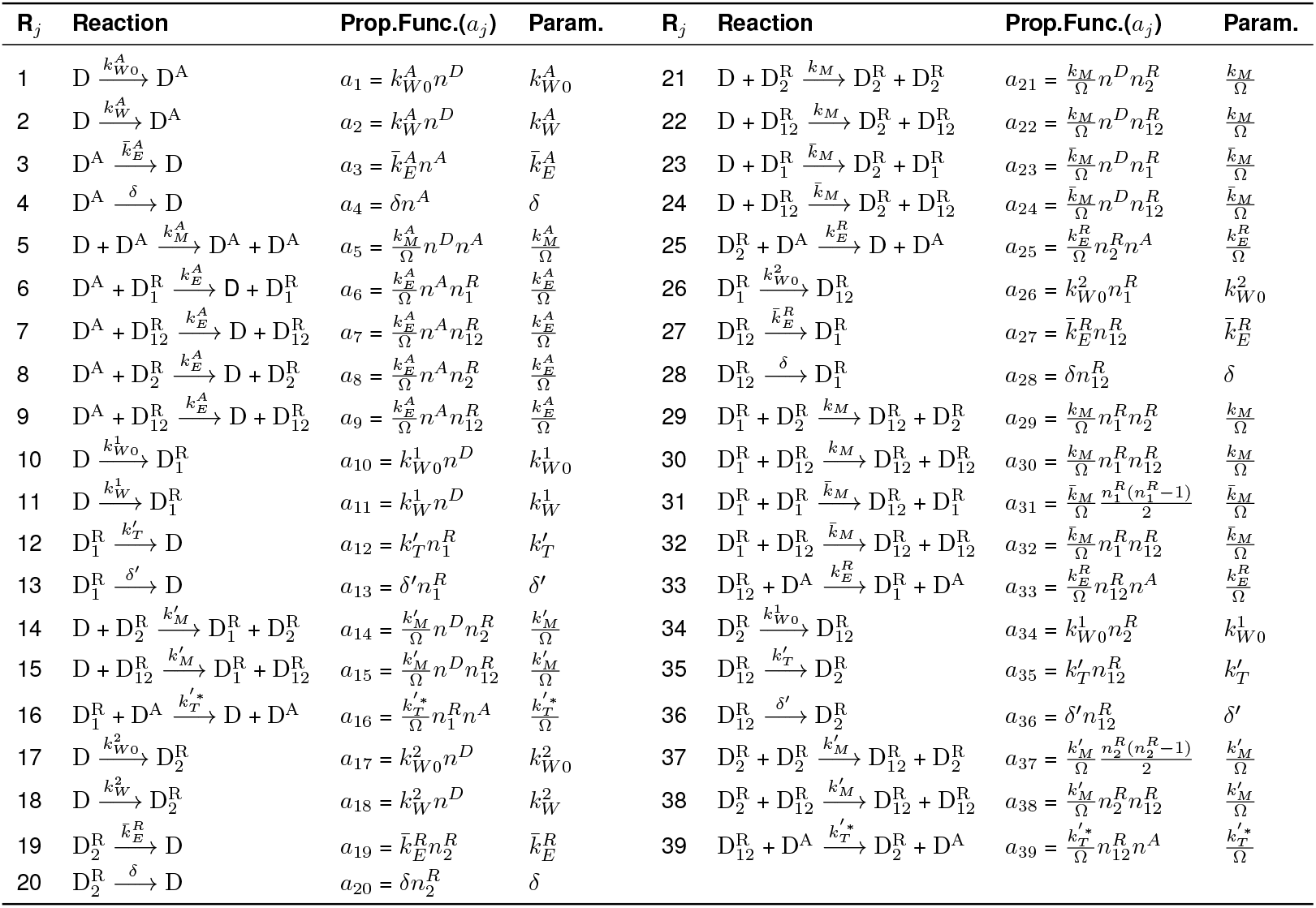
Full chromatin modification circuit model: reactions.

**Table SM.2:**
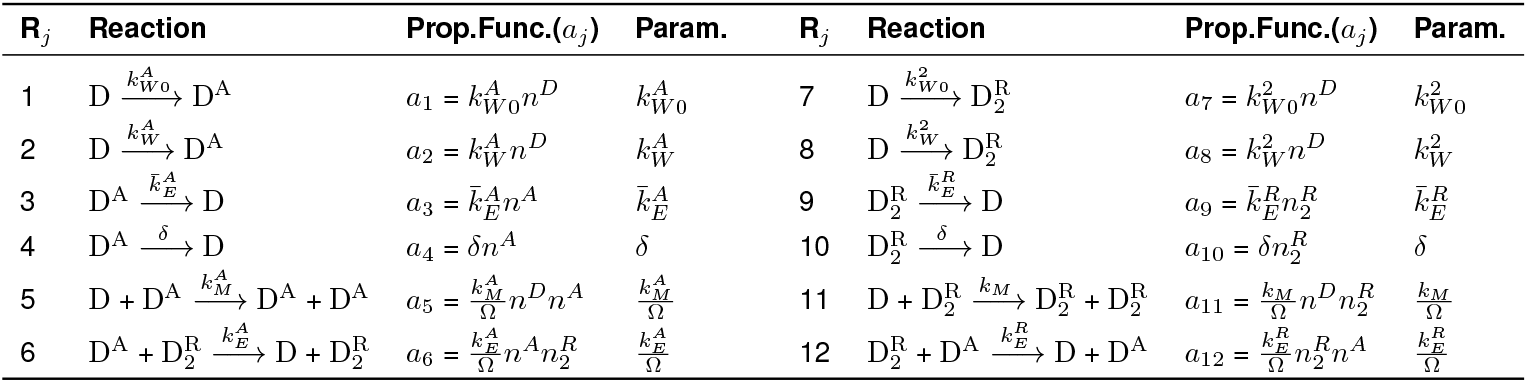
Histone modification circuit model: reactions.

When DNA methylation is not included in the system, the only species included in the model are D, D^A^, and 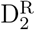, and the reactions included in the model become only the ones listed in SI Table SM.2.

Finally, the model represented in Fig. 2H can be obtained starting from the full chromatin modification circuit model (SI Table SM.1) and assuming negligible basal *de novo* establishment 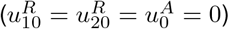, merging the rates linked to the enhancement of H3K9me3 establishment by 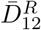 into a single rate, merging the rates linked to the recruited erasure of H3K4me3 by 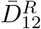 into a single rate, assuming negligible DNA methylation establishment by H3K9me3, and assuming a DNA demethylation rate, without transfection of rTetR-TET1, sufficiently low compared to histone modification dynamics. The reaction list associated with this model can be written as in SI Table SM.3 and the corresponding ODE model can be found in (SM.16).

**Table SM.3:**
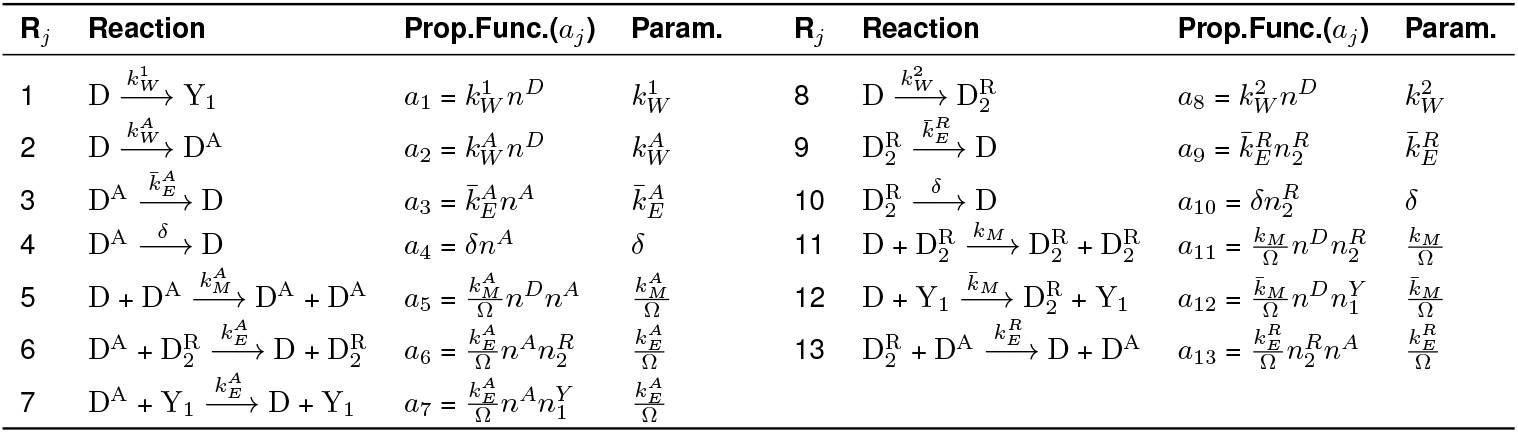
Chromatin modification circuit model represented in Fig. 2H: reactions.

| $R_j$ | Reaction | Prop.Func.($a_j$) | Param. | $R_j$ | Reaction | Prop.Func.($a_j$) | Param. |
| --- | --- | --- | --- | --- | --- | --- | --- |
| 1 | $D \xrightarrow{k_W^1} Y_1$ | $a_1 = k_W^1 n^D$ | $k_W^1$ | 8 | $D \xrightarrow{k_W^2} D_2^R$ | $a_8 = k_W^2 n^D$ | $k_W^2$ |
| 2 | $D \xrightarrow{k_W^A} D^A$ | $a_2 = k_W^A n^D$ | $k_W^A$ | 9 | $D_2^R \xrightarrow{\bar{k}_E^R} D$ | $a_9 = \bar{k}_E^R n_2^R$ | $\bar{k}_E^R$ |
| 3 | $D^A \xrightarrow{\bar{k}_E^A} D$ | $a_3 = \bar{k}_E^A n^A$ | $\bar{k}_E^A$ | 10 | $D_2^R \xrightarrow{\delta} D$ | $a_{10} = \delta n_2^R$ | $\delta$ |
| 4 | $D^A \xrightarrow{\delta} D$ | $a_4 = \delta n^A$ | $\delta$ | 11 | $D + D_2^R \xrightarrow{k_M} D_2^R + D_2^R$ | $a_{11} = \frac{k_M}{\Omega} n^D n_2^R$ | $\frac{k_M}{\Omega}$ |
| 5 | $D + D^A \xrightarrow{k_M^A} D^A + D^A$ | $a_5 = \frac{k_M^A}{\Omega} n^D n^A$ | $\frac{k_M^A}{\Omega}$ | 12 | $D + Y_1 \xrightarrow{\bar{k}_M} D_2^R + Y_1$ | $a_{12} = \frac{\bar{k}_M}{\Omega} n^D n_1^Y$ | $\frac{\bar{k}_M}{\Omega}$ |
| 6 | $D^A + D_2^R \xrightarrow{k_E^A} D + D_2^R$ | $a_6 = \frac{k_E^A}{\Omega} n^A n_2^R$ | $\frac{k_E^A}{\Omega}$ | 13 | $D_2^R + D^A \xrightarrow{k_E^R} D + D^A$ | $a_{13} = \frac{k_E^R}{\Omega} n_2^R n^A$ | $\frac{k_E^R}{\Omega}$ |
| 7 | $D^A + Y_1 \xrightarrow{k_E^A} D + Y_1$ | $a_7 = \frac{k_E^A}{\Omega} n^A n_1^Y$ | $\frac{k_E^A}{\Omega}$ | | | | |

#### S.6.2 Transcriptional regulation

The reactions associated with the gene expression model described in SI Section S.2 are listed in SI Table SM.4. Here, m is the mRNA, X is the gene product, *γ*_*m*_ is the decay rate constant of m due to dilution and degradation, *δ* is the decay rate constant of X due to dilution, 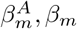 are the transcription rate constants, with 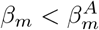, and *β* is the translation rate constant.

**Table SM.4:** Gene expression model: reactions.

| $R_j$ | Reaction | Prop.Func.($a_j$) | Param. | $R_j$ | Reaction | Prop.Func.($a_j$) | Param. |
| --- | --- | --- | --- | --- | --- | --- | --- |
| 1 | $D^A \xrightarrow{\beta_m^A} D^A + m$ | $a_1 = \beta_m^A n^A$ | $\beta_m^A$ | 5 | $D_{12}^R \xrightarrow{\beta_m} D_{12}^R + m$ | $a_5 = \beta_m n_{12}^R$ | $\beta_m$ |
| 2 | $D \xrightarrow{\beta_m} D + m$ | $a_2 = \beta_m n^D$ | $\beta_m$ | 6 | $m \xrightarrow{\beta} m + X$ | $a_6 = \beta n^m$ | $\beta$ |
| 3 | $D_1^R \xrightarrow{\beta_m} D_1^R + m$ | $a_3 = \beta_m n_1^R$ | $\beta_m$ | 7 | $m \xrightarrow{\gamma_m} \emptyset$ | $a_7 = \gamma_m n^m$ | $\gamma_m$ |
| 4 | $D_2^R \xrightarrow{\beta_m} D_2^R + m$ | $a_4 = \beta_m n_2^R$ | $\beta_m$ | 8 | $X \xrightarrow{\delta} \emptyset$ | $a_8 = \delta n^X$ | $\delta$ |

When DNA methylation, and then species 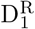 and 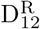 are not included in the chromatin modification circuit, then the gene expression model can be simplified as in SI Table SM.5.

**Table SM.5:** Gene expression model when we remove species 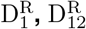: reactions.

| $R_j$ | Reaction | Prop.Func. ($a_j$) | Param. | $R_j$ | Reaction | Prop.Func. ($a_j$) | Param. |
| --- | --- | --- | --- | --- | --- | --- | --- |
| 1 | $D^A \xrightarrow{\beta_m^A} D^A + m$ | $a_1 = \beta_m^A n^A$ | $\beta_m^A$ | 4 | $m \xrightarrow{\beta} m + X$ | $a_4 = \beta n^m$ | $\beta$ |
| 2 | $D \xrightarrow{\beta_m} D + m$ | $a_2 = \beta_m n^D$ | $\beta_m$ | 5 | $m \xrightarrow{\gamma_m} \emptyset$ | $a_5 = \gamma_m n^m$ | $\gamma_m$ |
| 3 | $D_2^R \xrightarrow{\beta_m} D_2^R + m$ | $a_3 = \beta_m n_2^R$ | $\beta_m$ | 6 | $X \xrightarrow{\delta} \emptyset$ | $a_6 = \delta n^X$ | $\delta$ |

#### S.6.3 DNA methylation maintenance and refined chromatin modification circuit

To recapitulate the experimental results shown in Fig. 4G, modifications need to be made to the reaction system that models the DNA methylation maintenance process, as explained in SI Section S.5. The reaction model for the chromatin modification circuit can then be revised and expressed as detailed in SI Table SM.6.

**Table SM.6:**
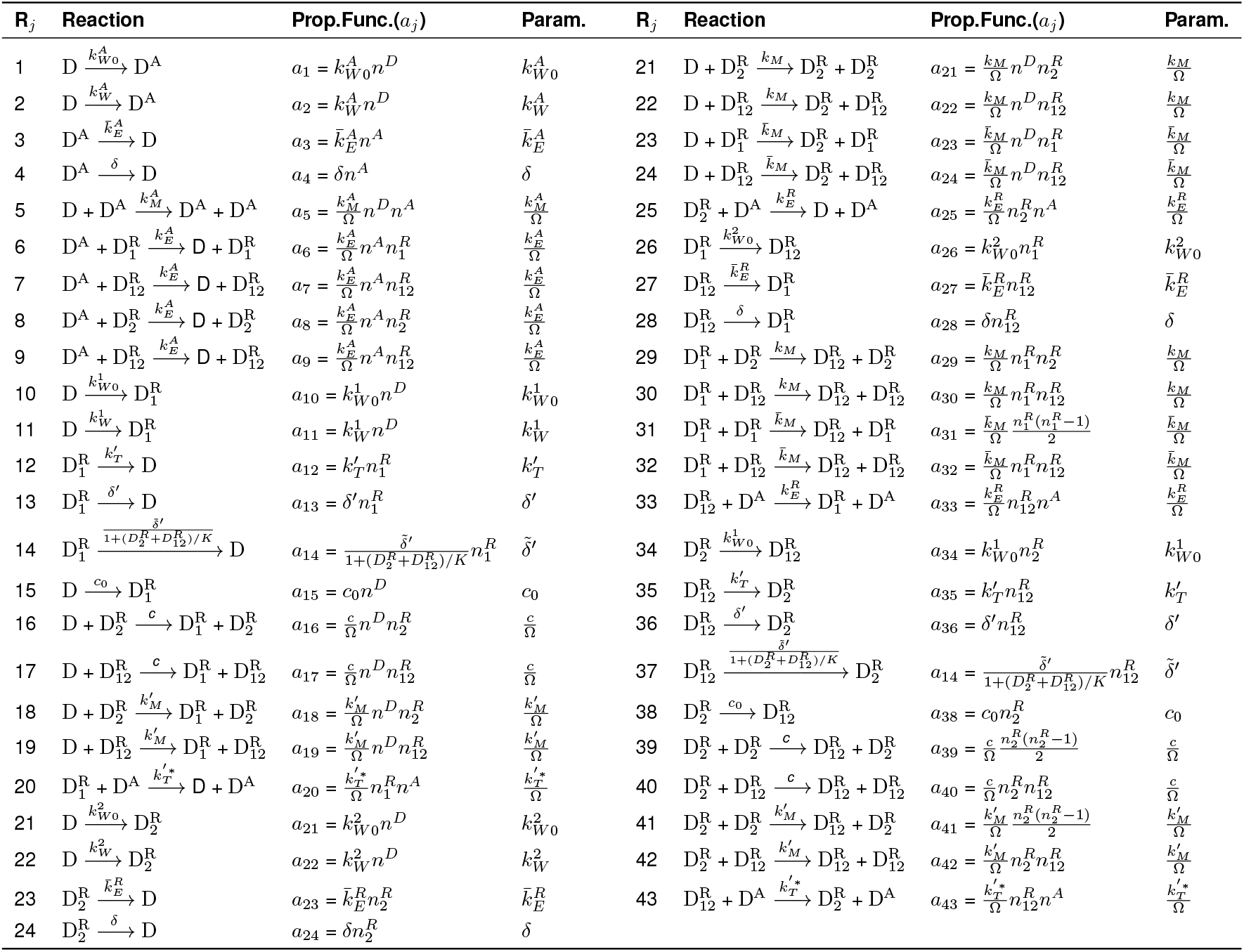
Full chromatin modification circuit model incorporating the refined DNA methylation maintenance model described in SI Section S.5: reactions.

### S.7 Modification to the chromatin modification circuit model to include more than one CpG per nucleosome

In the original model of the chromatin modification circuit, it is assumed that the basic unit, that is a nucleosome with DNA wrapped around, can have only one modifiable CpG [66]. There are cases in which this assumption is not verified. However, it is possible to show that this assumption does not affect the qualitative results related to the effect of the parameters *α*′ (normalized rate of DNA methylation establishment through repressive histone modifications) and *µ*′ (ratio between the DNA demethylation rate and the activating histone modification erasure rate) on the probability distribution of gene expression levels.

To this end, let us assume that the basic unit of the model, D, is the nucleosome with DNA wrapped around it with two modifiable CpGs. The conclusion obtained in this section would not change if we consider the DNA wrapped around the nucleosome having *n* ∈ ℤ^+^ modifiable CpGs. The basic unit of the model can then be modified with H3K4me3, D^A^, one DNA methylation, 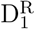, H3K9me3, 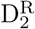, one DNA methylation and H3K9me3, 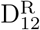, two DNA methylations, 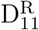, and two DNA methylation and H3K9me3,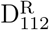. Now, defining the number of 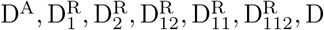, D as 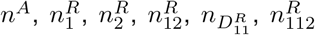 and *n*^*D*^, let us derive the related ordinary differential equation (ODE) model in terms of the fractions 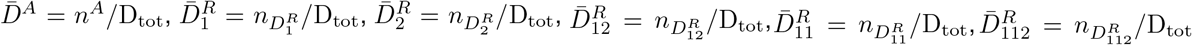,and 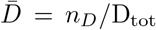, with 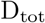 the total number of nucleosomes on a gene of interest. This can be done by assuming that D_tot_ is sufficiently large so that 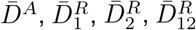 and 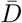 can be treated as real number. Now, considering the dimensionless parameters and normalized inputs introduced SI Section S.1 and the normalized time 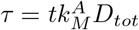, the ODEs associated with the chromatin modification circuit are

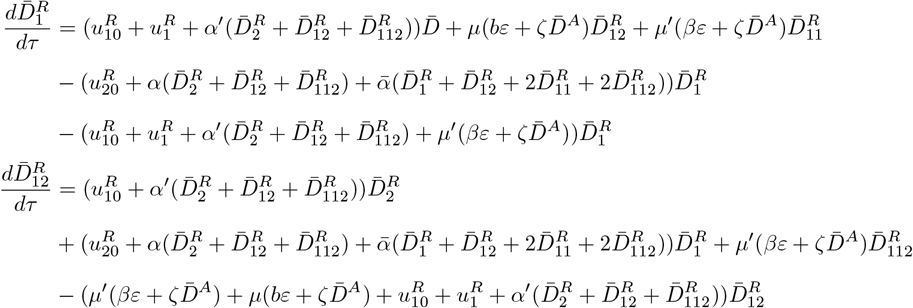

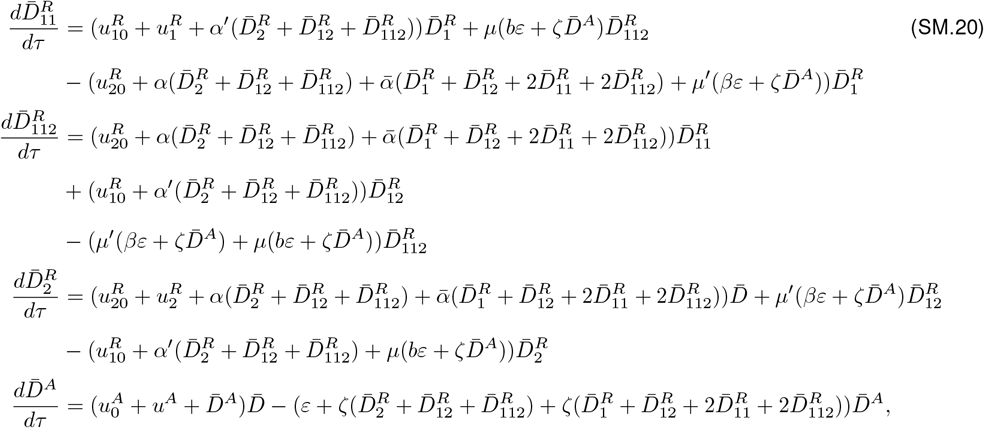

with 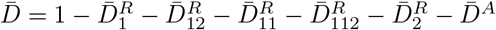.

As we did for the original ODE system (SM.3), let us merge the rates linked to the enhancement of H3K9me3 establishment by 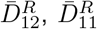, and 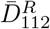 into a single rate. We will assume that this rate is identical to the one associated with the enhancement of H3K9me3 establishment by 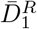. Similarly, let us merge the rates linked to the recruited erasure of H3K4me3 by 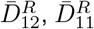, and 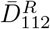into a single rate and assume that this rate is identical to the one associated with the recruited erasure of H3K4me3 by 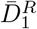. The ODE system (SM.20) can then be rewritten as

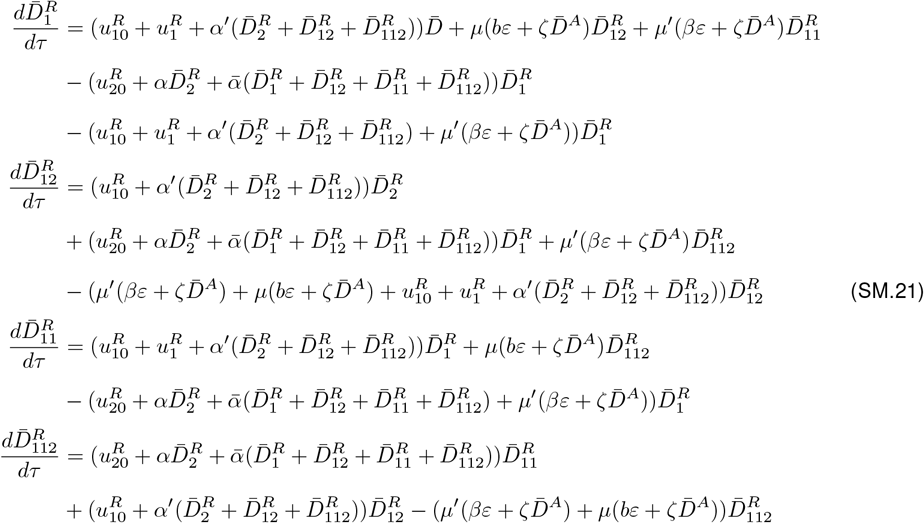

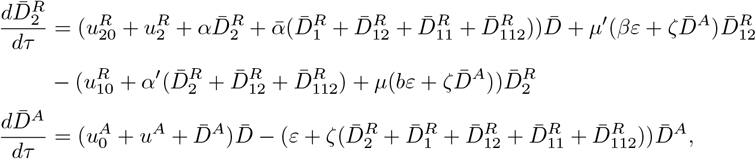

Now, let us rewrite system (SM.21) by assuming negligible basal establishment 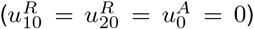 and introducing the variable 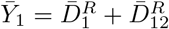, which corresponds to the fraction of nucleosomes with one methylated CpG, and the variable 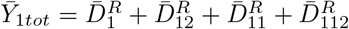, which corresponds to the fraction of nucleosomes with at least one methylated CpG. We then obtain

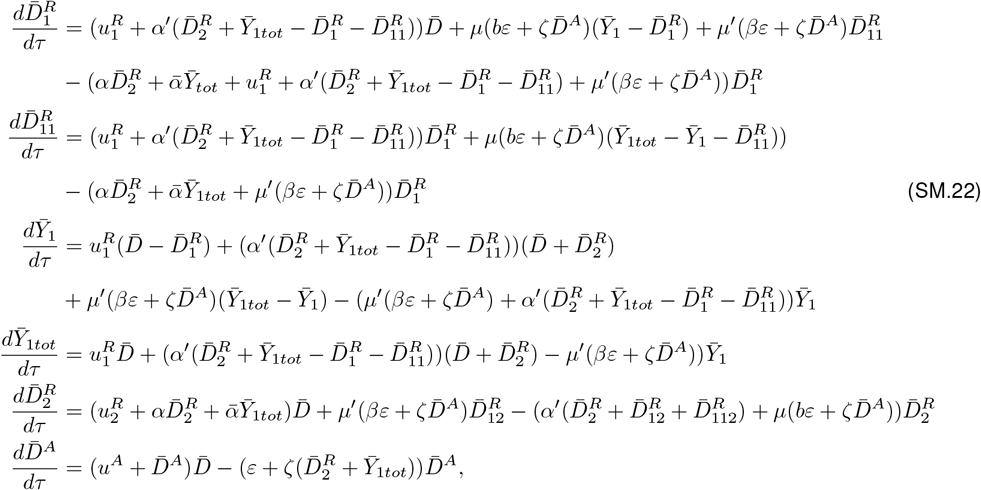

with 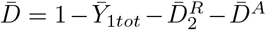. As we did for the ODE system associated with our original chromatin modification circuit in SI Section S.1, we can further simplify our model by assuming that the enhancement of DNA methylation establishment by H3K9me3 is negligible, i.e., *α*′ = 0. The ODE system (SM.22) can then be rewritten as

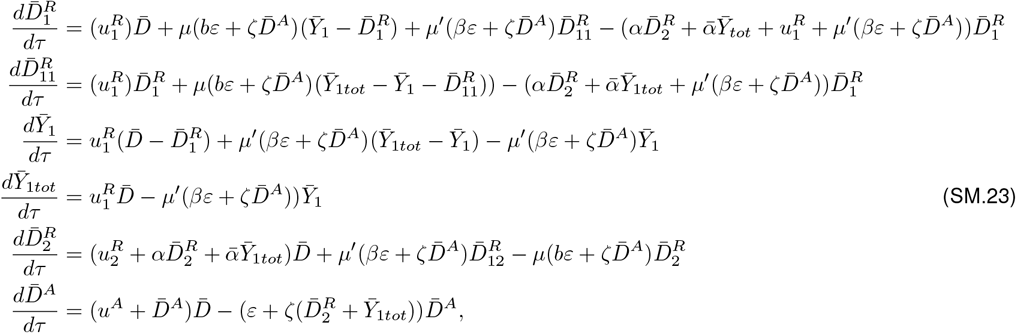

with 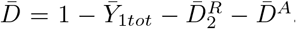. Now, introducing in (SM.23) the expressions for 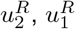, and *µ*′ derived in SI Section S.1.1 (Expressions (SM.5), (SM.7), and (SM.9)), we obtain:

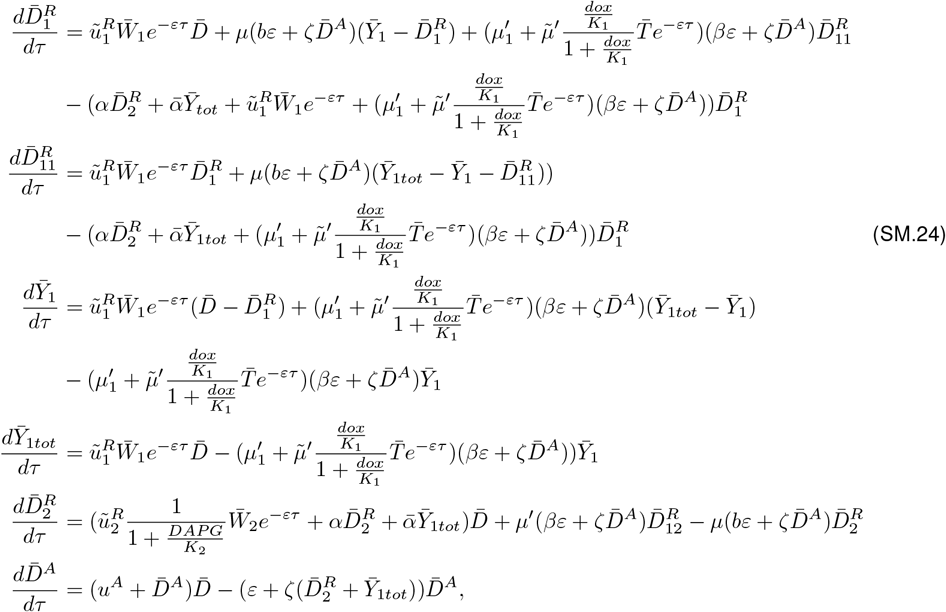

with 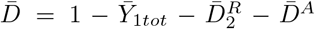. Furthermore, if we consider the parameter regime in which the rate of DNA demethylation, without transfection of rTetR-TET1, is sufficiently low compared to histone modification dynamics,i.e., 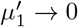, the system (SM.24) can then be rewritten as follows:

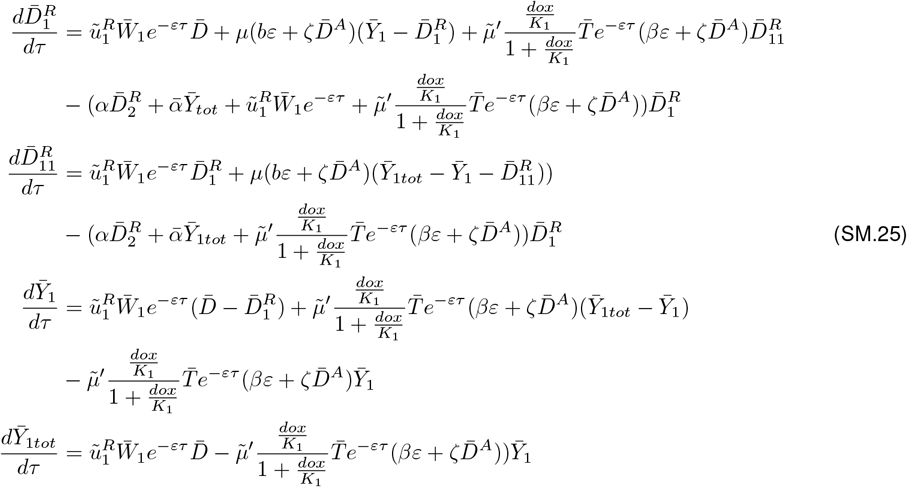

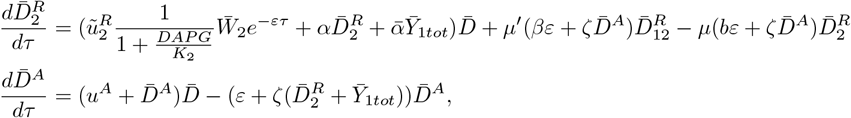

with 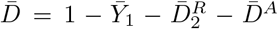. From the fourth equation in (SM.25), it is possible to notice that, following a temporary phase during which the externally transfected inputs decrease, *e*^−*ετ*^ ≈ 0, and then 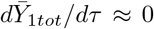. This implies that, once the transient phase concludes, 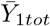 stabilizes at a constant value and the behavior of the original model can be captured by a reduced 2D model described by the subsequent ODEs:

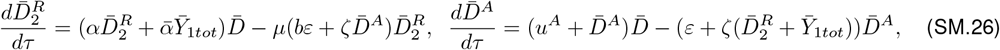

with 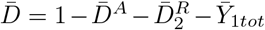 and 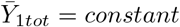. It is evident that the ODE model (SM.26) closely resembles the original model, in which we assume to have one CpG per nucleosome, i.e., (SM.11), with the only distinction being the presence of 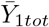 instead of 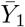. That is, in this revised model, the representation of the fraction of the number of methylated CpGs in the gene is denoted by 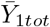. Then, the introduction of the simplifying assumption, having one CpG per nucleosome, does not affect the qualitative results of the analysis that focuses on identifying the impact of the parameters *α*′ (normalized rate of DNA methylation establishment through repressive histone modifications) and *µ*′ (ratio between the DNA demethylation rate and the activating histone modification erasure rate) on the probability distribution of gene expression levels.

### S.8 Supplementary Modeling Figures

**Fig. SM.1.**
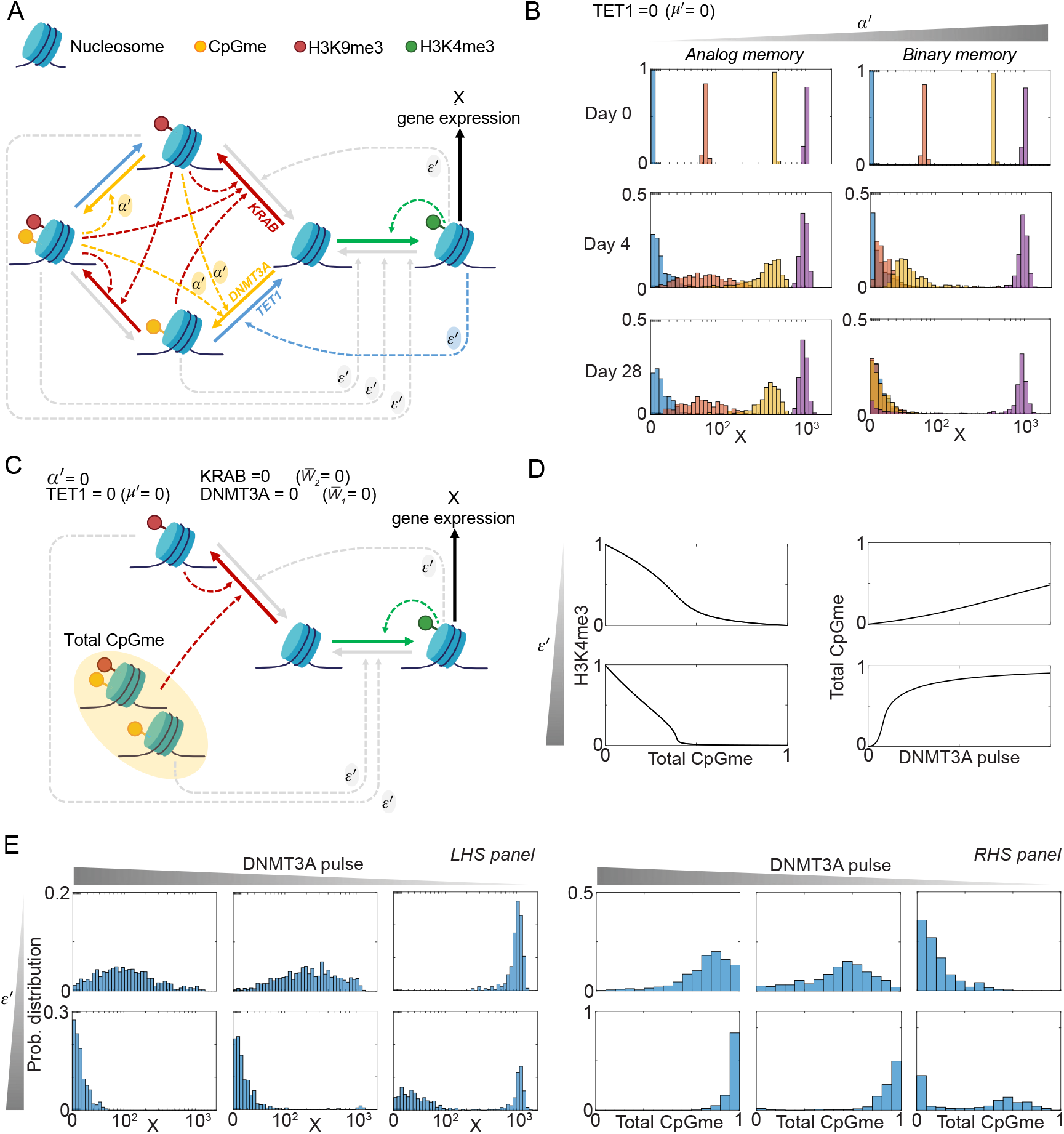
The dynamics of the chromatin modification circuit can be captured by a reduced model, under specific parameter regimes. **A** Full chromatin modification circuit diagram. The species involved are unmodified nucleosome, nucleosome with H3K4me3, nucleosome with H3K9me3, nucleosome with CpGme, and nucleosome with both H3K9me3 and CpGme. We use solid arrows to represent the nucleosome modifications and gene expression. We use dashed arrows to represent the recruitment process and the consequent effect on establishment or erasure. For each arrow, we use red for processes involved in H3K9me3 establishment, orange for processes involved in DNA methylation establishment, green for processes involved in H3K4me3 establishment, light blue for processes involved in DNA demethylation, gray for processes involved in the erasure of histone modifications, and black for gene expression. In particular, *α*_′_ = 0 when H3K9me3 does not recruit writers of DNA methylation. In order to simplify the diagram without compromising the essential interactions among modifications, we removed some non-critical dashed arrows (refer to Fig. 1 in [66] for the complete diagram). **B** Probability distributions of the system represented by reactions listed in SI Tables SM.1 and SM.4. The distributions are obtained computationally using SSA [52] and we indicate with X the gene expression level (logicle scale). In particular, we determine how *α*_′_ affects the probability distribution of the system when *µ*_′_ = 0. The parameter values used for these simulations are listed in SI Section S.9. In particular, we consider *α*_′_ = 0, 0.1 and four initial conditions (H3K9me3 and CpGme, CpGme only, H3K9me3 only, H3K4me3)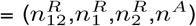: (14,0,1,0) (blue), (6,0,8,1) (red), (4,0,5,6) (yellow), and (1,0,1,13) (purple). For each case, the initial value of X at time *t* = 0 was set to its steady state. For all the simulations we set 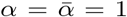, *ε* = 0.08, *ζ* = 25, *µ* = 0.1, *b* = *β* = 1. See SI Fig. SM.2 for how different values of *α*_′_ and *µ*_′_ influence the probability distribution of the system. **C** Simplified chromatin modification circuit diagram obtained when *α*_′_ = 0, TET1 = 0 (i.e., *µ*′ = 0), DNMT3A = 0 (i.e., 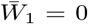), KRAB = 0 (i.e., 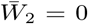). **D** Left-hand side plots: dose-response curve for the (Total CpGme,H3K4me3) pair, i.e, 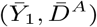, for different values of *ζ* obtained from simulations of system (SM.11) with (Total CpGme,H3K9me3,H3K4me3) = 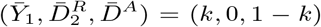 as initial conditions, with 0 ≤ *k* ≤ 1. Right hand-side plots: dose-response curve for the (DNMT3A,Total CpGme) pair, i.e, 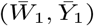, for different values of *ζ* obtained from simulations of system (SM.16) with (Total CpGme,H3K9me3,H3K4me3) 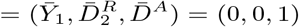 as initial conditions. The parameter values used for these simulations are listed in SI Section S.9. In particular, we consider *ε* = 0.1 and *ζ* = 1, 25 (See SI Fig. SM.3 for how different values of *ε* and *ζ* influence the deterministic behavior of the system). Here, the DNMT3A dynamics is modeled as a pulse that exponentially decreases over time. The value on the x-axis corresponds to the DNMT3A value at time 0. **E** Probability distributions of the system, represented by reactions listed in SI Tables SM.1 and SM.4, after t = 28 days. The distributions are obtained computationally using SSA [52]. In the left-hand side panel we show the probability distributions of the gene expression level X (logicle scale) and in the right-hand side panel we show the probability distributions of the total CpGme level. The parameter values used for these simulations are listed in SI Section S.9. In particular, we consider *ε* = 0.13, *ζ* = 1.5, 25, 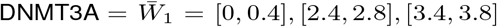 and (H3K9me3 and CpGme, CpGme only, H3K9me3 only, H3K4me3) 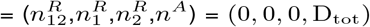 as initial condition (see SI Fig. SM.3 for how different values of *ε* and *ζ* influence the probability distribution of the system). In our model, *α*′ is the normalized rate of DNA methylation establishment through repressive histone modifications, *µ*′ is the ratio between the DNA demethylation rate and the activating histone modification erasure rate, and *ε* (*ζ*) is the parameter that scales the ratio between the basal (recruited) erasure rate and the auto-catalysis rate of each modification. For panels B and E, we consider D_tot_ = 15 and we consider *N* = 1000 samples to realize each distribution and, for each simulation, the value of 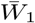 was randomly selected from a uniformly distributed range, whose extremes are listed above.

**Fig. SM.2.**
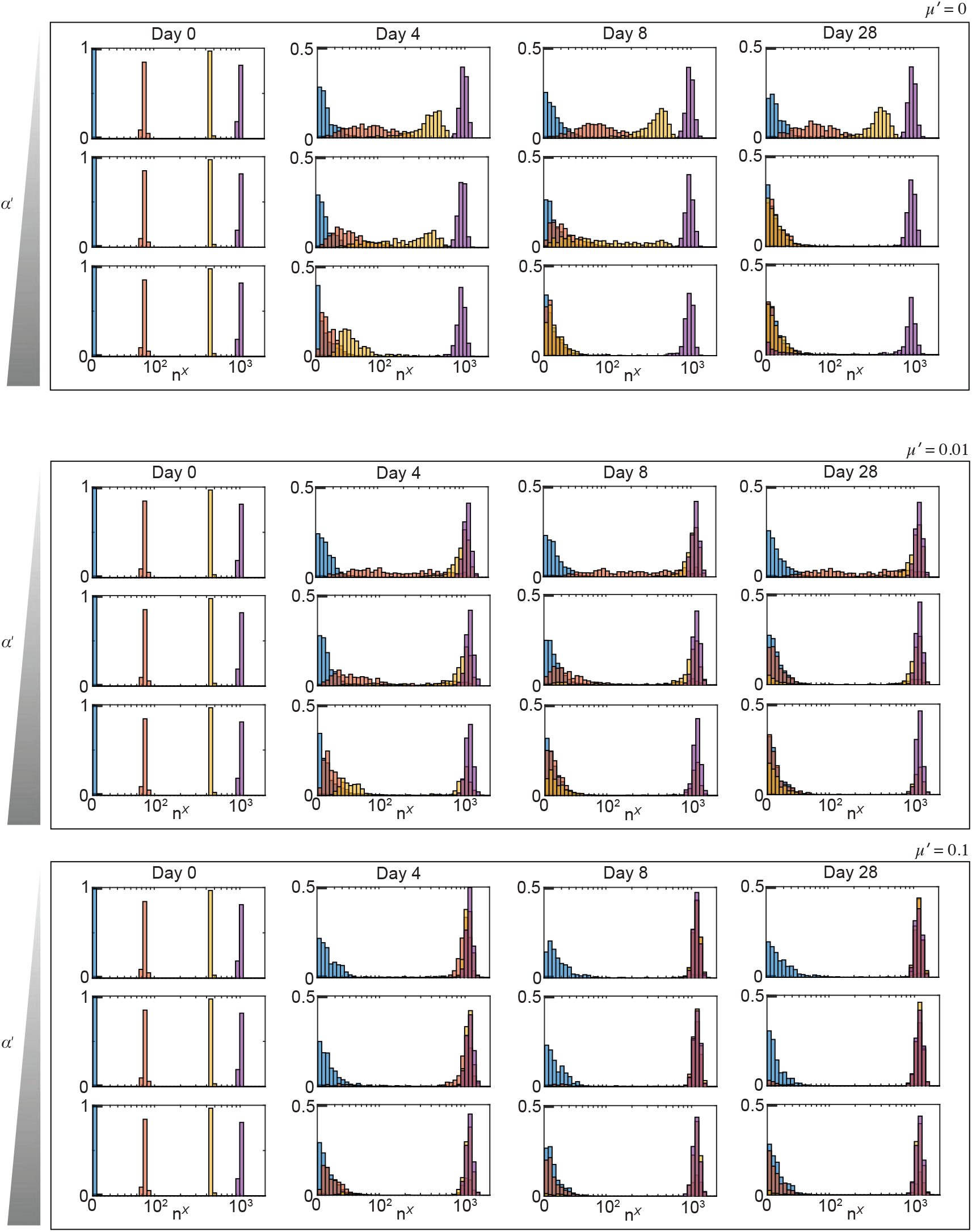
Achieving analog memory is only possible when *α*_′_ = 0 and *µ*_′_ = 0. Probability distributions of the system represented by reactions listed in SI Tables SM.1 and SM.4. The distributions are obtained computationally using SSA [52] and we indicate with *n*^*X*^ the gene expression level (logicle scale). In particular, we determine how *α*′ and *µ*′ affect the probability distribution of the system. The parameter values used for these simulations are listed in SI Section S.9. In particular, we consider *α*′ = 0, 0.01, 0.1 and *µ*′ = 0, 0.01, 0.1 and four initial conditions: 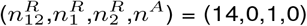 (blue), 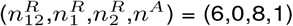(red), 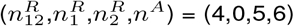 (yellow),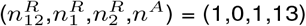 (purple). For each case, the initial values of *n*^*X*^ of *n*^*m*^ at time *t* = 0 were set to their steady states. For all the simulations we set 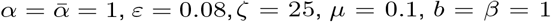. In our model, *α*′ is the normalized rate of DNA methylation establishment through repressive histone modifications and *µ*′ is the ratio between the DNA demethylation rate and the activating histone modification erasure rate. For all simulations, we consider D_tot_ = 15 and we consider *N* = 1000 samples to realize each distribution.

**Fig. SM.3.**
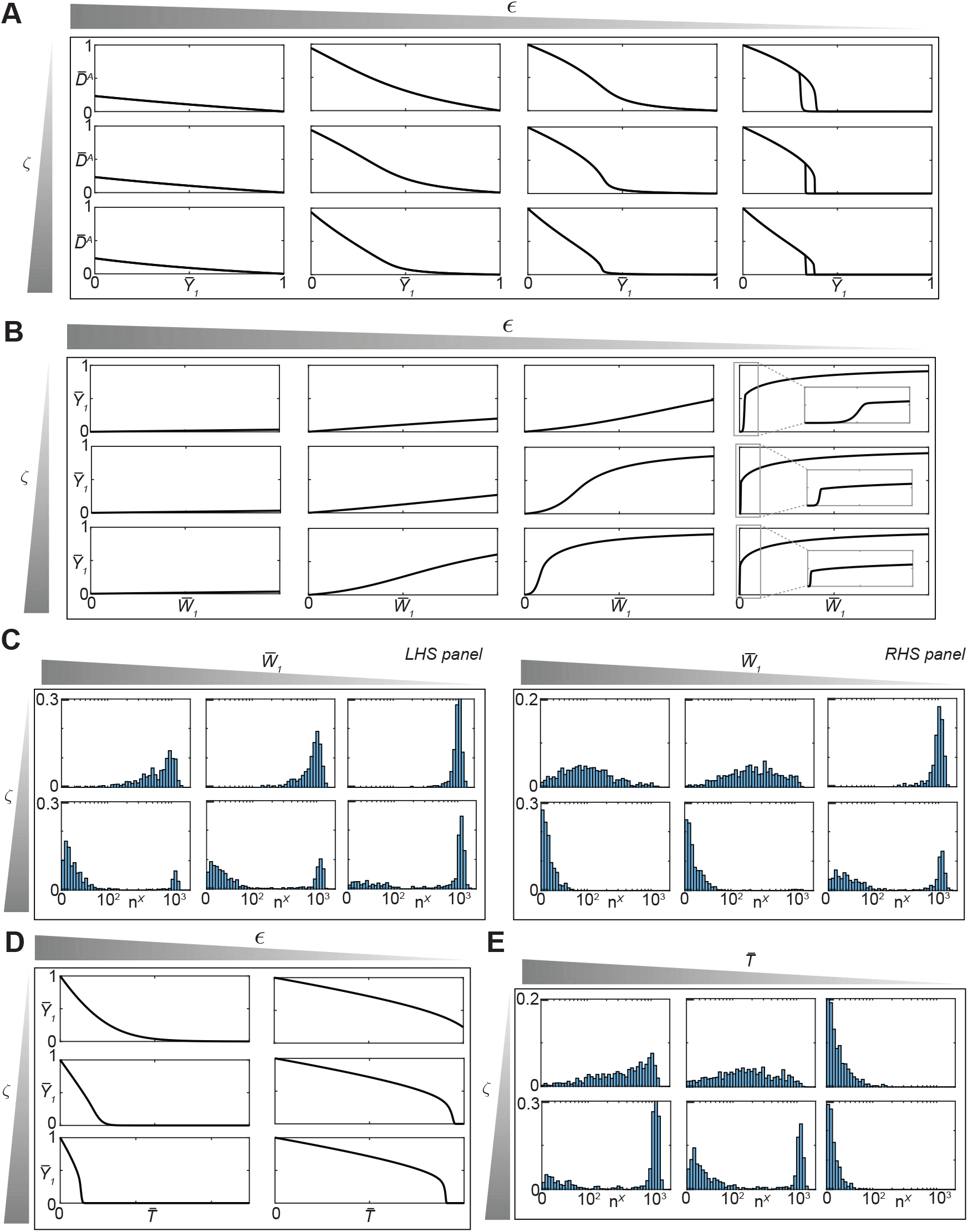
The parameters *ε* and *ζ* affect the shape of the probability distribution for gene expression levels. **A** Input/output steady state characteristics for the 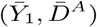 pair for different values of *ε* and *ζ* obtained from simulations of system (SM.11) with 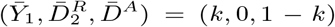 as initial conditions, with 0 ≤ *k* ≤ 1. The parameter values used for these simulations are listed in SI Section S.9. In particular, *ε* = 0.0001, 0.1, 1, 50 and *ζ* = 1, 5, 25. **B** dose-response curve for the 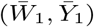 pair for different values of *ε* and *ζ* obtained from simulations of system (SM.16) with 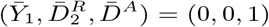 as initial conditions. The parameter values used for these simulations are listed in SI Section S.9. In particular, *ε* = 0.0001, 0.1, 1, 50 and *ζ* = 1, 5, 25. **C** Probability distributions of the system represented by reactions listed in SI Tables SM.1 and SM.4 after t = 28 days. The distributions are obtained computationally using SSA [52] and we indicate with *n*^*X*^ the gene expression level (logicle scale). The parameter values used for these simulations are listed in SI Section S.9. In particular, in the left-hand side panel we consider 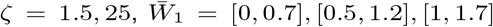 and 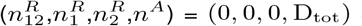 as initial conditions. The initial values of *n*^*X*^ of *n*^*m*^ at time *t* = 0 were set to their steady states of the ODEs. In the right-hand side panel, we consider 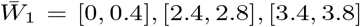 and all the other parameter values as the ones used for the simulations in the left-hand side panel. For all the simulations we set 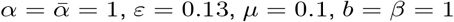, and we consider D_tot_ = 15 and *N* = 1000 samples to realize each distribution. For each simulation, the value of 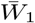 was randomly selected from a uniformly distributed range, whose extremes are listed above. **D** Dose-response curve for the 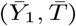 pair for different values of *ε* and *ζ* obtained from simulations of system (SM.16) with 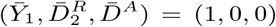 as initial conditions. The parameter values used for these simulations are listed in SI Section S.9. In particular, *ε* = 0.0001, 0.1 and *ζ* = 1, 5, 25. **E** Probability distributions of the system represented by reactions listed in SI Tables SM.1 and SM.4 after t = 28 days. The distributions are obtained computationally using SSA [52] and we indicate with *n*^*X*^ the gene expression level (logicle scale). The parameter values used for these simulations are listed in SI Section S.9. In particular, we consider 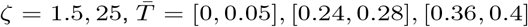 and 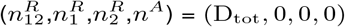 as initial conditions. The initial values of *n*^*X*^ of *n*^*m*^ at time *t* = 0 were set to their steady states of the ODEs. For all the simulations we set 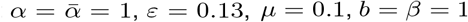, and we consider D_tot_ = 15 and *N* = 1000 samples to realize each distribution. For each simulation, the value of 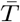was randomly selected from a uniformly distributed range, whose extremes are listed above. In our model, *ε* (*ζ*) is the parameter that scales the ratio between the basal (recruited) erasure rate and the auto-catalysis rate of each modification and 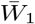 and 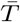 represent the normalized amounts of DNMT3A and TET1 at time 0, which exponentially decrease over time.

**Fig. SM.4.**
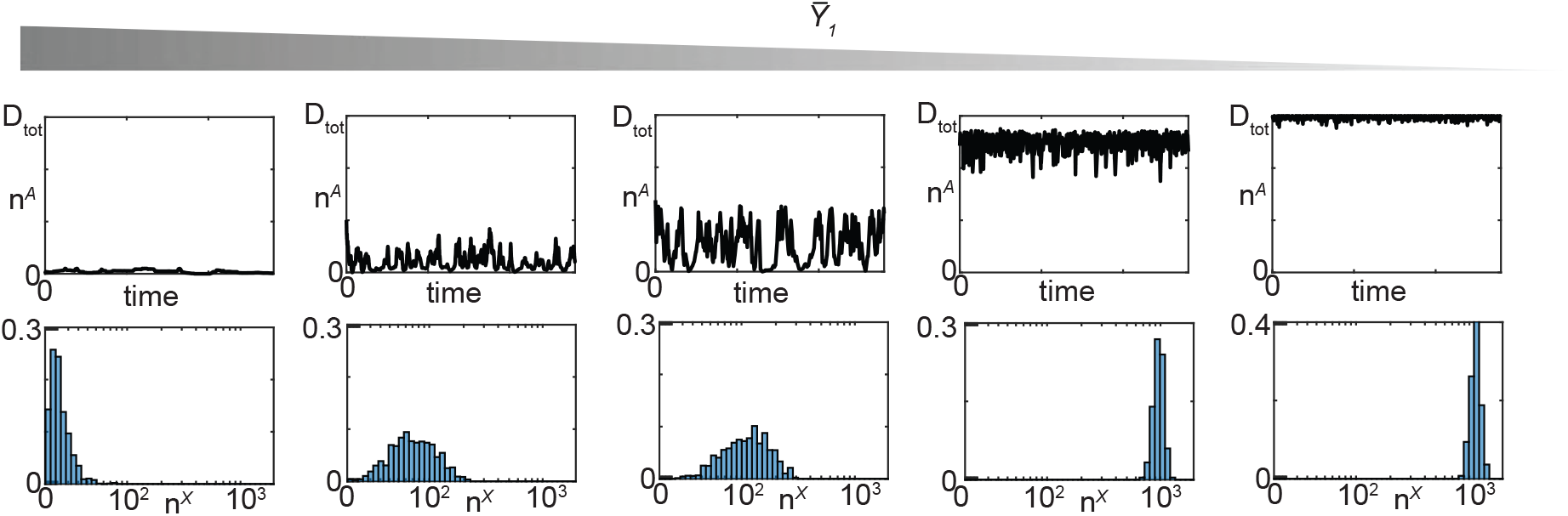
When *ε* is sufficiently small, intermediate DNA methylation grades results in broader gene expression distributions. **A** Time trajectories of n^*A*^ for different values of 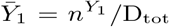 obtained by simulating reactions listed in SI Table SM.3. The trajectories are obtained computationally using SSA [52]. The parameter values used for these simulations are listed in SI Section S.9. In particular, we consider 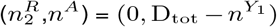 as initial conditions **B** Probability distributions of the system represented by reactions listed in SI Tables SM.3 and SM.4 after t = 28 days. The distributions are obtained computationally using SSA [52] and we indicate with *n*^*X*^ the gene expression level (logicle scale). The parameter values used for these simulations are listed in SI Section S.9. In particular, we consider 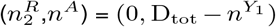 as initial conditions, and *N* = 1000 samples to realize each distribution. The initial values of *n*^*X*^ of *n*^*m*^ at time *t* = 0 were set to their steady states of the ODEs. For all the simulations in both panels A and B, we set 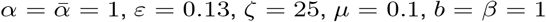, and we consider D_tot_ = 15 and *n*^*Y*1^ = 0, 1, 6, 8, 14. In our model, *ε* is the parameter that scales the ratio between the basal erasure rate and the auto-catalysis rate of each modification.

**Fig. SM.5.**
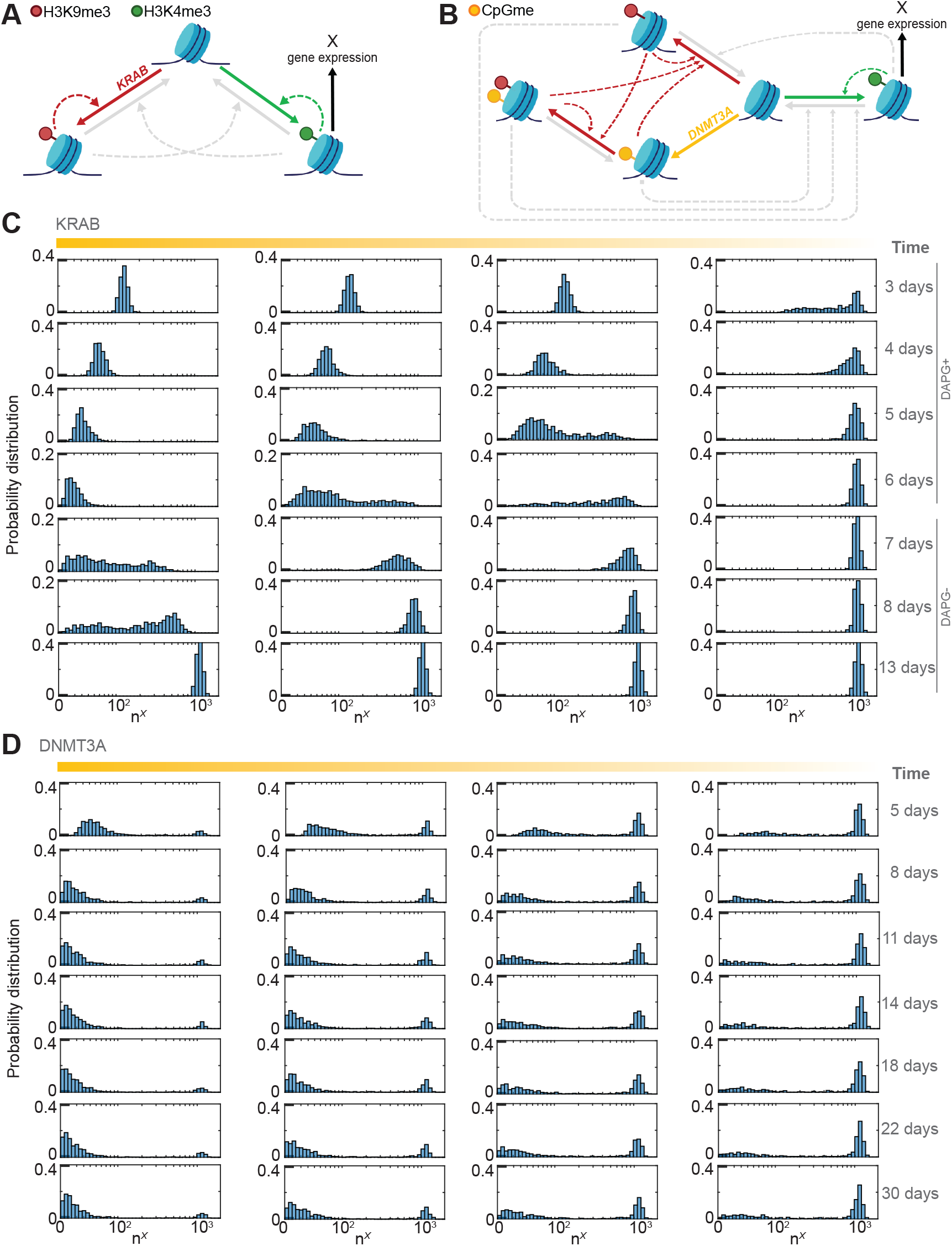
Simulations recapitulating the effect of transient transfection of KRAB and DNMT3A on the gene expression probability distribution. **A** Diagram of the simplified chromatin modification circuit, including H3K9me3 and H3K4me3 only. We use solid arrows to represent the nucleosome modifications and gene expression. We use dashed arrows to represent the recruitment process and the consequent effect on establishment or erasure. For each arrow, we use red for processes involved in H3K9me3 establishment, green for processes involved in H3K4me3 establishment, gray for processes involved in the erasure of histone modifications, and black for gene expression. **B** Diagram of the chromatin modification circuit including H3K9me3, H3K4me3, and CpGme. We use solid arrows to represent the nucleosome modifications and gene expression. We use dashed arrows to represent the recruitment process and the consequent effect on establishment or erasure. For each arrow, we use red for processes involved in the H3K9me3 establishment, green for processes involved in H3K4me3 establishment, orange for processes involved in DNA methylation establishment, gray for processes involved in the erasure of histone modifications, and black for gene expression. **C** Probability distributions of the system represented in panel A, whose reactions are listed in SI Tables SM.2 and SM.5. The distributions are obtained computationally using SSA [52] and we indicate with *n*^*X*^ the gene expression level (logicle scale). The parameter values used for these simulations are listed in SI Section S.9. In particular, we consider 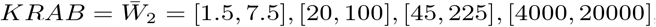, and 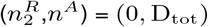 as initial conditions. The initial values of *n*^*X*^ of *n*^*m*^ at time *t* = 0 were set to their steady states of the ODEs. **D** Probability distributions of the system represented in panel B, whose reactions are listed in SI Tables SM.1 and SM.4. The distributions are obtained computationally using SSA [52] and we indicate with *n*^*X*^ the gene expression level (logicle scale). The parameter values used for these simulations are listed in SI Section S.9. In particular, we consider 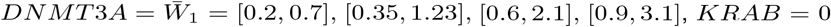, and 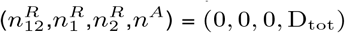 as initial conditions. The initial values of *n*^*X*^ of *n*^*m*^ at time *t* = 0 were set to their steady states of the ODEs. For all the simulations we set *α* = 1, *ε* = 0.13, *ζ* = 25, *µ* = 0.1, *b* = 1, we model the dynamics of KRAB and DNMT3A as pulses that exponentially decrease over time, and we consider D_tot_ = 15 and *N* = 1000 samples to realize each distribution. For each simulation, the value of 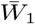 (or 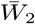) was randomly selected from a uniformly distributed range, whose extremes are listed above.

**Fig. SM.6.**
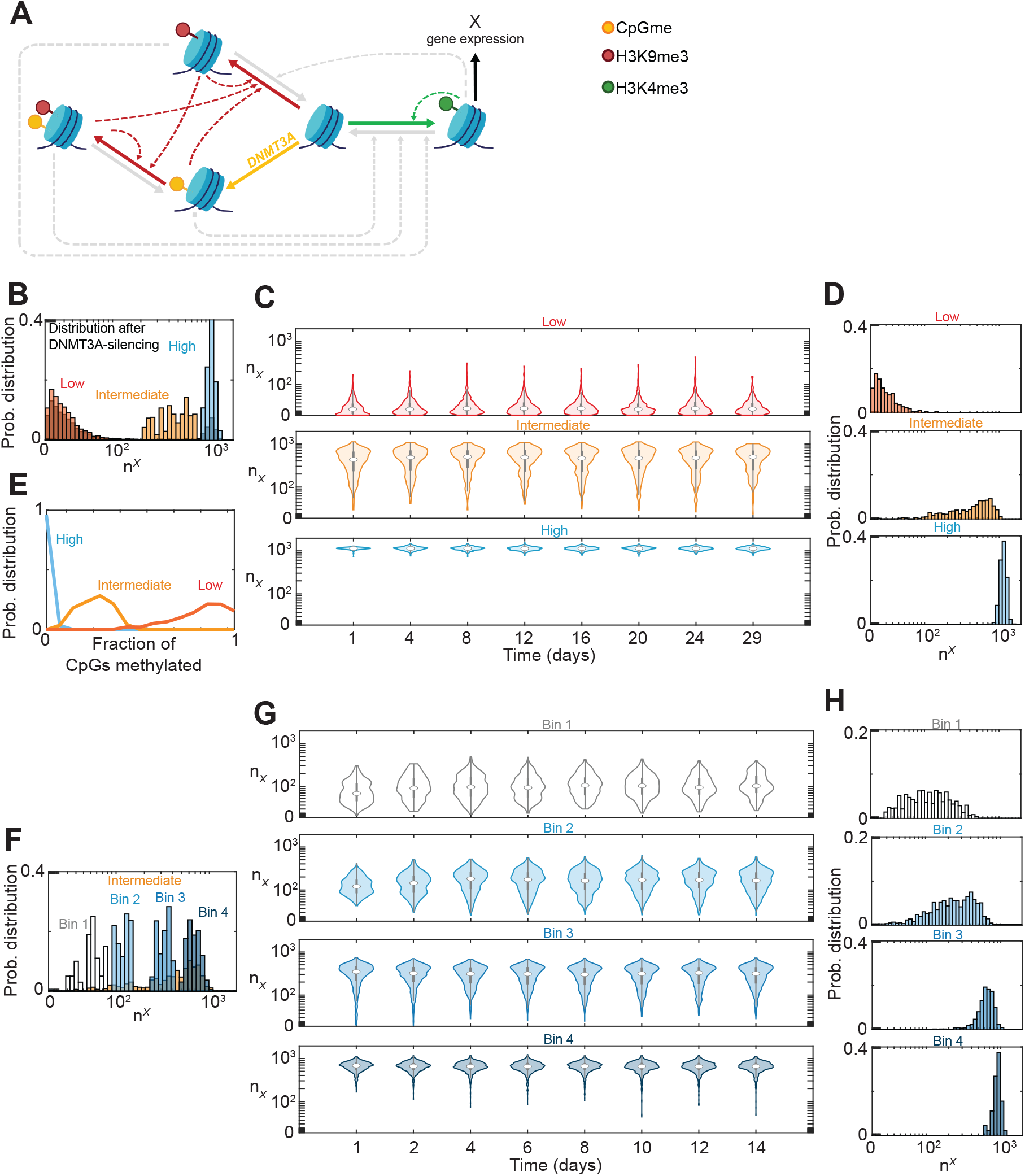
Simulations recapitulating the experimental results shown in Fig. 3, showing that intermediate gene expression levels are stable and exhibit intermediate DNA methylation levels. **A** Diagram of the chromatin modification circuit including H3K9me3, H3K4me3, and CpGme. We use solid arrows to represent the nucleosome modifications and gene expression. We use dashed arrows to represent the recruitment process and the consequent effect on establishment or erasure. For each arrow, we use red for processes involved in H3K9me3 establishment, green for processes involved in H3K4me3 establishment, orange for processes involved in DNA methylation establishment, gray for processes involved in the erasure of histone modifications, and black for gene expression. **B** Probability distributions of the system represented in panel A, whose reactions are listed in SI Tables SM.1 and SM.4. The distributions are obtained computationally using SSA [52] and we indicate with *n*^*X*^ the gene expression level (logicle scale). The black distribution represents the probability distribution of the system at day 30 after DNMT3A-silencing. The parameter values used for these simulations are listed in SI Section S.9. In particular, we consider 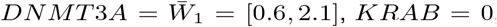, and 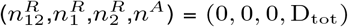 as initial conditions. The initial values of *n*^*X*^ of *n*^*m*^ at time *t* = 0 were set to their steady states of the ODEs. The low (red), intermediate (orange), and high (blue) distributions are obtained by subdividing the black distribution with respect to the following intervals of *n*^*X*^ :[0,100], [200,800], [900,10000]. **C** Probability distributions of the system represented in panel A obtained as described in panel B. The parameter values used for these simulations are listed in SI Section S.9. In particular, for each case, we consider *DNMT* 3*A* = 0, *KRAB* = 0, and, as initial conditions, points from the red, orange, and blue distributions shown in panel B, respectively. **D** Probability distributions of the system at day 29 from panel C. **E** Probability distribution of the fraction of CpGs methylated, i.e., 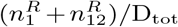, at day 29 obtained from the simulations conducted to obtain panel C. **F** Probability distributions of the system represented in panel A, whose reactions are listed in SI Tables SM.1 and SM.4. The distributions are obtained computationally using SSA [52] and we indicate with *n*^*X*^ the gene expression level (logicle scale). The orange distribution is the one obtained and described in panel D. The other ones are obtained by subdividing the orange distribution with respect to the following intervals of *n*^*X*^ : [0,80] (Bin 1), [90,150] (Bin 2), [250,400] Bin 3, [500,10000] (Bin 4). **G** Probability distributions of the system represented in panel A, obtained as described in panel F. The parameter values used for these simulations are listed in SI Section S.9. In particular, for each case, we consider *DNMT* 3*A* = 0, *KRAB* = 0, and, as initial conditions, points from the “Bin 1”, “Bin 2”, “Bin 3”, and “Bin 4” distributions shown in panel F, respectively. **H** Probability distributions of the system at day 14 from panel G.

**Fig. SM.7.**
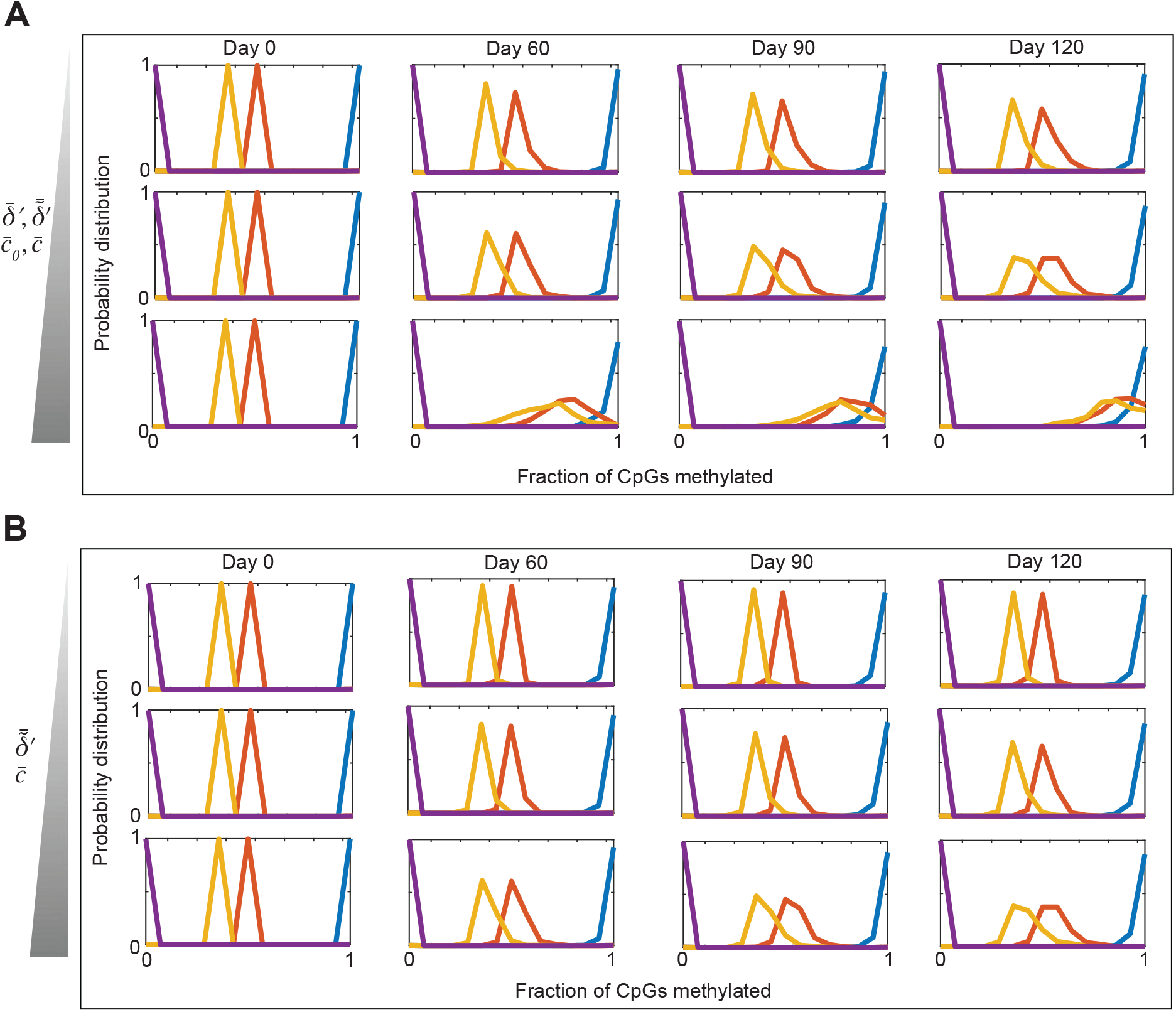
Stochastic analysis of the full chromatin modification circuit model including the refined DNA methylation maintenance model. **A** Probability distributions of the system represented by reactions listed in SI Tables SM.6 and SM.4. The distributions are obtained computationally using SSA [52] and, on the *x* we have the fraction of CpGs methylated. In particular, we determine how lower parameter values affect the probability distribution of the system. The parameter values used for these simulations are listed in SI Section S.9. In particular, we consider 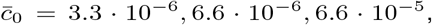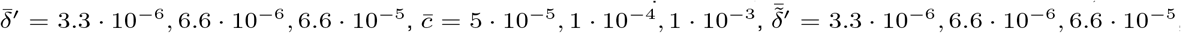, and four initial fractions of CpGs methylated, i.e., (0.93,0.4,0.27,0). Specifically, as initial conditions, we set 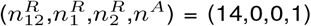 (blue), 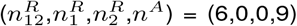 (red), 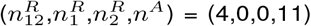 (yellow), 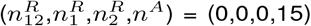 (purple). For each case, The initial values of *n*^*X*^ of *n*^*m*^ at time *t* = 0 were set to their steady states of the ODEs. For all the simulations we set 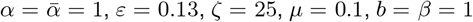. **B** Probability distributions of the system represented by reactions listed in SI Tables SM.6 and SM.4. The distributions are obtained computationally using SSA [52] and, on the *x* we have the fraction of CpGs methylated. In particular, we determine how 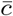 and 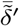, i.e., normalized rate constants associated with the reactions representing the effect of DNMT1 recruitment by H3K9me3 on the maintenance process, affect the probability distribution of the system. The parameter values used for these simulations are listed in SI Section S.9. In particular, we consider 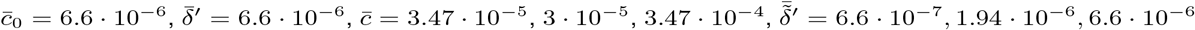.The values of the other parameters and the initial conditions are the same as those used for panel A. For all simulations, we consider D_tot_ = 15 and we consider *N* = 1000 samples to realize each distribution.

**Fig. SM.8.**
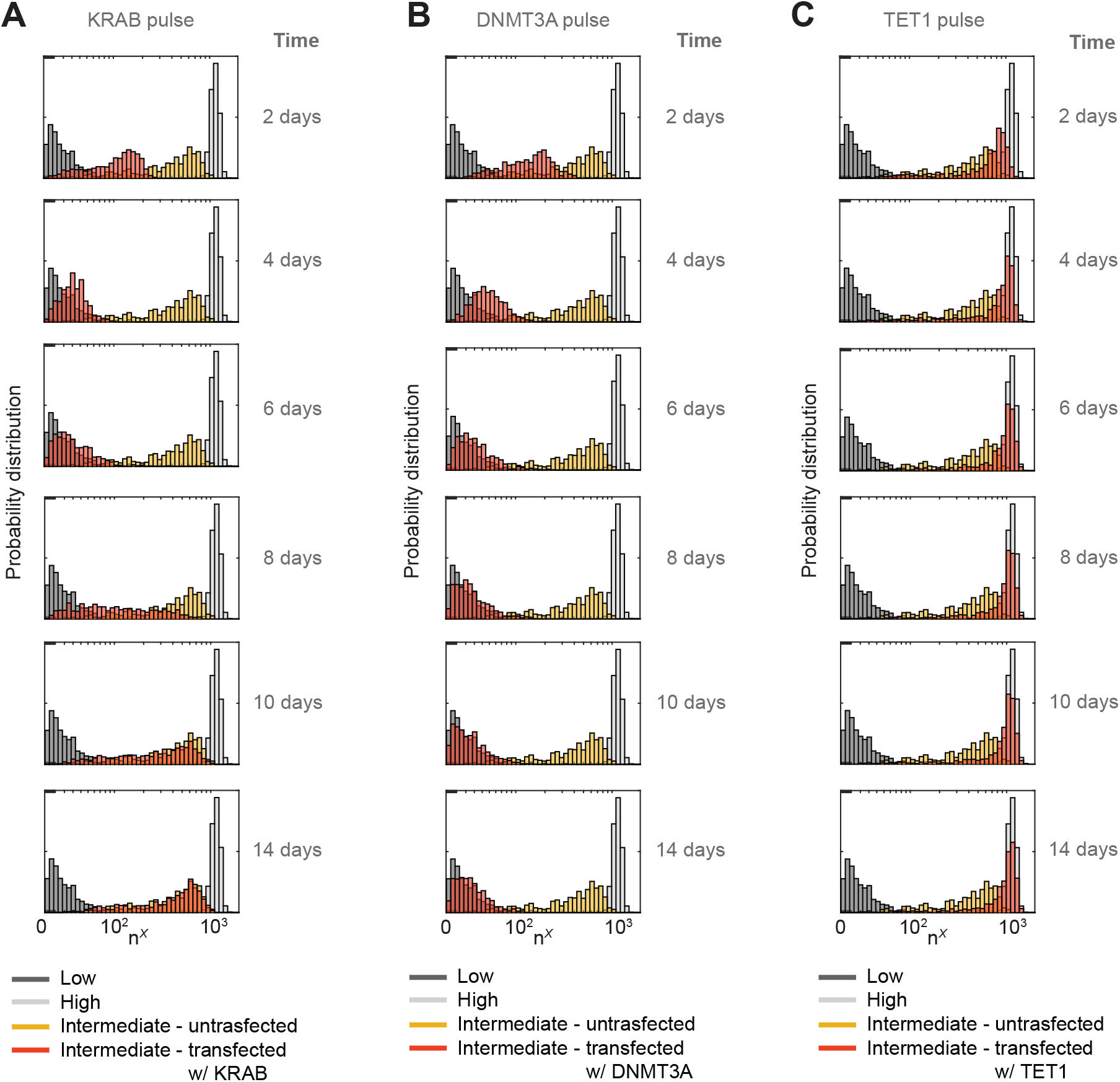
Simulations recapitulating the effect of KRAB, DNMT3A, and TET1 on the gene expression probability distribution, starting from an intermediate level of gene expression. **A** Probability distributions of the system represented in Fig. SM.5A, whose reactions are listed in SI Tables SM.2 and SM.5. The distributions are obtained computationally using SSA [52] and we indicate with *n*^*X*^ the gene expression level (logicle scale). The parameter values used for these simulations are listed in SI Section S.9. In particular, we consider 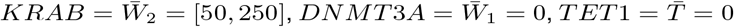. As initial conditions, we consider values of 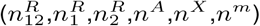 from the simulations associated with the orange distribution in SI Fig. SM.6D. **B** Probability distributions of the system represented in Fig. SM.5B, whose reactions are listed in SI Tables SM.1 and SM.4. The distributions are obtained computationally using SSA [52]. The parameter values used for these simulations are listed in SI Section S.9. In particular, we consider 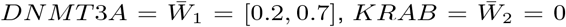, and 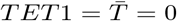. **C** Probability distributions of the system represented in Fig. SM.5B, whose reactions are listed in SI Tables SM.1 and SM.4. The parameter values used for these simulations are listed in SI Section S.9. In particular, we consider 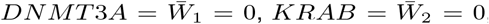, and 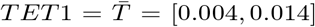. For all the simulations we set 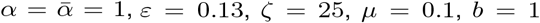, we model the dynamics of KRAB, DNMT3A, and TET1 as pulses that exponentially decrease over time, and we consider D_tot_ = 15 and *N* = 1000 samples to realize each distribution. For each simulation, the value of 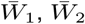 or 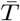 was randomly selected from a uniformly distributed range, whose extremes are listed above.

### S.9 Values of parameters and rationale for parameter selections used in generating figures in the main paper and SI

In our model, we defined the total number of nucleosomes within a gene of interest as D_tot_. Assuming approximately one nucleosome per 200 bp [71] and given that the length of our gene is ≈ 3000bp, then in our computational study we set the value of D_tot_ to 15.

With regard to the effect of dilution due to cell growth, we use a standard approach by modeling it as a first-order decay reaction [72]. The decay rate constant can then be expressed as *δ* = *ln*(2)*/T*, where *T* corresponds to the cell cycle length. Given that in our experiments we use CHO cells, whose cycle length has been estimated to be ≈ 20 hr [23], in the simulations aimed at replicating our experimental data, we set *δ* = 0.035 (corresponding to *T* = 19.8 h).

Additionally, our mathematical study in SI Section S.4 suggests that our 4D+X model can replicate the experimental data related to the transfection of KRAB and DNMT3A (Fig. 2) only within a parameter regime characterized by small *ε* and large *ζ*. The expressions defining these parameters are given in (SM.2). Therefore, we set 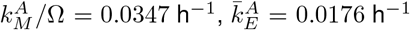, and 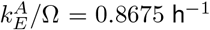, since these parameter values, along with the previously listed values of *δ* and D_tot_, allow us to be within the desired parameter regime and accurately replicate our experimental results.

Regarding the other parameters, we considered multiple values and selected those that allow us to accurately replicate all experimental results presented in this paper.

#### S.9.1 Parameter values used to realize the plots in Fig. 2I - top panel

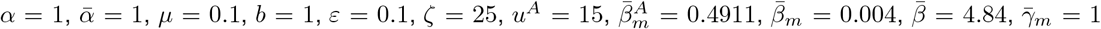, As initial conditions, we set 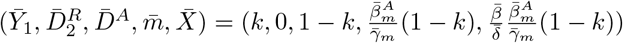, with 0 ≤ *k* ≤ 1.

#### S.9.2 Parameter values used to realize the plots in Fig. 2I - bottom panel

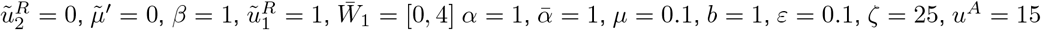. As initial conditions, we set 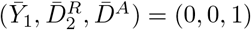.

#### S.9.3 Parameter values used to realize the plots in Fig. 2J

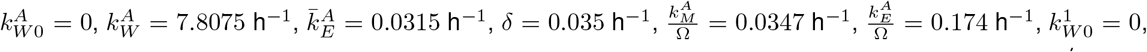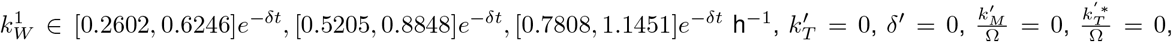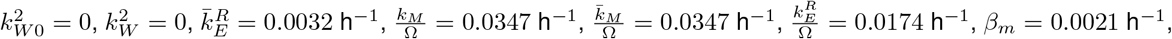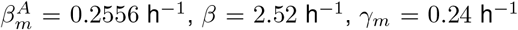. As initial condition, we consider 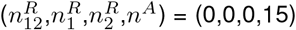 and the initial values of *n*^*X*^ of *n*^*m*^ at time *t* = 0 were set to their steady states of the ODEs.

#### S.9.4 Parameter values used to realize the plots in Fig. 2K

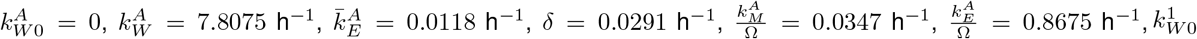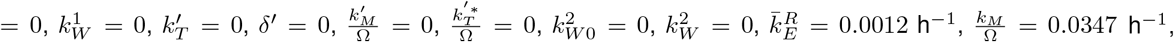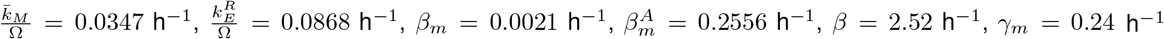. We consider four initial conditions: 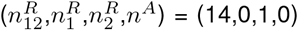 (blue), 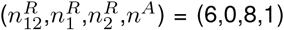 (red), 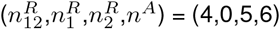 (yellow), 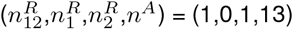 (purple). For each case, the initial values of *n*^*X*^ of *n*^*m*^ at time *t* = 0 were set to their steady states of the ODEs.

#### S.9.5 Parameter values used to realize the plots in Fig. 4H and J

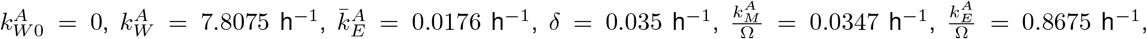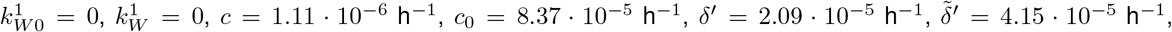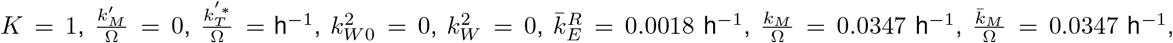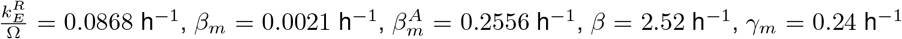. We consider eight initial conditions: 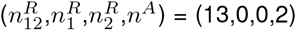(Bin 1), (12,0,0,3) (Bin 2), (8,0,0,7) (Bin 3), (6,0,0,9) (Bin 4), (5,0,0,10) (Bin 5), (4,0,0,11) (Bin 6), (1,0,0,14) (Bin 7), (0,0,0,15) (Bin 8). For each case, the initial values of *n*^*X*^ of *n*^*m*^ at time *t* = 0 were set to their steady states of the ODEs.

#### S.9.6 Parameter values used to realize the plots in SI Fig. SM.1B

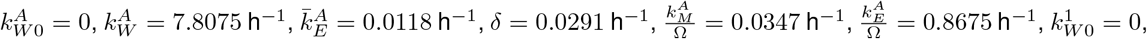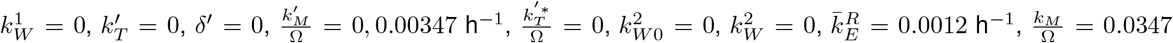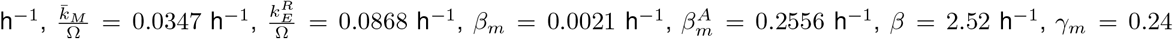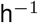. We consider four initial conditions: 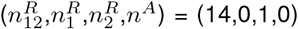 (blue), 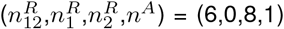 (red), 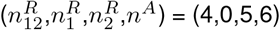 (yellow), 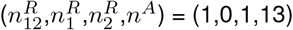(purple). For each case, the initial values of *n*^*X*^ of *n*^*m*^ at time *t* = 0 were set to their steady states of the ODEs.

#### S.9.7 Parameter values used to realize the plots in SI Fig. SM.1D - left hand-side panel

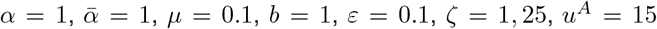. As initial conditions, we set 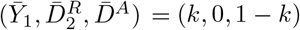, with 0 ≤ *k* ≤ 1.

#### S.9.8 Parameter values used to realize the plots in SI Fig. SM.1D - right hand-side panel

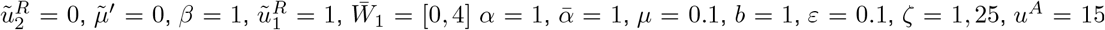. As initial conditions, we set 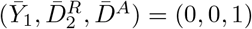.

#### S.9.9 Parameter values used to realize the plots in SI Fig. SM.1E

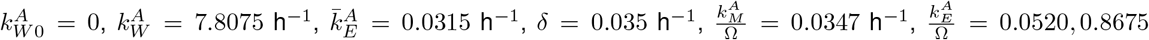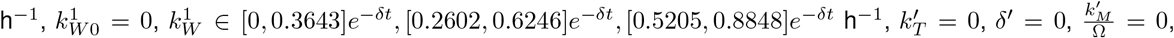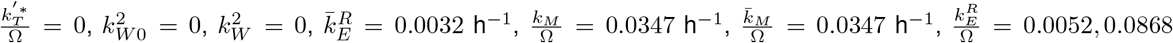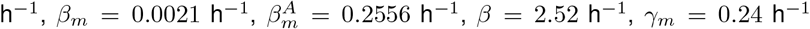. As initial condition, we consider 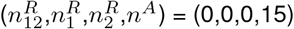 and the initial values of *n*^*X*^ of *n*^*m*^ at time *t* = 0 were set to their steady states of the ODEs.

#### S.9.10 Parameter values used to realize the plots in SI Fig. SM.2

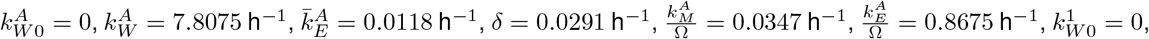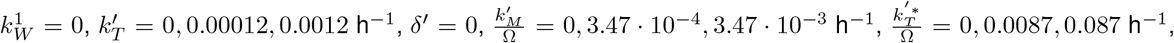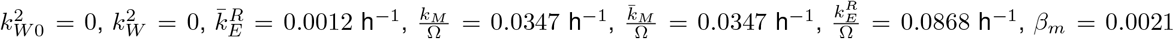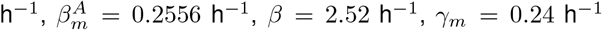. We consider four initial conditions: 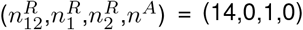 (blue), 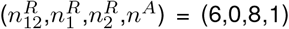 (red), 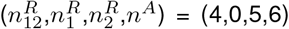 (yellow), 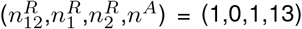 (purple). For each case, the initial values of *n*^*X*^ of *n*^*m*^ at time *t* = 0 were set to their steady states of the ODEs.

#### S.9.11 Parameter values used to realize the plots in SI Fig. SM.3A

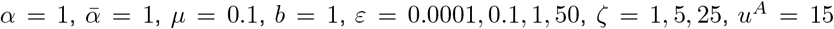. As initial conditions, we set 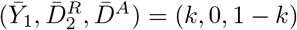,with 0 ≤ *k* ≤ 1.

#### S.9.12. Parameter values used to realize the plots in SI Fig. SM.3B

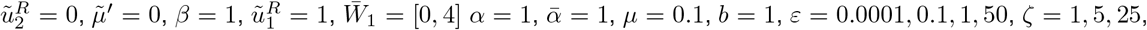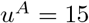. As initial conditions, we set 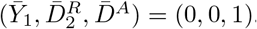.

#### S.9.13 Parameter values used to realize the plots in SI Fig. SM.3C

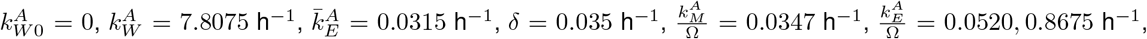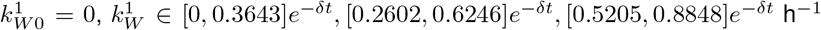 (left hand-side panel) and 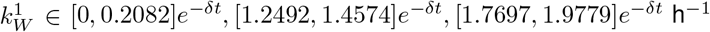 (right hand-side panel), 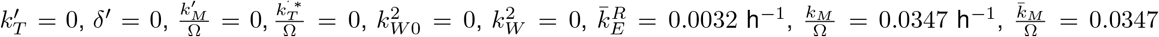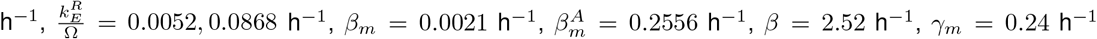. As initial condition, we consider 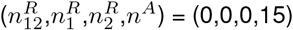 and the initial values of *n*^*X*^ of *n*^*m*^ at time *t* = 0 were set to their steady states of the ODEs.

#### S.9.14 Parameter values used to realize the plots in SI Fig. SM.3D

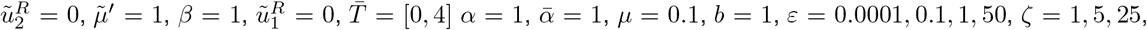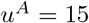. As initial conditions, we set 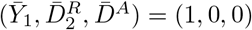.

#### S.9.15 Parameter values used to realize the plots in SI Fig. SM.3E

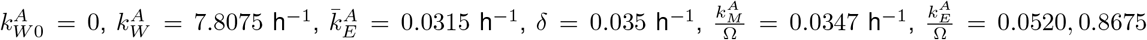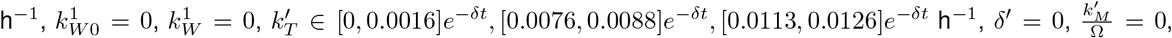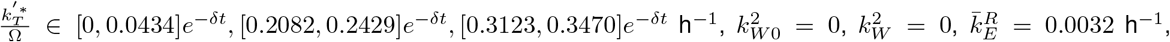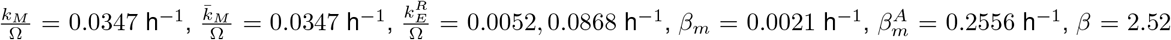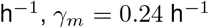. As initial condition, we consider 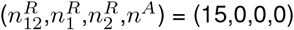 and the initial values of *n*^*X*^ of *n*^*m*^ at time *t* = 0 were set to their steady states of the ODEs.

#### S.9.16 Parameter values used to realize the plots in SI Fig. SM.4A

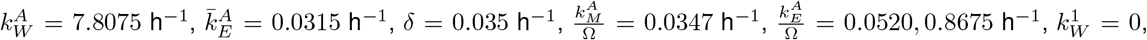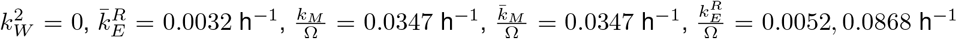. As initial condition, we consider 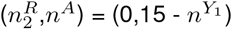.

#### S.9.17 Parameter values used to realize the plots in SI Fig. SM.4B

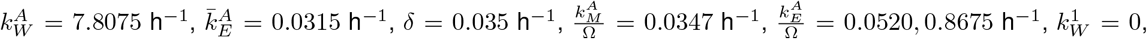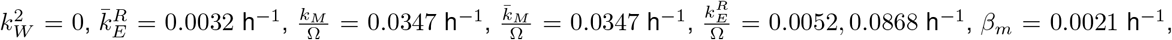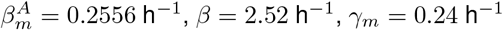. As initial condition, we consider 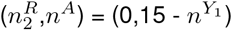 and the initial values of *n*^*X*^ of *n*^*m*^ at time *t* = 0 were set to their steady states of the ODEs.

#### S.9.18 Parameter values used to realize the plots in SI Fig. SM.5C

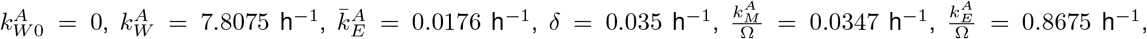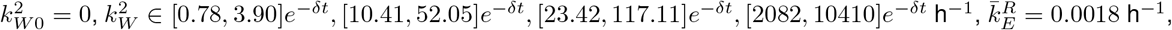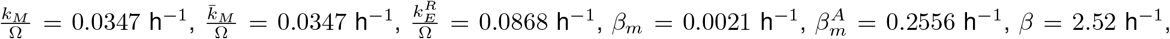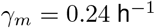. As initial condition, we consider 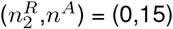 and the initial values of *n*^*X*^ of *n*^*m*^ at time *t* = 0 were set to their steady states of the ODEs.

#### S.9.19 Parameter values used to realize the plots in SI Fig. SM.5D

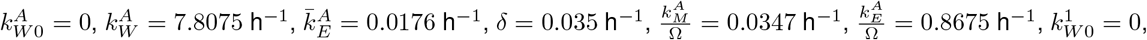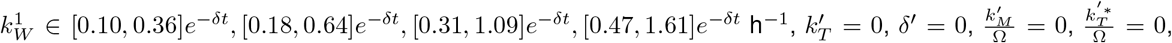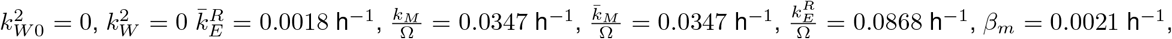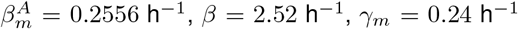. As initial condition, we consider 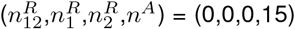 and the initial values of *n*^*X*^ of *n*^*m*^ at time *t* = 0 were set to their steady states of the ODEs.

#### S.9.20 Parameter values used to realize the plots in SI Fig. SM.6B

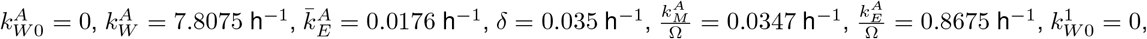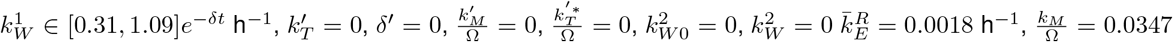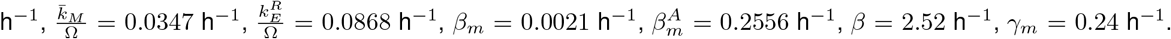. As initial condition, we consider 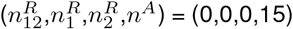 and the initial values of *n*^*X*^ of *n*^*m*^ at time *t* = 0 were set to their steady states of the ODEs. The distribution shown is after 30 days.

#### S.9.21 Parameter values used to realize the plots in SI Fig. SM.6C

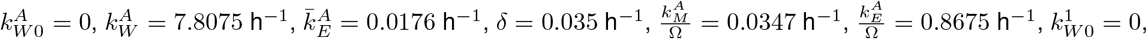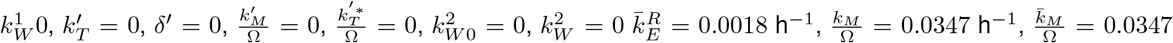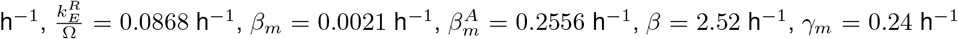. As initial condition, we consider values of 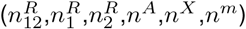 from the simulations shown in SI Fig. SM.6B and divide them with respect to their value of *n*^*X*^, following the following intervals: [0, 100] (red), [200, 800] (orange), [900, 10000] (blue).

#### S.9.22 Parameter values used to realize the plots in SI Fig. SM.6G

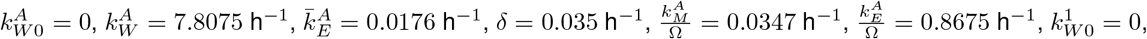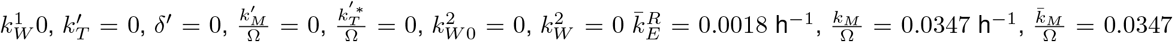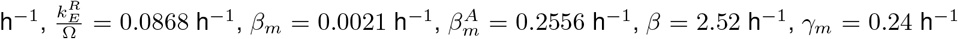. As initial condition, we consider values of 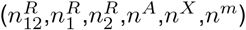 from the simulations associated with the orange distribution in SI Fig. SM.6D and divide them with respect to their value of *n*^*X*^, following the following intervals: [0, 80] (Bin 1), [90, 150] (Bin 2), [250, 400] Bin 3, [500, 10000] (Bin 4).

#### S.9.23 Parameter values used to realize the plots in SI Fig. SM.7A

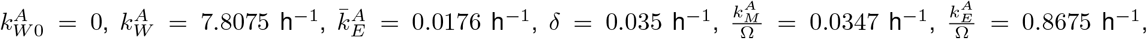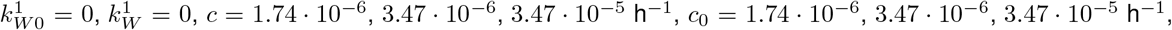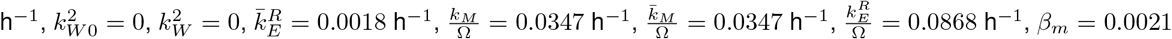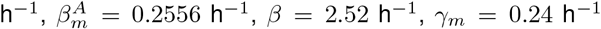. We consider four initial conditions: 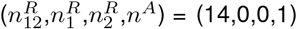 (blue), 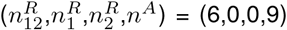 (red), 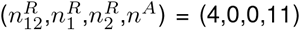 (yellow), 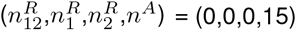 (purple). For each case, the initial values of *n*^*X*^ of *n*^*m*^ at time *t* = 0 were set to their steady states.

#### S.9.24 Parameter values used to realize the plots in SI Fig. SM.7B

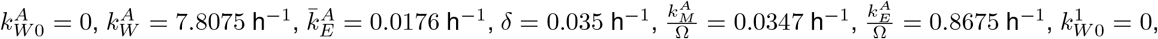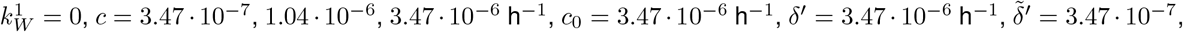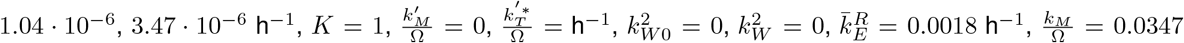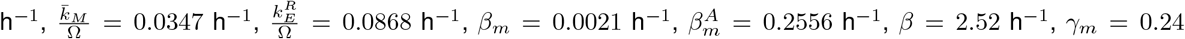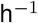. We consider four initial conditions: 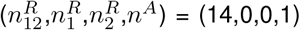 (blue), 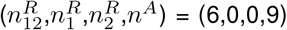 (red), 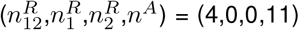 (yellow), 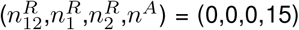 (purple). For each case, the initial values of *n*^*X*^ of *n*^*m*^ at time *t* = 0 were set to their steady states of the ODEs.

#### S.9.25 Parameter values used to realize the plots in SI Fig. SM.8A

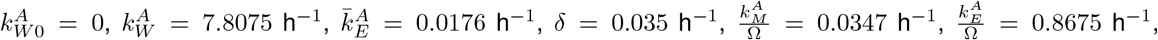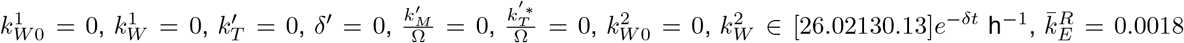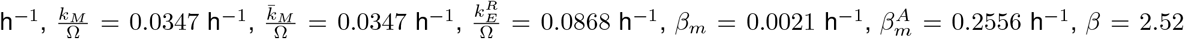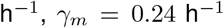. As initial condition, we consider values of 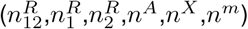 from the simulations associated with the orange distribution in SI Fig. SM.6C.

#### S.9.26 Parameter values used to realize the plots in SI Fig. SM.8B

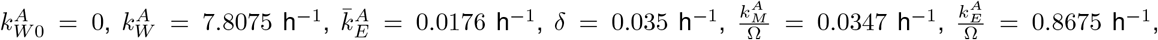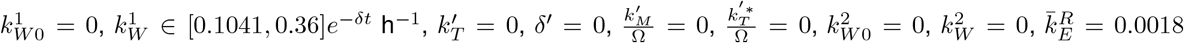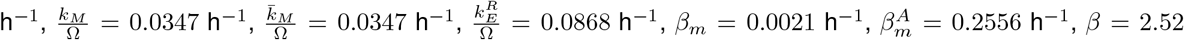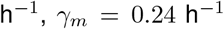. As initial condition, we consider values of 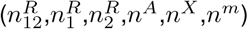 from the simulations associated with the orange distribution in SI Fig. SM.6C.

#### S.9.27 Parameter values used to realize the plots in SI Fig. SM.8C

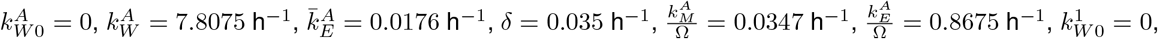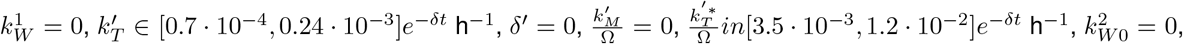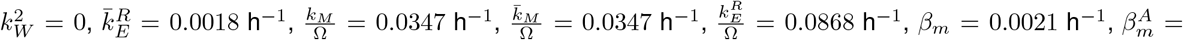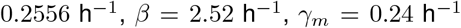. As initial condition, we consider values of 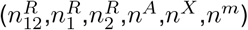 from the simulations associated with the orange distribution in SI Fig. SM.6C.

## S.10 Supplementary Experimental Figures

**Fig. ES.1.**
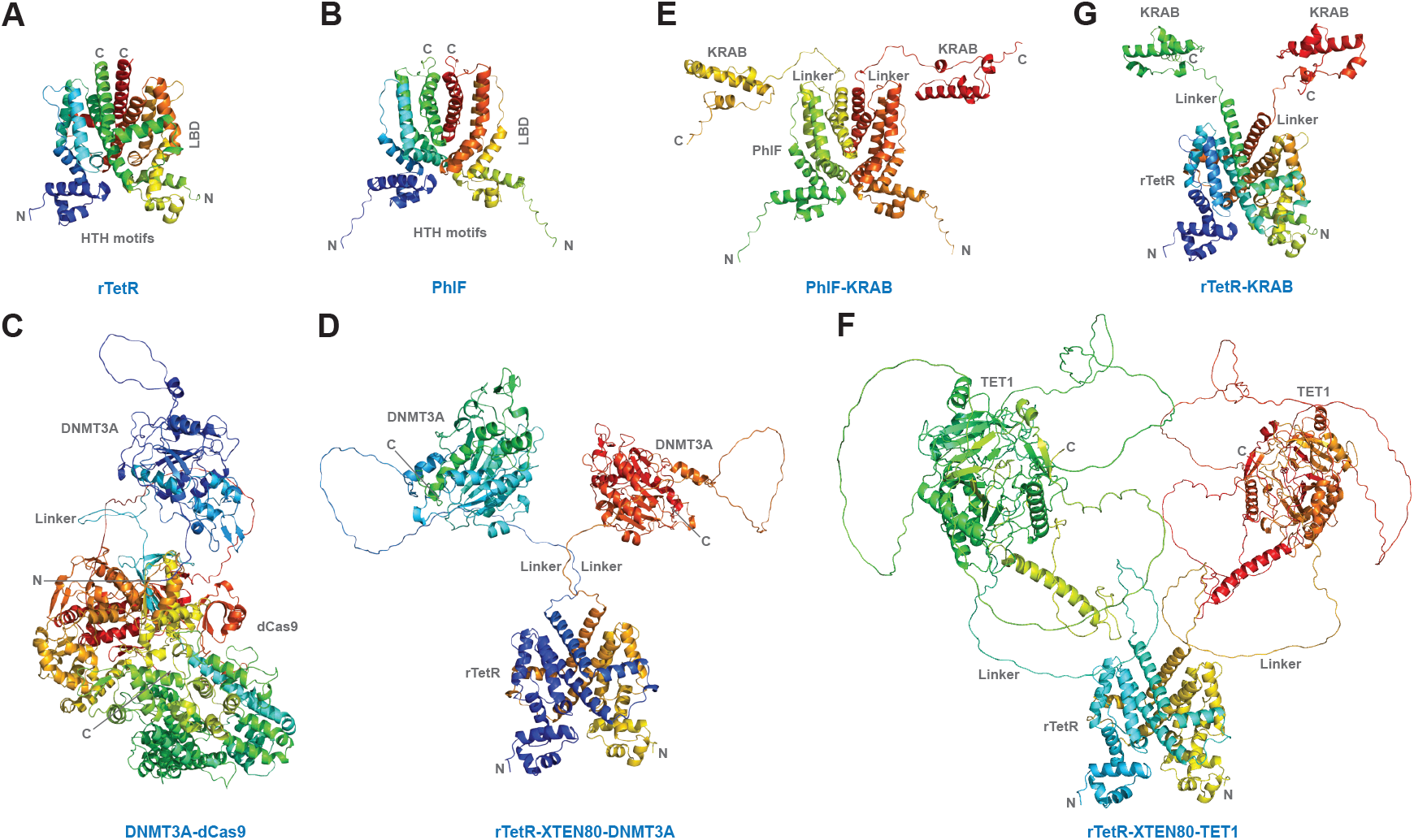
Protein structures of epigenetic regulators used in this study. **A** Homodimer structure of DNA-binding protein rTetR. **B** Homodimer structure DNA-binding protein PhlF. **C** DNMT3A, writer of DNA methylation, fused to programmable DNA-binding protein dCas9. **D** DNMT3A fused to DNA-binding protein rTetR. **E** KRAB, a regulator of H3K9me3, fused to DNA-binding protein PhlF. **F** TET1, an an eraser of DNA methylation, fused to rTetR. **G** KRAB fused to rTeR. Predictions performed using AlphaFold2 and PyMol (Methods). For rTetR and PhlF TetR-family proteins, the epigenetic effectors KRAB, DNMT3A, and TET1 are fused proximal to alpha helix 9 at the C-terminus using a flexible linker. Helix-turn-helix (HTH) motifs bind DNA. The ligand binding domain (LBD) binds dox in the case or rTetR and DAPG in the case of PhlF. For dCas9, DNMT3A is fused at the N terminus using a flexible linker.

**Fig. ES.2.**
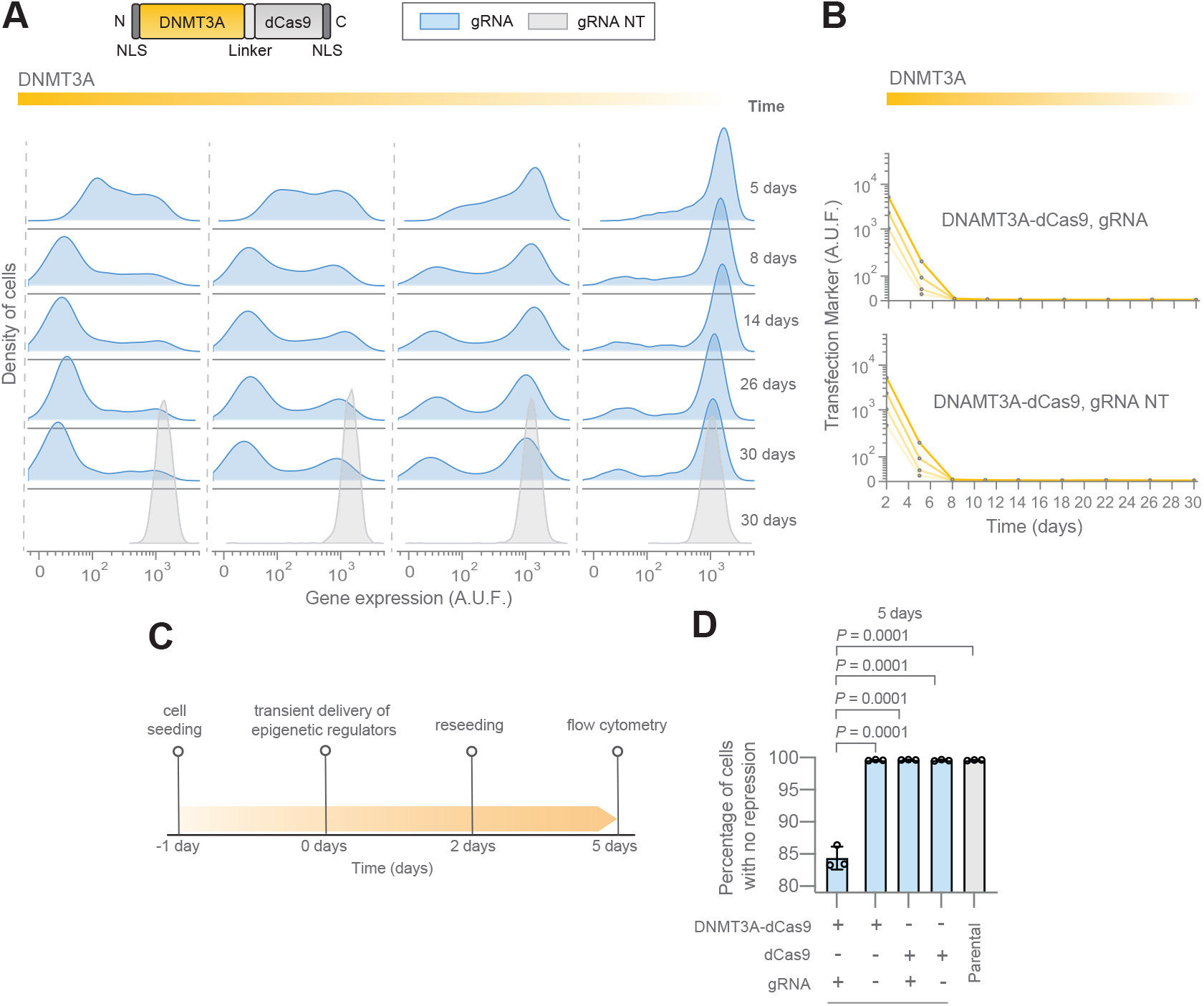
Stationary distributions of DNMT3A-silenced cells exhibit continuous gene expression. **A** Gene expression dynamics after epigenetic modulation using DNMT3A-dCas9 corresponding to the cells shown in Fig.2 D. Data shown are from a representative replicate from 3 independent replicates. DNMT3A-dCas9 was targeted to 5 target sites upstream of the promoter (gRNA). A gRNA with a scrambled target sequence (gRNA NT) was used as a control.**B** Transfection marker (EYFP) after epigenetic modulation using DNMT3A-dCas9 corresponding to the cells shown in Fig.2 D and Panel A. Data shown are from a representative replicate from 3 independent replicates. **C** Experimental overview of characterization shown in Panel D. **D** Experimental analysis of DNMT3A-dCas9 using various experimental conditions for repression of the reporter gene according to the experimental overview shown in Panel C. Cell densities are from a representative replicate from 3 independent replicates. Percentages and mean percentages are shown from three independent replicates (300 − 10^5^ gene expression A.U.F.). Statistical tests performed on all pairs and showing statistically significant results. Error bars are s.d. of mean percentage. \**P*<0.05, \*\**P*<0.01, \*\*\**P*<0.001, unpaired two-tailed t-test.

**Fig. ES.3.**
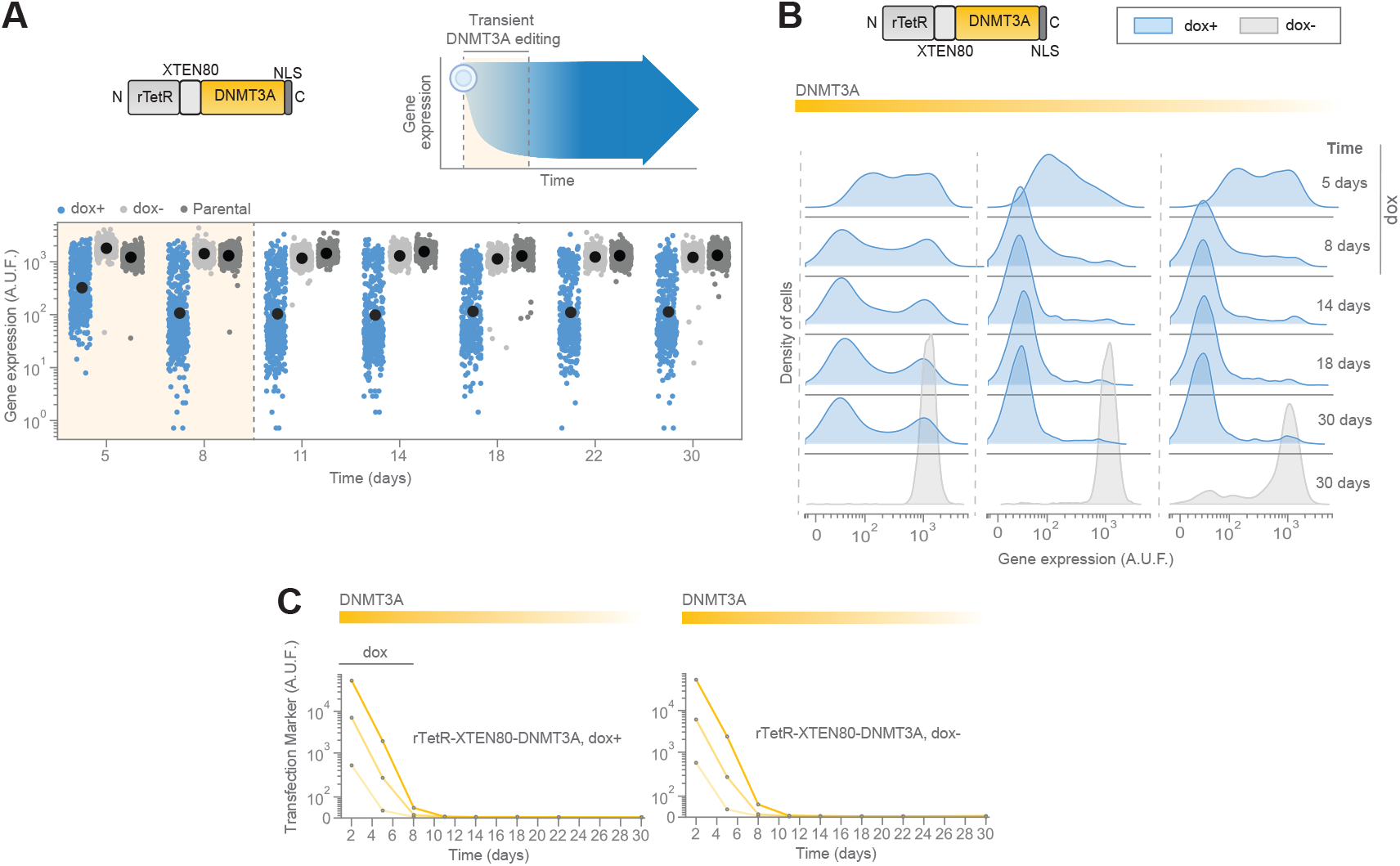
Stationary distributions of cells silenced using rTetR-XTEN80-DNMT3A exhibit intermediate levels of gene expression. **A** Single-cell gene expression measurements of cells edited with rTetR-XTEN80-DNMT3A. Shaded yellow corresponds to the time that doxycycline was applied while the transfection marker was also detected. Doxycycline was applied for 8 days. Data is from a representative replicate from 3 independent replicates. **B** Gene expression dynamics after DNMT3A-mediated editing with rTetR-XTEN80-DNMT3A correspoding to panel B for various trasfection levels. Shown are flow cytometry measurements of cells for three different levels of transfection that were sampled using fluorescence-activated cell sorting. **C** Transfection markers after epigenetic modulation using rTetR-XTEN80-DNMT3A corresponding to the cells shown in Panel A and B. Data shown are from a representative replicate from 3 independent replicates.

**Fig. ES.4.**
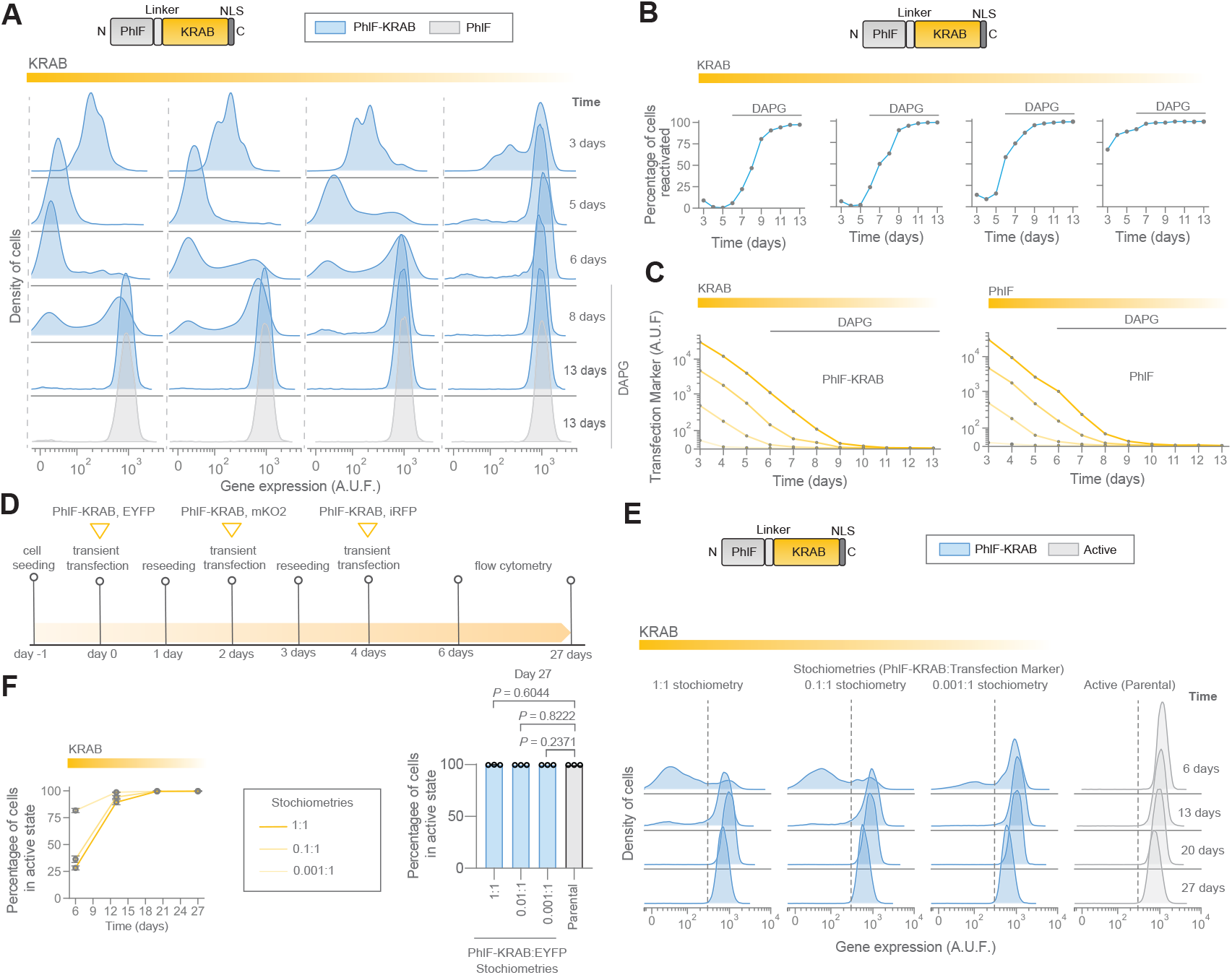
KRAB-mediated silencing does to confer memory. **A** Gene expression dynamics after PhlF-KRAB recruitment to the reporter gene. Shown are flow cytometry measurements of the reporter gene (EBFP2) for four different levels of transfection sorted using fluorescence-activated cell sorting. Data is from a representative replicate from 3 independent replicates. **B** Percentage of reactivated cells (400 − 10^5^ gene expression A.U.F.) corresponding to cell populations shown in panel A. **C** Transfection markers of epigenetic modulation using PhlF-KRAB corresponding to Fig. 2F,G and Panel A. Data shown are from a representative replicate of three independent replicates. **D** Experimental overview of KRAB-mediated silencing using PhlF-KRAB. **E** Density of cells for KRAB-mediated silencing using PhlF-KRAB according to Panel C. Data shown is from a representative replicate from 3 independent replicates. **F** Percentage of cell reactivation (300 − 10^5^ gene expression A.U.F.) in Panel D. Shown is the mean percentage from three independent replicates. Error bars are the s.d. of the mean. \**P*<0.05, \*\**P*<0.01, \*\*\**P*<0.001, unpaired two-tailed t-test.

**Fig. ES.5.**
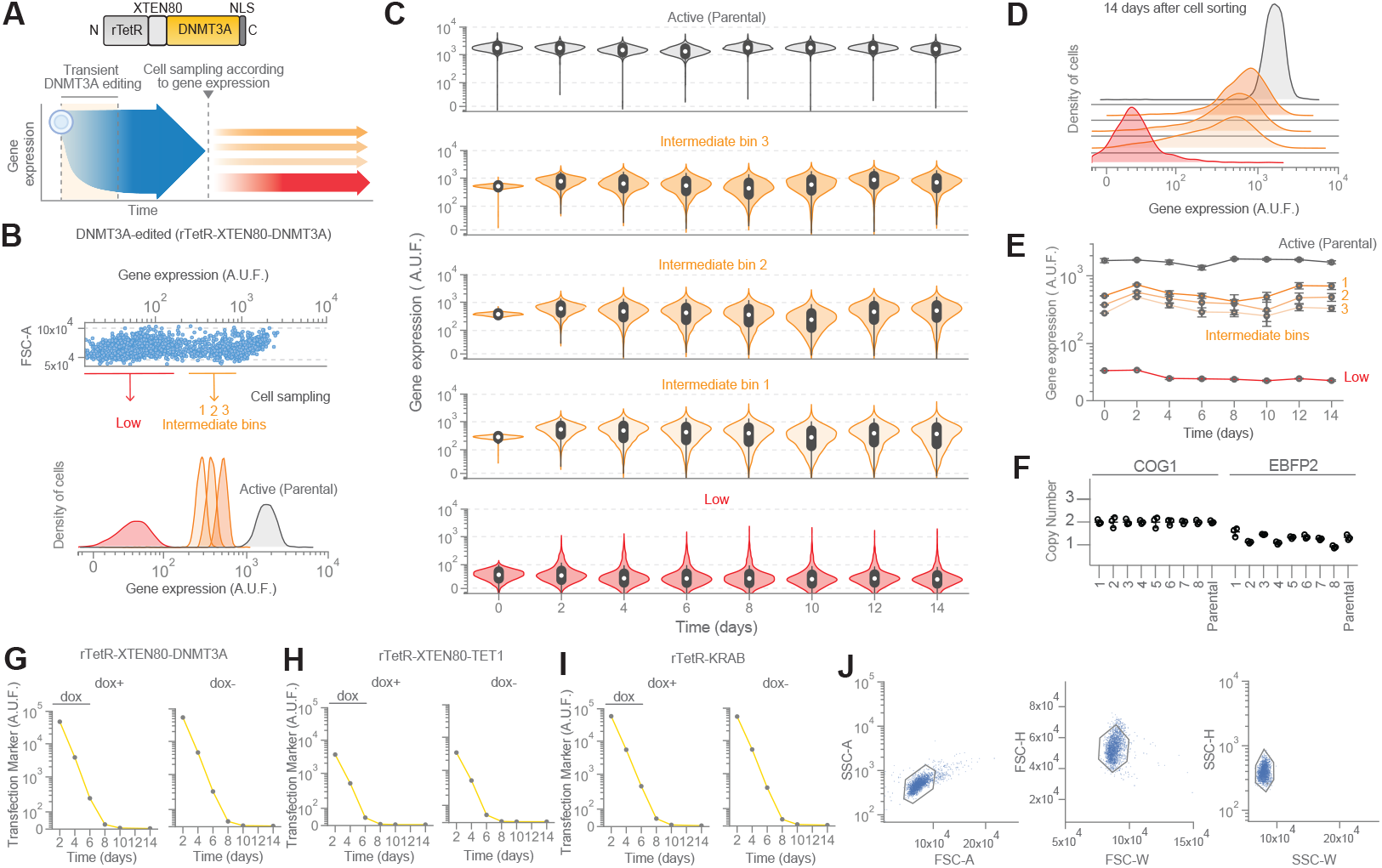
Intermediate levels of gene expression are stable in cell silenced using rTetR instead of dCas9 for targeted recruitment of DNMT3A. **A** Conceptual overview of experimental apprach. **AB** Fluorescence-activated cell sorting of cells exhibiting low, intermediate, and high levels of gene expression in DNMT3A-silenced cells (rTetR-XTEN80-DNMT3A). Data shown is from a representative replicate from three independent replicates. **C** Time-course flow cytometry analysis of the dynamics of gene expression exhibited by cells obtained in panel B. Data is from one representative replicate from three independent replicates. **D** Density of cells from flow cytometry measurements obtained on the last time point in panel C (14 days after cell sorting). Data shown is from one representative replicate from three independent replicates. **E** Dynamics of gene expression exhibited by cells shown in panels A-D. Shown is the mean of geometric mean fluorescence from three independent replicates. Error bars are s.d. of the mean. **F** Digital PCR corresponding to the parental cell line and monoclonal populations corresponding to Fig. 4 For each cell population, the average COG1 value was calculated (diploid). All data was then normalized to the COG1 average for each cell population separately. **G** Transfection markers corresponding to Fig. 5A. Data shown are from a representative replicate from 3 independent replicates. **H** Transfection markers corresponding to Fig. 5B. Data shown are from a representative replicate from 3 independent replicates. **I** Transfection markers corresponding to Fig. 5C. Data shown are from a representative replicate from 3 independent replicates. **J** Representative gating of cells used in this study. Data shown are from a representative replicate gating the cells bearing the reporter gene.

